# Hippocampal information topology breaks down in a mouse model of Alzheimer’s disease

**DOI:** 10.64898/2026.08.14.744888

**Authors:** Jess J. Yu, Hardik Rajpal, Mary Ann Go, Simon R. Schultz

## Abstract

Hippocampal spatial coding depends on coordination among neuronal assemblies, yet how network topology organises information processing across these assemblies, and how this is disrupted in disease, remain unknown. We apply Partial Information Decomposition to CA1 calcium imaging from young and aged wild-type and 5xFAD mice, quantifying redundant and synergistic information sharing within and between assemblies. In healthy CA1, between-assembly pairs carried more joint spatial information than within-assembly pairs, and this surplus was synergistic, establishing network topology as an organising principle of spatial coding. In aged 5xFAD CA1 this topological organisation broke down through two distinct routes: redundancy lost its topology dependence as modular assembly boundaries dissolved, and synergy lost context sensitivity during novel exploration, with the breakdown greatest where ageing and the 5xFAD genotype coincided. This functional decline was also accompanied by topological effects in the functional connectivity, where the genotype-age interaction resulted in reduced modularity, weighted clustering and small-worldness. Community-level emergence revealed a complementary cross-scale shift toward higher-order integration during ageing, which was reversed by the genotype-age interaction. We isolate the compounding effect of ageing in Alzheimer’s disease as the driver of disruption in information processing and functional connectivity across neuronal assemblies in the mouse hippocampus.

## Introduction

Individual hippocampal neurons encode an animal’s position in an environment^1,2^, and at the population level these place codes form ensemble representations that can be read out from simultaneously recorded cells^3,4^, even from short windows of population activity^5^, and from two-photon calcium imaging of large CA1 populations^6,7^. The information transmitted into CA1 across the Schaffer collaterals, and the limits on what a recurrent population can convey, have been characterised analytically^8–10^. Large-scale population recordings have revealed that the hippocampus generates flexible representations of the spatial environment by recruiting neurons into coordinated assemblies^11–13^ that tile a high-dimensional activity space^14,15^. This modular organisation, a defining feature of cortical and subcortical circuits more broadly^16,17^, is thought to support two competing computational demands: robust signal transmission within assemblies, which ensures reliable readout, and flexible integration across assemblies, which enables the formation of new, non-interfering spatial representations for novel contexts^18–20^. Both of these computational capacities are challenged by neurodegeneration under ageing and Alzheimer’s disease.

Ageing and Alzheimer’s disease (AD)-related amyloid pathology are each associated with degraded place-cell representations and impaired spatial memory^21^. Prior studies have documented degraded entorhinal grid-cell tuning, alongside a largely preserved hippocampal place code, in transgenic AD mouse models^22^. Degraded place-cell coding has been reported in the Tg2576^23^ and 5xFAD^24^ lines, and in 3xTg mice, which additionally carry tau pathology^25^; remapping is impaired in an *App* knock-in model^26^ and in 5xFAD^27^; and clusters of hyperactive neurons arise near amyloid plaques^28^. Normal ageing alone alters hippocampal place coding in a subregion-specific way^29,30^. Such observations, however, describe symptoms at the level of individual neurons or bulk population statistics. They do not resolve whether the circuit-level deficit is a uniform degradation of neural coding or a specific disruption of the network architecture that determines *how* neurons share spatial information. In particular, it remains unknown whether ageing and 5xFAD genotype act on the same circuit mechanism, on independent mechanisms, or whether their combination produces a qualitatively distinct failure that neither factor causes alone.

Information theory has increasingly offered a natural framework to characterise computational differences between neural circuits, and has been applied directly to neuronal recordings and spiking network models to quantify how populations of neurons jointly encode information^31–34^. These decompositions have exposed circuit-level properties that are invisible to conventional pairwise analyses, including synergistic subsystems and higher-order interactions that no individual neuron or correlation reveals^35–38^. This power rests on the distinction between redundant and synergistic population codes. Correlated firing between neurons can render a population code redundant, so that either neuron reports much of what the other does, or synergistic, so that the pair jointly specifies more than the sum of its parts^39–43^. Earlier work decomposed the information carried by a population into rate, correlational and temporal components^44,45^; Partial Information Decomposition extends that programme by separating the redundant and synergistic contributions explicitly^46,47^. Redundancy in particular has been treated as a design variable of neural codes since the efficient coding hypothesis^48,49^. Across cortical systems, redundant interactions concentrate in structurally coupled, modular circuits and confer robustness, whereas synergistic interactions predominate in the higher-order networks that support integrative computation^36^. In the hippocampus, the coactivity structure of CA1 is itself dynamic, reorganising from robust, rigid ensembles toward more flexible configurations as memories are stabilised and updated^50^. How these two modes of information processing are organised by network topology, and how this organisation is disrupted in disease, have not been directly tested using information-theoretic decomposition.

To address these questions, we combined Partial Information Decomposition (PID)^38,46,51^ with multiscale community detection^52,53^ to dissect topology-dependent information sharing in CA1 ensembles from both young and aged, wild-type (WT) and 5xFAD mice. PID decomposes the joint spatial information carried by neuron pairs into redundant, unique, and synergistic components, enabling us to quantify whether the information shared by a pair is robust and duplicated (redundancy) or emerges only from their joint activity (synergy). By intersecting these information-theoretic measures with the community structure of the functional network, we tested whether the *topological identity* of a neuron pair (whether it lies within or across assembly boundaries) determines its information-processing strategy. This cellular-resolution approach extends recent information-decomposition studies of AD, which reveal a global shift from synergy toward redundancy at the macroscale of whole-brain human functional imaging^54^ but cannot localise the deficit to specific circuit elements. By resolving the balance of redundancy and synergy at the level of individual neuron pairs and their assembly membership (Fig. 1), we investigate not only whether information sharing degrades but where in the network architecture it does so.

**Figure 1.**
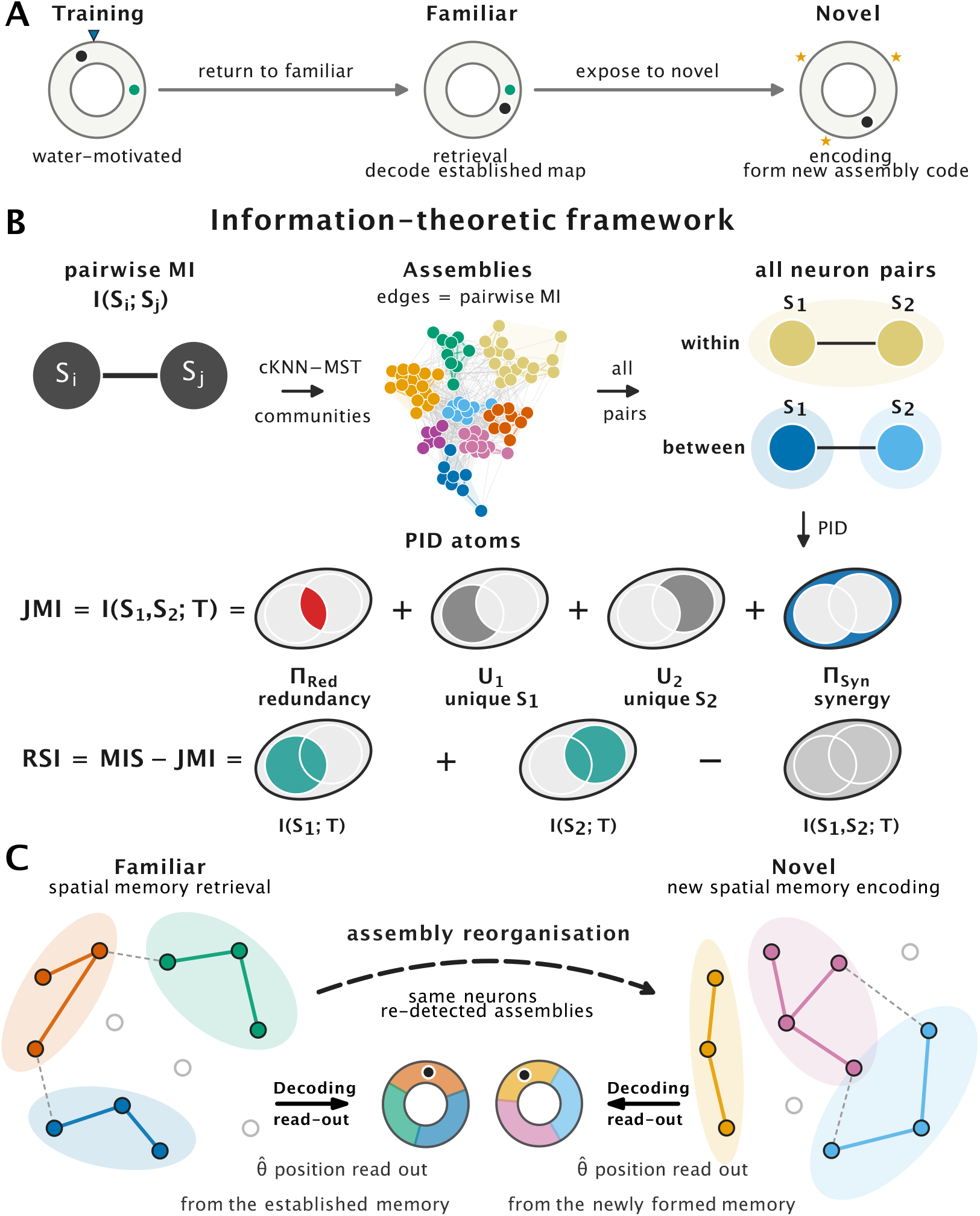
Experimental paradigm, information-theoretic framework and assembly reorganisation between spatial-memory retrieval and encoding. Dataset and acquisition follow Go et al.^21^, framed by the “activity-space” view of assembly coding^11,14^. **A**, Paradigm. Mice ran laps on a circular track for water reward, imaged across three stages: a *training* environment (habituation, track fam1), the *familiar* environment (retrieval), and a *novel* environment (same track, new cues; encoding a new map). Orange stars, novel cues; other markers, reward location and a fixed landmark. Cohort and imaging details in Methods. **B**, Information-theoretic framework (example session, case 12, 86 neurons). Pairwise NSB mutual information *I*(*S*_*i*_; *S* _*j*_) defines a cKNN–MST functional network, partitioned into assemblies by multiscale Markov stability; pairs are then within- or between-assembly. Each pair with position *T* enters the Partial Information Decomposition, splitting *I*(*S*_1_, *S*_2_; *T*) into redundancy (Π_Red_, red), unique components (*U*_1_, *U*_2_) and synergy (Π_Syn_, blue) under the Minimum Mutual Information lattice, summarised by the Redundancy–Synergy Index (RSI *>* 0 redundancy-dominated, RSI *<* 0 synergy-dominated; Methods). **C**, Assembly reorganisation. *Left* (familiar, retrieval): activity partitions into location-tuned assemblies tiling the track, so position 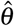 is decoded from the established community code. *Right* (novel, encoding): the novel context drives a new assembly code, re-detected *independently per session*, so membership, identity and colour are *not* matched across environments. Dots are CA1 neurons (filled/coloured = assembly-assigned, open = silent); coloured hulls are assemblies; edges are significant mutual-information connections; the lower ring is track position, with wedges where each assembly is tuned. **4/60**

We report four principal findings. First, in healthy CA1, network topology organises spatial coding: between-assembly pairs carry more joint spatial information than within-assembly pairs, and this surplus is synergistic (Fig. 2). Second, this topological organisation breaks down through two distinct routes: redundancy losing its topology dependence as assembly boundaries dissolve, and synergy losing context sensitivity. This breakdown is greatest in aged 5xFAD CA1, where ageing and the 5xFAD genotype coincide (Figs. 3, 4). Third, this functional breakdown has a structural correlate: backbone modularity, measured as the excess over a strength-preserving rewired null, shows a genotype × age deficit 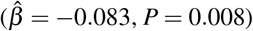, with weighted clustering and small-world organisation shifting in the same direction, so network structure and function decline in parallel in aged 5xFAD CA1 (Fig. 5). Fourth, community-level emergence reveals a complementary cross-scale shift: ageing moves CA1 toward higher-order synergistic integration, which the genotype × age interaction reverses toward redundancy (Fig. 6). Across these findings, it is the interaction between ageing and the 5xFAD genotype that disrupts the circuit-level structure of information processing in CA1.

**Figure 2.**
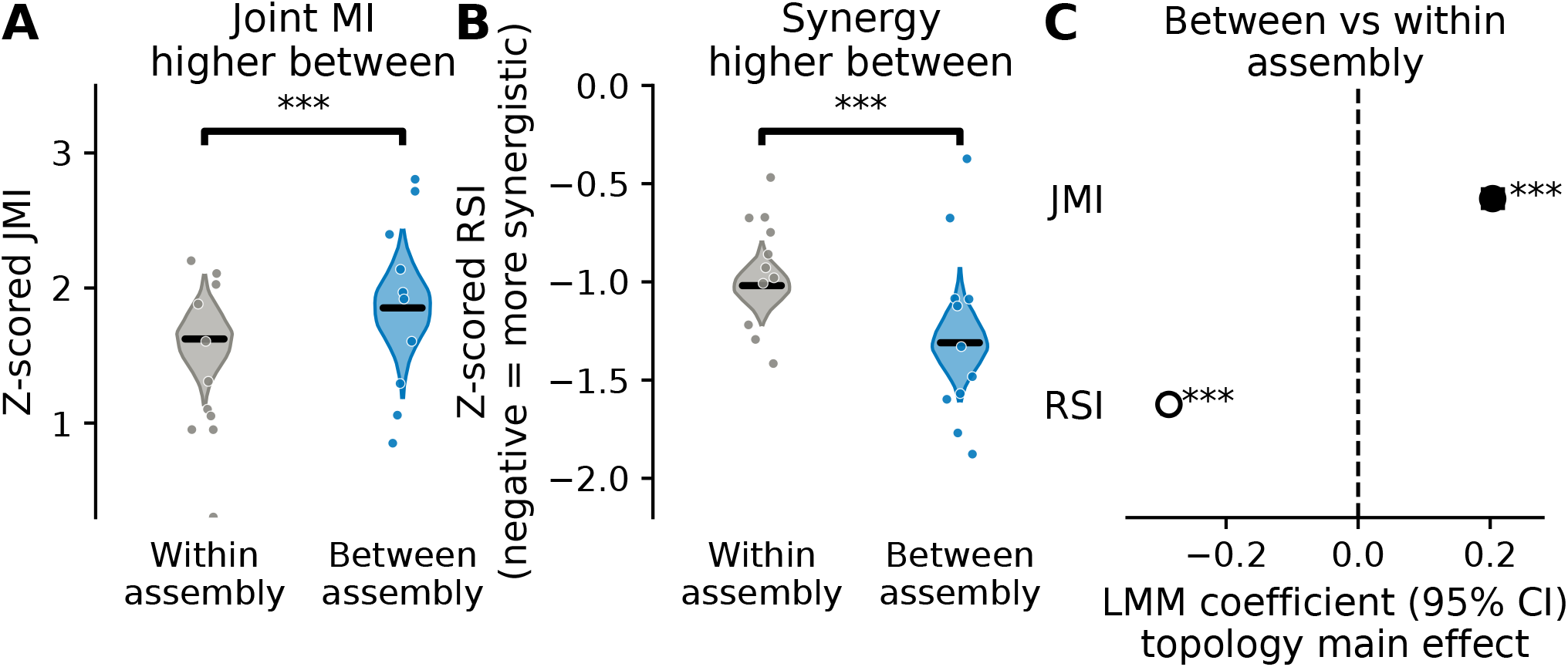
Network topology organises spatial information processing in healthy CA1. **A**, Z-scored joint mutual information (JMI) for within-assembly and between-assembly neuron pairs in young wild-type (WTY) animals during familiar-environment exploration. Between-assembly pairs carry more joint spatial information; the significance bracket shows the LMM topology main effect (pair type; * * *; cf. panel C). **B**, Z-scored Redundancy–Synergy Index (RSI) for the same comparison. Between-assembly pairs are more synergy-biased (more negative RSI; bracket, LMM topology main effect, * * *; cf. panel C). Negative RSI indicates synergy-dominated coding; positive RSI indicates redundancy-dominated coding. **C**, LMM coefficient estimates (95% CI) for the topology main effect on JMI 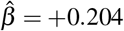 and RSI 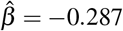, *P <* 10^−100^), confirming topology as the dominant fixed effect in a full-factorial model simultaneously accounting for age, genotype, and environment. Violins, hierarchical bootstrap (*B* = 10,000); dots, individual animal–session medians; stars and brackets, LMM fixed-effect test (Methods). WTY *n* = 11 familiar-environment sessions (A,B); full-factorial LMM *N* = 77 sessions (C). * * *P <* 0.01, * * \**P <* 0.001.

**Figure 3.**
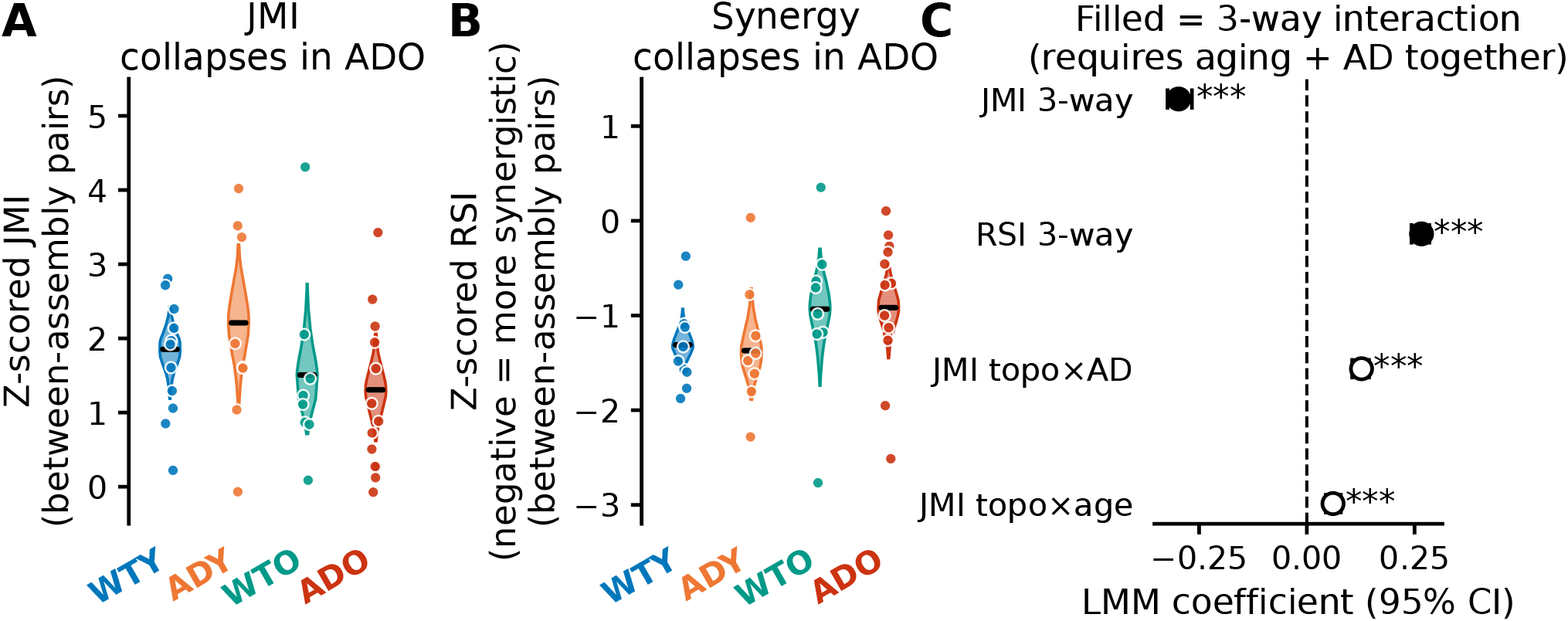
The topology-dependent synergy advantage collapses only when ageing and amyloid pathology coincide. **A**, Z-scored JMI for between-assembly pairs across all four experimental groups during familiar-environment exploration. Distributions: hierarchical-bootstrap grand-mean violins (*B* = 10,000); dots show per-case (animal × environment) medians. The group medians fall in the order ADY *>* WTY *>* WTO *>* ADO (ADO smallest), but no pairwise WTY-vs-other contrast reaches significance under the two-sided hierarchical bootstrap at the present per-group sample sizes (all *P* ≥ 0.15). Primary statistical inference is provided by the LMM three-way interaction in panel C. **B**, Z-scored RSI for between-assembly pairs. WTY and ADY animals show the most synergy-biased (most negative) values; WTO and ADO are shifted toward zero, indicating reduced synergy bias and a partial shift toward redundancy-dominated coding. Pairwise contrasts against WTY are again non-significant under the hierarchical bootstrap at this *n* (all *P* ≥ 0.10); the ageing-driven RSI shift is established by the LMM Age main effect and Topology × genotype × age interaction (see text). **C**, LMM coefficient estimates (95% CI) for the three-way interaction (Topology × genotype × age) for JMI 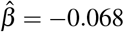 and RSI 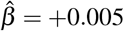, shown alongside the constituent two-way interactions (open markers). The three-way terms are the largest interaction coefficients in the model, indexing the topology-dependent collapse in aged 5xFAD CA1. *N* = 40 familiar-environment sessions (WTY *n* = 11, ADY *n* = 8, WTO *n* = 8, ADO *n* = 13; A,B); full-factorial LMM *N* = 77 sessions (C).

**Figure 4.**
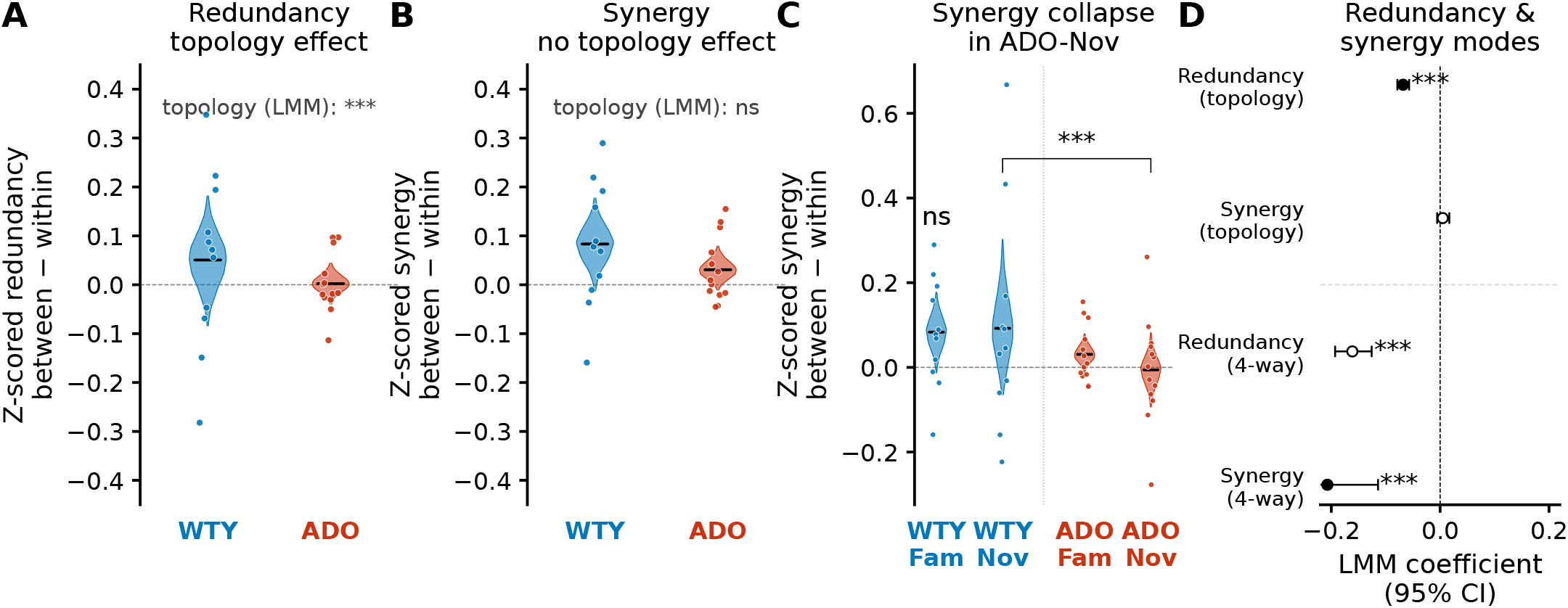
Redundancy and synergy are disrupted through mechanistically distinct pathways. **A**, Topology-dependent redundancy, expressed as the per-case contrast of medians, (between − within) (*z*-scored), for WTY and ADO animals in the familiar environment. Values below the dashed zero line indicate that within-assembly pairs carry more redundant spatial information than between-assembly pairs. WTY shows a negative contrast (LMM topology main effect 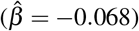, *P* = 8.3 × 10^−30^); the contrast is attenuated in ADO, consistent with the dissolution of modular structure. **B**, Topology-dependent synergy contrast of medians, (between − within) (*z*-scored), for the same groups and environment. Values above the dashed zero line would indicate that between-assembly pairs carry more synergistic spatial information than within-assembly pairs. The contrast sits near the zero line in both WTY and ADO (LMM topology main effect 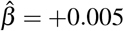, *P* = 0.43; ns): synergy carries no topology main effect. **C**, Topology-dependent synergy contrast (between − within) (*z*-scored) per case across the four group × environment conditions (WTY familiar, WTY novel, ADO familiar, ADO novel). The contrast is preserved at or above zero in WTY in both environments and in ADO-familiar, and collapses below zero selectively in ADO-novel, where between-assembly synergy falls below within-assembly synergy. The bracket shows the LMM synergy four-way topology × environment × genotype × age interaction (* * *). This selective collapse reflects the loss of on-demand cross-assembly integration during novel-environment exploration. **D**, LMM coefficient estimates (95% CI) summarising the mechanistic asymmetry: redundancy carries a significant topology main effect 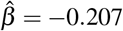 while synergy does not 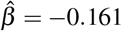, overlapping zero). Both atoms additionally carry a significant four-way topology × environment × genotype × age interaction (synergy 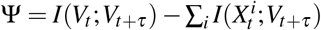, *P* = 1.3 × 10^−6^; redundancy 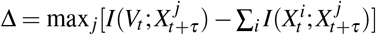, *P* = 6.8 × 10^−21^); the distinguishing feature is therefore the baseline topology main effect, present for redundancy and absent for synergy. Filled markers denote these two distinguishing terms; open markers the complementary terms. In **A**–**C**, violins and CIs are the hierarchical bootstrap (*B* = 10,000) of the per-case topology contrast, with overlaid dots the per-case (animal × environment) contrasts; stars and brackets (**A**–**D**) are LMM fixed-effect tests (Methods). Full within- vs. between-assembly distributions are shown in Supplementary Fig. 6. WTY *n* = 11 and ADO *n* = 13 familiar-environment sessions (A,B); WTY 11*/*11 and ADO 13*/*13 familiar/novel sessions (C); full-factorial LMM *N* = 77 sessions (D). ns, not significant; \**P <* 0.05, * * *P <* 0.01, * * \**P <* 0.001.

**Figure 5.**
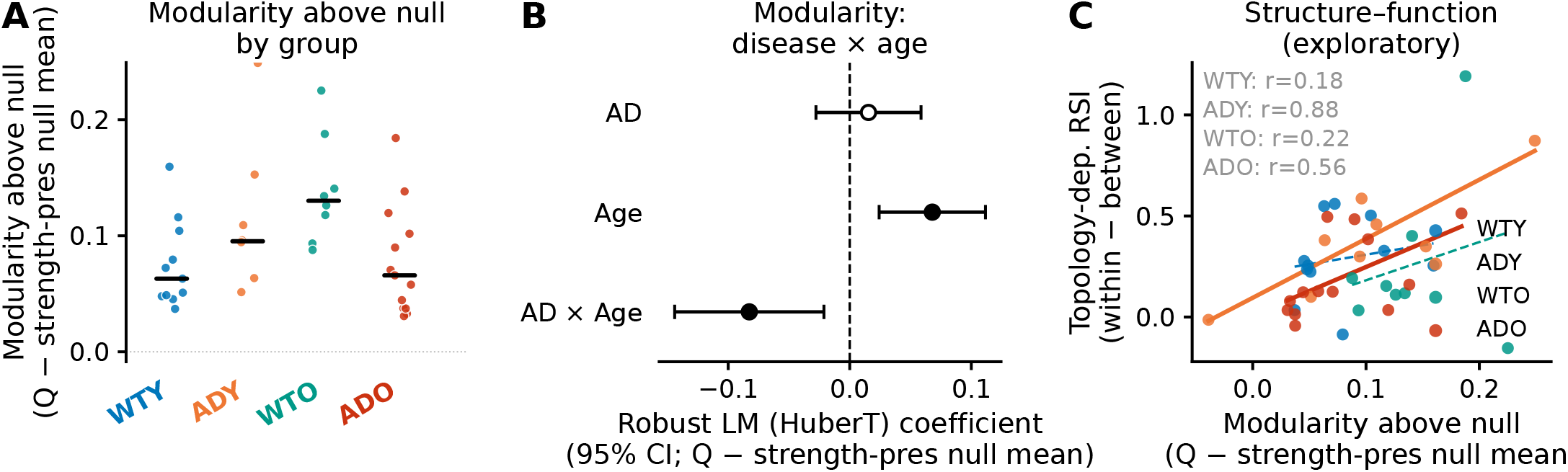
A genotype × age modularity deficit in aged 5xFAD CA1. **A**, Per-session modularity above the strength-preserving null (Δ*Q*^str^ = *Q*_obs_ −⟨*Q*_null_⟩; Markov-stability partition, familiar environment) for all four groups (WTY blue, ADY orange, WTO teal, ADO red; dots individual sessions, black bars group medians). Ageing raises modularity above the strength-preserving null, but the aged-5xFAD combination (ADO) is no higher than either single factor alone (ADY, WTO), the signature of the negative genotype × age interaction quantified in **B. B**, Robust linear-model coefficients (Huber M-estimator, 95% confidence intervals) for Δ*Q*^str^, showing the genotype (AD), age, and genotype × age (AD × age) terms of the full-factorial model (only these three drawn; full specification in Methods). Markers are monochrome black; significance is shown by marker *fill* (filled, *P <* 0.05; open, non-significant) and the adjacent star level marks the *P* threshold. The age main effect (*P* = 0.002) and the negative genotype × age interaction (*P* = 0.008) are significant while the genotype main effect is not (*P* = 0.48); the interaction is the aged-5xFAD modularity deficit. The full eight-metric coefficient forest (including the non-significant novelty terms) and the per-metric null-corrected strips are in Supplementary Fig. 11 (panels A–E). **C**, Exploratory structure–function link: per-session Δ*Q*^str^ versus the topology-dependent RSI (within − between) in the familiar environment, with per-group regression lines (solid where *P <* 0.05, dashed otherwise) and Pearson *r*. Larger values on the *x* axis mean within-assembly pairs are more redundancy-biased relative to between-assembly pairs. Both 5xFAD groups reach significance (ADY: *r* = 0.88, *n* = 8; ADO: *r* = 0.56, *n* = 13); neither wild-type group does. Stars in **B**: ^*^*P <* 0.05, ^**^*P <* 0.01, ^***^*P <* 0.001. *N* = 40 familiar-environment sessions (**A, C**); full Huber-model model *N* = 77 sessions (**B**).

**Figure 6.**
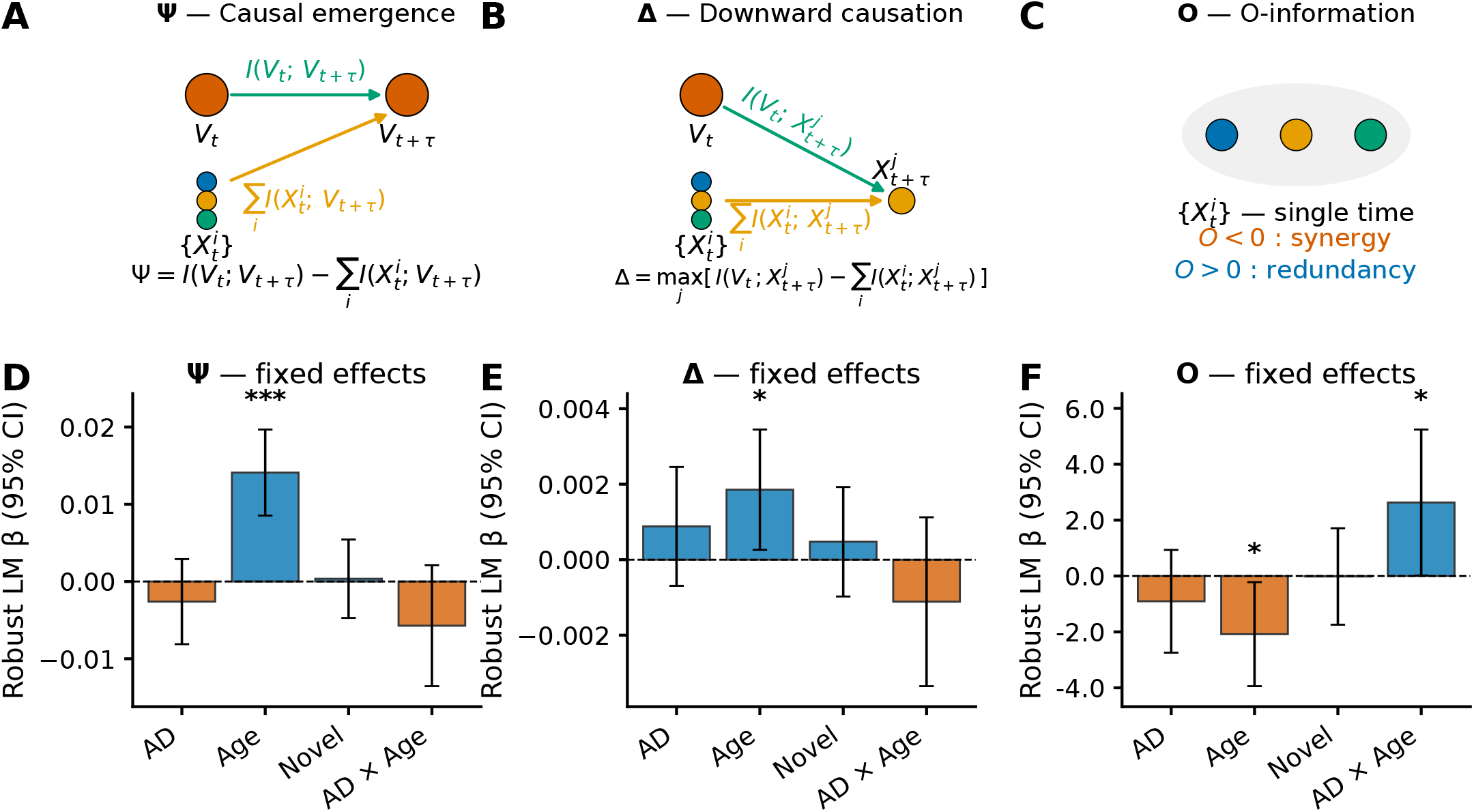
Multivariate community-level emergence: ageing shifts information toward higher-order synergy, which amyloid reverses to redundancy in aged animals. **A–C**, Schematics of the three multivariate emergence measures computed on the assembly community time-series, one measure per panel, with each operation labelled directly on the design. **A**, Causal emergence 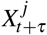 (macro auto-prediction beyond the sum of parts). **B**, Downward causation 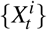 (macro state predicts a part beyond what other parts predict; the target 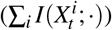 is drawn on the micro row, so the macro-term arrow points downward). **C**, Single-time O-information among parts (*O <* 0 synergy, *O >* 0 redundancy). Definitions follow Rosas et al.^57^ as applied to neuronal-population emergence^58^. Across A and B, *V* is the macroscopic (community) state and 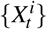 the individual micro parts; the green “macro term” arrow is the predictive information carried by *V* (*I*(*V*_*t*_;)) and the orange “parts term” arrow the corresponding sum over parts 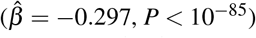, so each measure is macro parts. Panel C is the single-time O-information among the micro parts 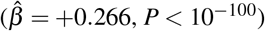 alone and has no macro term; its three coloured discs only distinguish nodes. **D–F**, Robust linear-model (Huber M-estimator) fixed-effect coefficients with 95% confidence intervals for Ψ (D), Δ (E) and O-information (F); each panel plots the AD, Age and Novel main effects and the AD × Age interaction. The model additionally fits the remaining two- and three-way interactions among Genotype (5xFAD), Age (Old) and Novel (novel environment) and a mean-centred community-count covariate (not shown). Stars: ^*^*P <* 0.05, ^**^*P <* 0.01, ^***^*P <* 0.001. Ageing alone increases Ψ and Δ and decreases O-information (toward synergy); the genotype × age interaction reverses O-information toward redundancy, paralleling the pair-level synergy collapse in Fig. 4. Per-delay traces and per-group integrated surrogate-corrected score boxplots (Familiar vs Novel; for Ψ/Δ: sum of [observed − surrogate median] over delays; for O-information: standardised *z*-score against surrogate *σ*) are provided in Supplementary Fig. 13. *N* = 77 sessions across 4 groups (Ψ/Δ); O-information *N* = 75 (two undefined-score sessions excluded).

## Results

### Experimental approach and dataset

We recorded CA1 pyramidal-cell ensembles via two-photon calcium imaging during circular track navigation in four experimental groups: young wild-type (WTY, *n* = 11 familiar-environment sessions), young 5xFAD (ADY, *n* = 8), aged wild-type (WTO, *n* = 8), and aged 5xFAD (ADO, *n* = 13), each explored in both familiar and novel environments (Fig. 1). The calcium traces (jGCaMP7s/GCaMP6s) were deconvolved to estimate spike amplitudes, which were globally discretised into four equaloccupancy amplitude bins plus a silent (zero-amplitude) state (five states total, *k*_neu_ = 5) across all 40 familiar-environment sessions, ensuring comparable entropy scales across animals, groups, and environments. Spatial position along the circular track was binned into 50 equal-width spatial bins. For each session, a significance-constrained functional connectivity backbone was constructed by intersecting topology-adaptive Continuous k-Nearest Neighbours (CkNN) edges with a Minimum Spanning Tree (MST) overlay, retaining only statistically significant mutual information edges (200 circular-shift surrogates per pair; *P <* 0.05; Fig. 1B). Functional assemblies were identified within this backbone by scanning the Markov stability function across resolution scales using PyGenStability and selecting the most robust partition (Fig. 1B; see Methods). Example assembly-level spatial tuning curves on the circular track (high- vs low-MI assemblies in the same session), a single-cell place-field example and the underlying MI / JMI values are shown in Supplementary Fig. 2. All downstream PID analyses classify neuron pairs as within-assembly (both neurons in the same community) or between-assembly (neurons in different communities) based on this partition.

### Network topology organises spatial information processing in healthy CA1

In young wild-type (WTY) animals exploring the familiar environment, the topological identity of a neuron pair determined both how much spatial information it carried and how that information was shared. Pairs spanning *different* functional assemblies (between-assembly) carried more joint spatial information (JMI) than pairs drawn from the *same* assembly (within-assembly; Fig. 2A), and the additional information was synergistic: between-assembly pairs exhibited more negative Redundancy–Synergy Index (RSI) values, indicating that their joint spatial information exceeded what either neuron contributed independently (Fig. 2B). Assemblies are defined as in Fig. 1B and Methods (validation in Supplementary Fig. 2; conceptual overview in Fig. 1C).

This topology-dependent synergy bias was established by a full-factorial linear mixed-effects model (LMM) fitted to all pair-level z-scored metrics across groups and environments, in which topology (within vs. between assembly) was the single strongest fixed effect for both JMI 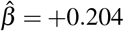, LMM-subsampling 95% interval [0.189, 0.219]) and RSI 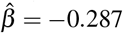, LMM-subsampling 95% interval [−0.296, −0.278]; both intervals exclude zero; nominal Wald *P <* 10^−100^; Fig. 2C), surviving alongside age, genotype, and environment as competing predictors. Assembly membership therefore outranked every biological factor we varied as a predictor of both the amount of spatial information a pair carried and its mode of sharing. This primacy is in line with the long-standing view of cell assemblies as the fundamental unit of hippocampal coding^12,13^.

### The topology-dependent synergy advantage collapses only when ageing and amyloid pathology coincide

Disruption of the topology-dependent advantage was detectable only where ageing and amyloid pathology coincided; neither factor on its own produced a detectable reduction at the present per-group sample sizes. Across the four experimental groups (young wild-type WTY, young 5xFAD ADY, aged wild-type WTO, aged 5xFAD ADO), the topology × genotype × age interaction was the single largest interaction coefficient in the full-factorial LMM, for both JMI 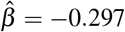 LMM subsampling interval [−0.323, −0.267]) and RSI 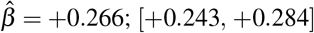; Fig. 3C), whereas the constituent two-way interactions (topology × genotype and topology × age) were smaller and opposite in sign: the coefficient pattern expected if either factor alone is partly compensated and the advantage fails only when they act together. Among the main effects, ageing was the proximate driver of the RSI shift 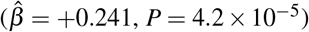, further amplified in the aged 5xFAD background. Topology-dependent spatial coding therefore collapsed where ageing and amyloid pathology coincided, consistent with a non-linear, two-hit interaction at the circuit level.

The per-group distributions shift in the same direction: ADO showed the smallest between-assembly JMI advantage, and WTO and ADO RSI distributions were displaced toward zero relative to the synergy-biased (negative) values of WTY and ADY, indicating reduced synergy bias and a partial shift toward redundancy-dominated coding in the aged groups (Fig. 3A,B; per-group within- vs. between-assembly distributions in Supplementary Fig. 6). No single pairwise contrast against WTY reached significance at the present per-group sample sizes (*n* = 8–13) for either between-assembly JMI (ADY *P* = 0.48, WTO *P* = 0.48, ADO *P* = 0.15; two-sided hierarchical bootstrap, *B* = 10,000) or RSI (ADY *P* = 0.82, WTO *P* = 0.24, ADO *P* = 0.11). These are failures to reject at *n* = 8, not evidence of equivalence, and the factorial interaction rather than any pairwise comparison carries the claim: the plotted distributions carry between-animal uncertainty from the hierarchical bootstrap while the stars index the pair-level LMM (Methods; Supplementary Fig. 8).

### Redundancy and synergy are disrupted through mechanistically distinct pathways

The two constituent atoms, redundancy and synergy, proved to be organised, and to fail, in different ways: redundancy carried a topology main effect that was attenuated in aged 5xFAD CA1, whereas synergy carried no such main effect and instead lost its environmental gating. The RSI integrates the two into a single index, so each atom was modelled separately.

Redundancy showed a topology main effect in healthy networks: between-assembly pairs carried significantly less redundant spatial information than within-assembly pairs in WTY animals (Fig. 4A; LMM topology main effect 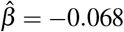, 95% CI [−0.079, −0.057], *P* = 8.3 × 10^−30^). This difference was attenuated in ADO animals, where within- and between-assembly redundancy distributions largely overlapped, consistent with a dissolution of assembly boundaries as the modular network structure degraded (Supplementary Fig. 11). Because the MMI redundancy function provides an upper bound on redundancy^51^, an alternative decomposition such as *I*_BROJA_^55^, for which robust estimators exist^56^, can only lower the redundancy estimates, so the magnitude but not the direction of this topology effect could differ; the RSI result above is defined as interaction information and is therefore lattice-independent and unaffected by this choice (see Methods, “Partial Information Decomposition (PID)”).

Synergy, by contrast, showed no topology main effect in any group: within- and between-assembly pairs carried equivalent synergy in both WTY and ADO animals (Fig. 4B; LMM: 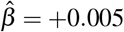, 95% CI [−0.006, +0.017], *P* = 0.43). Synergy was therefore not organised by assembly membership in the simple within-versus-between sense. Instead, synergy expressed its topology dependence through a *context-sensitive* mechanism captured by the four-way interaction in the LMM (see below). The topology-dependent synergy index (between − within) was preserved (around zero or slightly positive) in WTY in both environments and in ADO-Familiar; it collapsed selectively in ADO-Novel, where between-assembly synergy fell below within-assembly synergy (Fig. 4C; LMM synergy four-way interaction, * * *). The effect was carried by the between-versus-within contrast rather than by a change in average synergy magnitude.

The LMM confirmed this mechanistic asymmetry at the level of the *main* effects: redundancy carried a significant topology main effect 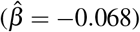 whereas synergy did not 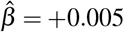, *P* = 0.43). Both atoms additionally carried a significant four-way topology × environment × genotype × age interaction (synergy 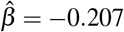 95% CI [−0.291, −0.114], *P* = 1.3 × 10^−6^; redundancy 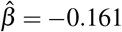, *P* = 6.8 × 10^−21^), so context-dependent gating is not unique to synergy. The dissociation is therefore that redundancy is organised by assembly membership at baseline (a topology main effect) while synergy is not, even though both atoms lose their topology-dependent structure under the combined four-way perturbation (Fig. 4D; full fixed-effect coefficient overview in Supplementary Fig. 10).

### Modularity and small-world organisation show a genotype × age deficit in aged 5xFAD CA1

The functional backbones of aged 5xFAD animals were less modular than the additive expectation of ageing and genotype, so the pair-level coding deficits are accompanied by a measurable change in network architecture. Each backbone was characterised by its modularity (*Q*), weighted clustering, average path length and small-world coefficient (*σ*), each modelled with a robust Huber regression (Methods); the backbone is connected by construction (Methods), so the PID deficit is not a trivial consequence of fragmentation.

Modularity is evaluated on the same Markov-stability partition that defines the assemblies used for the PID analysis, so structure and coding are read from one set of communities rather than two, and is reported as the excess over a strength-preserving rewired null, Δ*Q*^str^ = *Q*_obs_ − ⟨*Q*_null_⟩ (Methods). What carries the comparison is therefore how far each session sits above its own null, not the absolute value.

The deficit takes the form of a negative genotype × age interaction (Fig. 5A,B). Δ*Q*^str^ rose with age 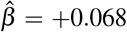, *P* = 0.002), while the 5xFAD genotype main effect was not significant on its own 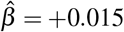, *P* = 0.48); the aged-5xFAD combination fell far below the additive prediction (interaction 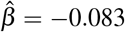, 95% CI [−0.144, −0.021], *P* = 0.008). Aged 5xFAD animals, in other words, fail to gain the age-driven modularity increase seen in wild-type. Two related modular-family metrics moved in the same direction: weighted clustering and the small-world coefficient (*σ*) each showed a same-sign, subadditive genotype × age interaction (Supplementary Fig. 11). On the strength-preserving-null *z* scale, however, only modularity reached significance; clustering and *σ* each trended negative but individually did not 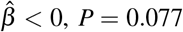 and 0.079; small-world *σ* interaction 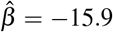 on the *z* scale, though *σ* was significant on the ratio scale, *P* = 0.043; full eight-metric model in Supplementary Fig. 11), and average path length carried no significant term under the Markov-stability community-count covariate. We therefore read this as one individually significant metric (modularity) supported by a convergent, direction-level shift across the modular family, rather than as three independent effects; because modularity’s nominal *P* = 0.008 would not survive Bonferroni correction over the eight metrics, the coherence of direction, not any single metric, carries the argument. An apparent age × novelty gating of *σ* was not robust (Methods), leaving a non-novelty structural signature.

The reduced modular backbone was present in the same animals that lost topology-dependent coding (Fig. 4). An exploratory within-group analysis linked the two measures directly: session-level Δ*Q*^str^ correlated positively with topology-dependent RSI (within − between) in the familiar environment (Fig. 5C) in both 5xFAD groups, young (ADY, *r* = 0.88, *n* = 8, *P* = 0.004) and aged (ADO, *r* = 0.56, *n* = 13, *P* = 0.046), but in neither wild-type group (*r* = 0.18 and *r* = 0.22, both ns). Given the small *n* and the four uncorrected within-group tests, of which only the young-5xFAD coupling survives Bonferroni correction, we treat this as hypothesis-generating.

### Multivariate community-level emergence confirms the pair-level collapse

The pair-level collapse extends to higher-order information dynamics: at the community scale, ageing produced a coherent multivariate signature, which the genotype × age interaction reversed. Pairwise PID captures the redundancy/synergy structure of two-neuron interactions but cannot resolve information genuinely distributed across three or more units, so we computed three community-level emergence measures on the assembly partition time-series of each session: causal emergence Ψ^57, 58^, downward causation Δ^57, 58^, and O-information^35^. Each has a directional reading. Positive Ψ means the community state predicts its own future beyond what the individual neurons predict, so the macroscopic description carries causal information the parts do not; positive Δ means the community state predicts an individual neuron’s future beyond what the other neurons predict, that is, macro-to-parts influence. Negative O-information indicates predominantly synergistic higher-order interactions and positive O-information predominantly redundant ones^35^. Ψ and Δ were compared against time-shifted surrogates and O-information against column-permuted surrogates (per-delay surrogate-median subtraction for Ψ and Δ; standardisation by surrogate standard deviation for O-information; see Methods) and entered into a robust linear model (Huber M-estimator) with full three-way interactions among genotype, age, and environment, controlling for the number of communities per session (*N* = 77 sessions for Ψ/Δ; *N* = 75 for O-information, two sessions with undefined surrogate-normalised score excluded; Fig. 6; per-group integrated surrogate-corrected scores in Supplementary Fig. 13).

Integrated causal emergence Ψ (per-session integrated surrogate-corrected score; sum over delays of [observed surrogate median]) increased with age (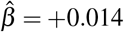, *P <* 10^−3^), as did downward causation Δ (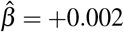, *P* = 0.022): aged networks carry more whole-system causal information and more macro-to-parts influence than young ones. O-information became significantly more negative with age (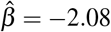, *P* = 0.028), a shift toward synergy-dominated multivariate integration. This shift was reversed by the genotype × age interaction (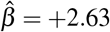, *P* = 0.049), restoring redundancy at the community scale specifically in aged 5xFAD animals. No other terms (genotype main, environment main, or higher-order interactions) reached significance. Although the two O-information coefficients are individually marginal (*P* = 0.028 and *P* = 0.049), they are corroborated by the convergent, independently surrogate-corrected Ψ (*P <* 10^−3^) and Δ (*P* = 0.022) age effects, which point in the same direction (increased higher-order integration with age) under an outlier-robust estimator.

The age effect at the community scale was opposite in sign to the pair-level age effect on RSI (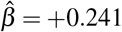, *P* = 4.2 × 10^−5^, i.e. toward redundancy; Fig. 3). The genotype × age interaction was disruptive at both scales: pair-level synergy lost its environmental gating (Fig. 4, four-way interaction 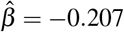, *P* = 1.3 × 10^−6^), while multivariate O-information reversed toward redundancy (Fig. 6F; 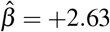, *P* = 0.049).

## Discussion

### Network topology as an organising principle of hippocampal spatial coding

Functional community structure constrains the *mode* of information sharing in CA1, not just its magnitude (Fig. 2), which moves the account of hippocampal dysfunction beyond single-neuron descriptions to a circuit-level computational failure. The relevant variable is not simply which neurons are active, but which assembly they belong to. Because topology outranked genotype, age, and environment in a model that fitted all of them simultaneously, it is better read as an organising principle of spatial coding than as one covariate among many. This extends prior work on hippocampal assembly coding^14,50^ by showing that the topological identity of a neuron pair (whether it spans assembly boundaries) is a stronger determinant of its coding strategy than the biological state of the animal.

The division of labour it imposes is an intuitive one: dense local connectivity supports a noise-robust redundant readout within an assembly, while cross-assembly interactions generate genuinely new representational content. The same division has been described across cortical systems, where redundant interactions concentrate in modular, structurally coupled circuits while synergistic interactions predominate in the higher-order networks that support integrative computation^36^. PID atoms have previously been related to a neuron’s position in a functional network^59^, correlated activity has been shown to favour synergistic processing in cortical cultures^60^, and synergistic subsystems sit between canonical networks in the human cortex^37^. To our knowledge, however, the equivalent decomposition has not been resolved within a single hippocampal subfield at the level of individual neuron pairs, nor related to assembly boundaries.

### Spatial coding collapse requires both ageing and amyloid pathology

That the collapse in topology-dependent spatial coding is selective to aged 5xFAD animals is a central finding (Fig. 3), and it is consistent with a two-hit model in which age-related network deterioration and amyloid-driven pathology each produce subthreshold perturbations that, when combined, cross a critical threshold for topology-dependent failure. The model makes a structural prediction that the coefficients bear out: were either factor sufficient on its own, its two-way interaction would carry the same sign as the three-way term and the single-factor groups would show intermediate deficits. Neither holds here, so the collapse behaves as a threshold crossing rather than as the accumulation of two independent linear insults, a statement about the shape of the interaction rather than about equivalence of the single-factor groups. Two-hit formulations of this kind, in which neither insult is sufficient alone but their convergence is pathogenic, have been proposed for Alzheimer’s disease at the molecular level^61^. The present data extend that logic to circuit computation, a level at which network dysfunction and ageing are each established contributors in their own right^62,63^: the two hits are ageing and amyloid pathology, and what they jointly disable is the topology dependence of information sharing rather than any single cellular process.

### Redundancy and synergy fail through distinct mechanisms

The two atoms fail by mechanistically different routes (Fig. 4), and the dissociation is legible in which term of the model carries each failure. Redundancy is organised by assembly membership at baseline, so its failure registers as the loss of that main effect: a *structural* failure, in which the dissolution of assembly boundaries erases the distinction between a within- and a between-assembly pair, in step with the reduced modular backbone measured in the same animals (Fig. 5). Synergy carries no baseline topology main effect and therefore cannot fail in the same way. Its topology dependence resides entirely in the interaction with environment, so its failure is the loss of that gating: a *functional* failure, in which average synergy is preserved but the network can no longer up-regulate on demand the cross-assembly integration that new spatial-map formation would otherwise draw on. The functional deficit is the more subtle of the two: because it is defined by the contrast between environments rather than by a change in level, it is invisible to analyses that average over environmental context, which may be one reason that AD-related coding deficits have more often been characterised as losses of magnitude than as losses of context-specificity. Both routes are also consistent with reports that single-cell spatial tuning in this model can remain largely intact^64^: each is a property of how a *pair* of neurons divides information about position, and neither requires the individual tuning curves to have degraded.

A convergent picture is emerging in humans. Applying integrated information decomposition to resting-state fMRI across cognitively healthy, mildly impaired, and Alzheimer’s populations, Down et al.^54^ report a widespread reduction in synergistic information alongside increased redundancy in patients. The measurement scale could hardly be more different from ours, whole-brain haemodynamic signals in people versus pairs of imaged CA1 neurons in mice, yet the direction of the shift is the same. Our data locate one circuit-level instance of that shift and add that it is not uniform, since the synergy loss appeared specifically when the network had to integrate a novel environment.

### Parallel structural and functional decline, with structure–function coupling in 5xFAD animals

Structure and function decline in parallel: the same animals that lose topology-dependent synergy at the pair level also build less modular backbones (Fig. 5). This parallel is what the mechanistic account above requires. If redundancy is organised by assembly boundaries, then a network that fails to maintain those boundaries should lose the redundancy contrast, and the modularity deficit is the structural measurement of exactly that failure. Because the structural signature carries no novelty dependence, pair-level synergy gating remains the load-bearing context-dependent effect (Fig. 4).

The within-group correlation between modularity and topology-dependent coding was positive in both 5xFAD groups and absent in both wild-type groups (Fig. 5C), which would place structure–function coupling under genotype-specific control rather than making it a general property of CA1. We treat this as a working hypothesis rather than a result: on four small, uncorrected within-group tests, distinguishing a genotype-specific coupling from a coupling that is simply easier to detect where both measures vary more will require independent cohorts.

### Cross-scale convergence of the information-theoretic collapse

Ageing redistributes CA1 information toward higher-order synergistic integration even as pair-level redundancy rises, and the genotype × age interaction reverses that redistribution (Fig. 6). This is independent multivariate corroboration of the pair-level collapse (Fig. 4) and of the structural deficit (Fig. 5). Collective dynamics that are visible only at the population scale have been a recurring theme in cortical circuits since the description of neuronal avalanches^65^, and the causal-emergence framework used here was developed by Rosas et al.^57^ and has been adapted to biological^58^ and artificial neural networks^66^; applying it to spatial-coding assemblies shows that such emergence is not fixed but is selectively reorganised by ageing and amyloid pathology.

The opposite age signs at the two scales (pair → redundancy, community → synergy) are scale-complementary rather than contradictory. The two analyses ask different questions of the same sessions: pairwise PID asks how two neurons divide the information they carry about position, while the emergence measures ask what the assembly ensemble as a whole predicts beyond the sum of its parts. A network that duplicates spatial information across neurons within an assembly is, at the same time, a network whose assemblies become individually more reliable units, and reliable units are the substrate on which higher-order, distributed structure can be built. Compensatory pair-level redundancy can in this way enable emergent multivariate synergy among assemblies^35^, an interplay of functional segregation and integration that has long been proposed as the organising axis of cortical complexity^67^, so the two age effects are better read as one reorganisation observed at two scales than as two opposing effects. What distinguishes the aged 5xFAD condition is that the genotype × age interaction disrupts both scales at once, which argues that the amyloid + ageing combination constitutes a coordinated reorganisation of CA1 information dynamics rather than a simple loss of function.

### Limitations and future directions

One limitation of our study is that statistical power is reduced by the fact that young and old 5xFAD mice are separate groups. A more powerful, albeit more experimentally challenging, approach would be to image longitudinally across age groups in the same mice. This was unfortunately not feasible in our case. While such an experimental design would improve statistical power, it would also have to contend with the slow turnover of the CA1 place code across days^7^, which may limit gains in interpretability. The 5xFAD model itself is limiting. While it is amongst the most prevalent mouse models of AD, it models aggressive amyloidosis (through overexpression of five familial AD mutations) without tau pathology, and has shown limited clinical translatability to date. Future work might make use of other models such as the APP NL-G-F model^68^. In addition, we did not here quantify amyloid burden per animal, which might have improved statistical power instead of using genotype as a proxy for pathology^69,70^.

Several analytical choices may also limit interpretability. Assemblies here are functional communities detected from coactivity, not anatomically defined ensembles, so “assembly boundary” means a statistical partition of the recorded population rather than a histological structure, with the usual caveats on modularity maximisation^71,72^, small-world summary statistics^73^ and the choice among assembly-detection schemes^74^. The redundancy function bounds the magnitude, though not the direction, of the redundancy topology effect (Results). Pairwise PID also cannot resolve information distributed across three or more neurons, which is what motivated the community-level emergence analysis; a full higher-order decomposition remains to be performed. Our analysis also makes use of deconvolved calcium event amplitudes (from OASIS), rather than electrophysiologically defined spikes, and slow calcium kinetics together with the 33 ms frame resolution result in limited temporal event resolution. This can inflate apparent pairwise dependence, which is why every quantity is reported against its own circular-shift surrogate distribution. The *z*-scores measure departure from a null preserving each signal’s autocorrelation rather than information in absolute bits, and the topology contrasts are differences on that common scale.

Every metric here is computed over a whole session and is therefore a static summary, yet the process the novel environment engages, encoding a new spatial map, unfolds dynamically over the first minutes of that session. The collapse of (between − within) in the novel environment in old AD mice is thus revealed by a single value averaged across the whole recording. The between-assembly deficit may be present from the moment the animal enters the novel environment, or emerge only as the map begins to stabilise, or appear and then decay. This reduces sensitivity. Resolving PID atoms from shorter segments within a session would distinguish these; however, this may require improved PID estimators.

We have made use of the MMI (Minimum Mutual Information) redundancy function in our PID estimation strategy, because it is the most interpretable function and gives a non-negative decomposition; however, other functions could be used. We nevertheless validated our findings against the Redundancy-Synergy Index, which does not require a redundancy function, and found that they held up.

Reading hippocampal coding through the topology of its assemblies, in summary, turns a diffuse account of dysfunction into a specific one: what ageing and amyloid jointly disable is not only the amount of spatial information CA1 carries, but also the way neurons interact within the functional network architecture. This reframes the failure from neuron to circuit level, and may yield new candidate targets for therapeutic development.

## Methods

### Animals and Data Acquisition

Detailed experimental procedures, including animal husbandry, cranial window surgery, and two-photon calcium imaging, were performed as previously described^21^. We analysed CA1 pyramidal-cell activity recorded during active exploration in **wild-type (WT)** mice and the **5xFAD** transgenic mouse line^75^, an established amyloidosis model (mutant human APP/PSEN1, progressive amyloid-*β* deposition) that defines the genotype factor (5xFAD vs. wild-type throughout; labelled *AD* vs. *WT* in figures and statistical tables). The 5xFAD genotype is the experimental driver of amyloid pathology in this model; we did not directly quantify per-animal amyloid burden, so genotype serves as the proxy for amyloid pathology and genotype-associated effects are attributed to amyloid pathology (reported by genotype and age). Four groups were studied: young wild-type (WTY), young 5xFAD (ADY), aged wild-type (WTO), and aged 5xFAD (ADO). Data were analysed from 26 mice in total: 6 WTY, 6 ADY, 8 WTO and 6 ADO^21^. Ages at imaging were 1.7–2.0 months for the young groups and 5.6–9.3 months for the aged groups (verified from per-animal acquisition records). Imaging was performed at ~ 30 Hz^21^ using a resonant-scanning two-photon microscope with a 16 × /0.8 NA water-immersion objective. Neurons expressed the genetically encoded calcium indicators jGCaMP7s^76^ or GCaMP6s^77^, as described previously^21^. Mice were trained to run laps on a familiar circular track (fam1) over several days of habituation, with water reward used to motivate running; CA1 activity was then imaged both during re-exposure to this familiar environment (engaging retrieval of an established representation) and during exploration of a novel environment (the same track with altered cues), which engages encoding of a new spatial memory, whose newly formed assembly code we read out (decode)^21^. Session counts: WTY *n* = 11 familiar, *n* = 11 novel; ADY *n* = 8 familiar, *n* = 8 novel; WTO *n* = 8 familiar, *n* = 5 novel; ADO *n* = 13 familiar, *n* = 13 novel (total: 40 familiar + 37 novel sessions). WTO novel sessions (*n* = 5) reflect available recordings passing quality control from this group; WTO familiar (*n* = 8) were all included. To isolate spatial coding epochs and mitigate behavioural confounds, analysis was restricted to periods when running speed was 20–500 mm s^−1^ (Gaussian smoothed, *σ* = 2.5 samples). Position entropy was confirmed to be matched across groups during these high-speed epochs to rule out spatial-sampling bias.

Calcium fluorescence traces (Δ*F/F*) were deconvolved with the OASIS algorithm^78^ as implemented in the MATLAB version of CaImAn^79^, following motion correction and ROI segmentation in the same pipeline^21^. This yields an event train preserving both the onset time and the amplitude of each inferred calcium transient, removing the slow temporal decay component of the indicator signal. Amplitudes are in arbitrary units inherited from the ROI time series, which is why every metric below is computed on a discretised version of this event train rather than on raw amplitudes. Two discretisations are used, each defined with the analysis that needs it: a five-state amplitude alphabet for the information-theoretic quantities (“Information-Theoretic Calculations”), and a binary active/silent coding for the pairwise couplings that weight the functional network (“Functional Network Construction”).

### Information-Theoretic Calculations

Entropy, mutual information and PID all require a discrete alphabet, so the neural and positional variables were binned before estimation. Continuous deconvolved spike amplitudes were discretised into *k*_neu_ = 5 states using **quantile boundaries pooled across all 40 familiar-environment sessions from all four groups** (*>* 23 million timepoints; Supplementary Fig. 1): bin 0 is the silent state (amplitude = 0), and bins 1–4 are four equal-occupancy quartiles of the nonzero amplitudes (Supplementary Fig. 1A). Global rather than per-session bin edges keep entropy scales comparable across animals, groups and environments, and *k*_neu_ = 5 balances the graded rate information retained against under-sampling of the joint state space 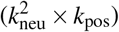. Spatial position along the circular track was discretised into *k*_pos_ = 50 equal-width bins, a resolution of approximately 7.2° per bin. Every information quantity reported below is computed on this alphabet, with one exception: the pairwise mutual information used to weight the functional network is estimated from binarised event trains, for the reasons given in “Functional Network Construction”.

All information metrics were computed using the **Nemenman–Shafee–Bialek (NSB)** estimator to correct for the systematic downward bias in entropy estimation inherent in finite neural datasets^80–89^. The NSB estimator places a Dirichlet mixture prior over the space of probability distributions and integrates over this prior to yield a bias-corrected entropy estimate. Limited sampling is the dominant systematic error in spike-train information estimates, and the practical treatment of that bias follows established guidance^88,90^. General treatments of information-theoretic analysis and decoding in neural populations are given elsewhere^91,92^. All entropies were computed with the `ndd` Python package (v1.10.6)^89^, which evaluates the identical mixture-of-Dirichlets NSB estimate from the *multiplicities* (counts of counts) of the sampled distribution via a compiled backend and also returns the posterior standard deviation of each estimate. This is a numerically optimised reimplementation of the original NSB estimator^82^, not a different estimator: the statistical mechanism (mixture-of-Dirichlets prior, integration over the concentration parameter) is unchanged, but the counts-of-counts representation makes it tractable in the under-sampled regime. Alphabet sizes were supplied explicitly to the estimator: *k* = 5 for spike marginals, *k* = 25 for the joint spike distribution, and *k* = 1,250 for the full triplet *H*(*S*_1_, *S*_2_, *T*). Pairwise mutual information was calculated as:

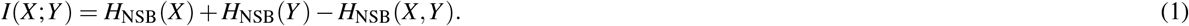

### Partial Information Decomposition (PID)

To dissect how pairs of source neurons (*S*_1_, *S*_2_) encode the target spatial position (*T*), we used the **Partial Information Decomposition (PID)** framework^46^. The joint mutual information *I*(*S*_1_, *S*_2_; *T*) was decomposed into four atomic components: redundant information shared by both sources (Π_red_), unique information from each source (Π_unq1_, Π_unq2_), and synergistic information available only from their joint state (Π_syn_).

We implemented the **Minimum Mutual Information (MMI)** redundancy function^51^, in which redundancy is defined as the minimum information provided by either source about the target:

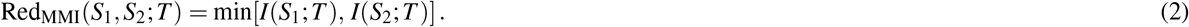

Because unique information is non-negative, Red ≤ min[*I*(*S*_1_; *T*), *I*(*S*_2_; *T*)] in any decomposition satisfying the standard axioms, so this choice provides an *upper* bound on redundancy and a correspondingly lower bound on synergy. It is also computationally tractable for the large number of neuron pairs analysed here. Synergy is then obtained by subtracting the larger single-source information from the joint information:

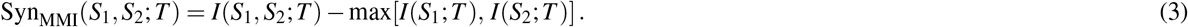

To complement the lattice-dependent PID atoms, we also computed the **Redundancy–Synergy Index (RSI)**, defined as the interaction information:

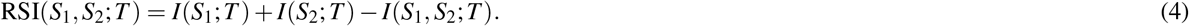

RSI depends only on Shannon mutual information terms and is therefore independent of the specific PID lattice choice. Under the MMI formulation used here, this quantity is algebraically equal to the difference between redundancy and synergy

(RSI = Red_MMI_ − Syn_MMI_). Positive RSI values indicate redundancy-dominated coding, whereas negative values indicate synergy-dominated coding. Because RSI is a standard Shannon-information quantity, our headline results (topology-dependent shifts in redundancy–synergy balance) do not depend on the choice of PID decomposition.

To verify that the MMI–PID readout recovers known information structure, we ran ground-truth simulations using canonical logic-gate benchmarks^46,92^. Simulated Poisson spike trains driven by binary latent states were discretised through the same global equal-occupancy *k* = 5 amplitude binning used for the real data and passed through the MMI–PID estimator. A COPY gate, in which one neuron copies another’s target encoding, yields redundancy-dominated coding; an XOR gate, in which two neurons share a latent state whose combination determines the target, yields synergy-dominated coding in which redundancy falls to zero. The estimator recovered both regimes, and Markov stability resolved a designed multiscale community structure across Markov times whereas single-resolution Louvain did not (Supplementary Fig. 5; full construction in Supplementary Methods, “ground-truth simulation model and logic-gate benchmarks”).

#### Surrogate Normalisation and z-Scored PID Metrics

All PID and mutual-information quantities were surrogate-normalised before statistical analysis. For each neuron pair, 200 circular-shift surrogate datasets were generated by circularly shifting the target position time series relative to the aligned neural activity by a random offset drawn uniformly over the recording duration; for these multi-minute recordings the offset almost always far exceeds the autocorrelation timescale of the position signal. This procedure destroys neuron–position relationships while preserving the autocorrelation structure and cross-neuronal temporal dependencies of the neural signals^92^.

Each PID metric *M* (JMI, redundancy, synergy, RSI) was recomputed for every surrogate, yielding a shuffle distribution with mean *µ*_shuf_ and standard deviation *σ*_shuf_. We report **z-scored, surrogate-normalised metrics** rather than raw information values (bits):

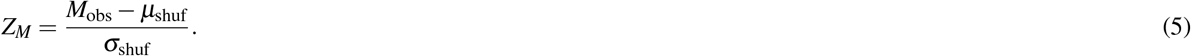

Thus, all downstream analyses (hierarchical bootstrap and linear mixed-effects models) were performed on surrogate-normalised z-scores, not on raw information estimates. This normalisation controls for finite-sample bias and pair-specific baseline structure in information estimates: dividing by the per-pair shuffle standard deviation places every pair on a common scale where one unit equals one shuffle-distribution standard deviation, making metrics directly comparable across pairs, sessions, animals, groups, and environments. A z-scored value of zero indicates that the observed metric is indistinguishable from the circular-shift null, while large positive (or negative) values indicate spatial information content (or structure) beyond that expected by chance.

### Topological Assembly Analysis

#### Functional Network Construction (Significance-Constrained Backbone)

Network edges are built from a deliberately coarser representation of the same event trains: for this step the deconvolved estimates were **binarised** (*k* = 2 states, silent/active) on the presence of any event within a 33 ms window (one imaging frame). What an edge has to report is whether two neurons are co-active more often than chance, not how much their graded amplitudes say about position, and a binary alphabet is better matched to that question: it reduces the pairwise state space from 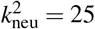 to 4, so each edge weight and its surrogate distribution are estimated from far more samples per state, at a scale where one *N* × *N* matrix of edges, each with its own shift test, must be estimated per session. It also keeps amplitude fluctuations of the deconvolution output, which are in arbitrary units, from being read as coupling strength. The graded *k*_neu_ = 5 alphabet is retained where amplitude resolution is load-bearing, in the entropy, MI and PID estimates themselves (“Information-Theoretic Calculations”).

Functional connectivity was quantified using pairwise mutual information *I*(*S*_*i*_; *S* _*j*_) between these binarised spike trains, estimated via the NSB estimator. Statistical significance of each edge was assessed via 200 circular shifts of one spike train relative to the other, with a minimum shift of ± 10 s.

To recover topological features while controlling for the high noise floor of pathological networks, we used a **significance-constrained hybrid filtering strategy** that combines local geometric constraints with edge-wise significance testing:

1. The MI matrix was first restricted to statistically significant edges (*P <* 0.05 vs. circular shuffles). **Continuous k-Nearest Neighbours (CkNN)**^93^ with *k* = 20 and *δ* = 1.0 (robustness of the structural result to this choice: Supplementary Note, “robustness of the modularity deficit to the CkNN neighbourhood size”) was then applied to the distance graph derived from this significance-filtered matrix. Unlike global thresholding, CkNN adapts edge density to the local neural manifold and can therefore recover structure even in sparsely active sub-circuits, while operating only on couplings that already passed the significance test.
2. To prevent the network from fragmenting into isolated components (a prerequisite for the ergodicity of the random walker used in community detection), the **Minimum Spanning Tree (MST)** of the same significance-filtered MI matrix was superimposed onto the graph. The MST preserves global connectivity with the minimum total edge weight, so even weakly connected modules remain reachable by the diffusion process; where an MST bridge spanned a pair carrying no significant MI, it was assigned a small nominal weight (0.1× the minimum non-zero backbone weight) rather than its raw MI, restoring reachability without adding a spurious strong edge.

The final adjacency matrix combined the CkNN and MST edges, both computed on the significance-filtered matrix:

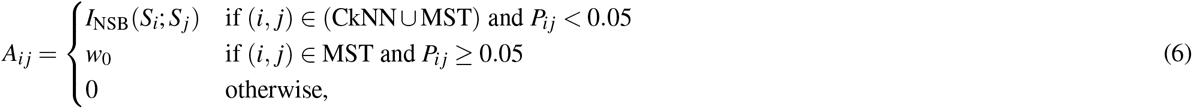

where 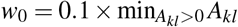 is a small nominal bridge weight. Because CkNN and MST both operate on the significance-filtered matrix, every non-bridge edge carries a coupling that passed the significance test, while the MST (with nominal weights on the few bridges required) guarantees global reachability.

#### Multiscale Community Detection

Functional assemblies were identified using the **PyGenStability** framework^94^, an implementation of Markov-stability community detection^52,53^: communities are found by tracking how long a diffusion (random walk) remains trapped within candidate modules as a function of Markov time *t*, a single continuous resolution parameter (small *t* yields many fine communities, large *t* a few coarse ones). The random-walk construction and the Markov-stability objective it optimises are given in Supplementary Methods (PyGenStability community detection).

Starting from each session’s significance-constrained backbone **A**, Markov times *t* ∈ [10^−1^, 10^1.5^] were scanned on a logarithmic grid of 50 points, using the continuous normalised Markov-stability constructor. At each *t* the stability objective was optimised with the Louvain algorithm^95^ using *N*_tries_ = 50 random restarts (Louvain is the optimiser supplied by PyGenStability; the Leiden refinement^96^ guarantees connected communities but was not required here, as the analysed partitions were checked for cross-scale and across-restart consistency) to account for near-degenerate solutions, and partition reproducibility at that scale was quantified by the Variation of Information^97^ across the restarts, normalised as in ref.^98^. PyGenStability’s block detection then flagged the *optimal scales*: Markov times whose partitions are mutually consistent over a contiguous band (Supplementary Methods); among these, the analysed partition was the scale of *highest Markov stability* returning at most 10 communities. Communities smaller than 5 neurons were finally merged into their strongest neighbour, so that every neuron is retained. The rule involves no per-session hand tuning: its parameters were fixed in advance and applied identically to all sessions and groups, and its robustness is characterised in Supplementary Fig. 3. We further assessed whether the partition depends on the CkNN neighbourhood size *k* by repeating the full pipeline at *k* ∈ {10, 15, 25, 30} for every session and comparing each partition to the analysed *k* = 20 partition by their Normalised Variation of Information^98^. The NVI was flat ( ~ 0.5) with no discontinuity at the operating point and of the same order as the within-session Louvain-restart variability (median 0.34), so community *membership* is moderately *k*-dependent, comparable to the algorithm’s intrinsic run-to-run lability. The network findings are accordingly reported on relabelling-invariant aggregate statistics (the modularity deficit; Supplementary Figs. 3D and 12) rather than on individual community memberships.

#### Population-vector position decoding

To confirm that the detected communities carry decodable spatial information, we decoded the animal’s track position from community-level activity with a cross-validated population-vector (template) decoder, benchmarked against the full neuron population (an upper reference) and a position-shuffled control (chance). Communities decoded position with a median error well below chance in every group, retaining substantial (though coarser) positional information relative to the full population. Full formulation in Supplementary Methods (population-vector decoding of position from community activity); results in Supplementary Fig. 4.

### Statistics and Reproducibility

The dataset comprises 26 mice (13 male, 13 female): WTY *n* = 6, ADY *n* = 6, WTO *n* = 8, ADO *n* = 6. Each animal was imaged in a familiar and a novel environment, giving 77 sessions in total (familiar environment: WTY *n* = 11, ADY *n* = 8, WTO *n* = 8, ADO *n* = 13); a session is one animal in one environment and is the independent biological replicate throughout. Sessions from the same animal in different environments are repeated measures of that animal and are treated as such by the case random intercept and by the case-level resampling described below. No data were excluded except where stated, no randomisation or blinding was applied (group membership is set by genotype and age), and no statistical method was used to predetermine sample size. All tests are two-sided and *α* = 0.05 unless stated otherwise; reported *P* values are nominal and not corrected for multiple comparisons across the fixed-effect family, and exact values are given for non-significant contrasts as well as significant ones.

Every headline claim in this paper is an interaction rather than a single group difference: whether the effect of the 5xFAD genotype depends on age and, at the pair level, whether that dependence itself differs between within- and between-assembly pairs. At each analysis level we therefore fitted a linear model carrying the full set of interaction terms instead of running pairwise tests. The levels differ only in the unit of observation and in whether a random effect is required (Supplementary Table 1). The alternatives considered and the diagnostics behind these choices are set out in Supplementary Methods (“choosing the statistical model at each analysis level”) and Supplementary Fig. 7.

A *topology contrast* throughout this paper is a signed difference between the two topology classes of a pair-level metric, written as a subtraction with the order given explicitly at each use. The order is not the same for every metric: redundancy and synergy contrasts are (between − within), whereas RSI and *Z*-JMI contrasts are (within − between). A contrast of zero means the metric does not separate within-from between-assembly pairs, and the sign of a non-zero contrast is interpreted in the text and figure legend at each point of use.

#### Hierarchical Bootstrap

Neuron pairs are not independent observations: all pairs in a session share an animal, a field of view and a functional network, and pair counts vary more than hundred-fold across cases. Pooling pairs therefore pseudoreplicates^99^, while collapsing each case to a single mean sacrifices power^100^. All distributions and confidence intervals in the pairwise PID figures come from the hierarchical bootstrap of Saravanan et al.^100^: in each of *B* = 10,000 iterations, cases (animal × session units) are drawn with replacement within group, neuron pairs are drawn with replacement within each selected case, the resampled pairs are reduced to a per-case mean, and the iteration statistic is the grand mean over those means. Two-sided *P* values on a difference distribution Δ = Dist_*A*_ − Dist_*B*_ are 2 min(Pr[Δ *>* 0], Pr[Δ *<* 0]), floored at 1*/B*. Cases were resampled identically across conditions for paired contrasts (topology, environment) and independently for unpaired ones (group); comparisons differing in both group and environment were excluded. Calibration under a null relabelling of animals is shown in Supplementary Fig. 8, and the full procedure in Supplementary Methods (“hierarchical (Saravanan) bootstrap for pair-level metrics”).

#### Linear Models

Fixed-effect significance comes from the linear models set out below and never from the bootstrap. All of them share one form: a null-corrected response (pair-level PID atoms as surrogate-normalised *z*-scores; session-level metrics as defined in their own subsections below) regressed on the full factorial expansion of the experimental factors, treatment-coded with reference levels topology = within-assembly, environment = familiar, genotype = WT and age = Young. The intercept is thus the within-assembly WT Young familiar condition and each coefficient is a contrast relative to it with the remaining factors held at their reference levels, so interaction terms quantify how such a contrast changes across levels of another factor. Where an observation is a neuron pair, the model is a linear mixed-effects model fitted by maximum likelihood with a random intercept per case; where an observation is a session, each session contributes one value and a robust linear model (Huber M-estimator) is fitted with no random effect, specified in full under “Network Structural Analysis” and “Community-Level Emergence Measures” below.

Because pair counts are so unbalanced, the pair-level models are fitted 100 times to case-balanced subsamples (at most 5,000 within-assembly and 5,000 between-assembly pairs per case per iteration). Each reported coefficient is the mean across iterations, its standard error the across-iteration standard deviation, and the Wald *z* and two-sided *P* follow from those two quantities; *P* values themselves are not averaged (Supplementary Methods, “balanced subsampling of the pair-level LMM”; stability and convergence in Supplementary Fig. 9). That standard deviation measures estimator stability under pair-count imbalance, not between-animal uncertainty, which is carried by the hierarchical bootstrap, and confidence intervals on the LMM coefficients are accordingly the 2.5th–97.5th percentiles of the per-iteration estimates rather than hierarchical-bootstrap intervals. Reported *P* values are nominal and not corrected across the fixed-effect family, and because these Wald statistics are computed at the pair level their magnitudes are not read as effect strength; the substantive claims rest on coefficient sign and cross-iteration reproducibility together with the animal-level hierarchical bootstrap. Because each pair was first normalised against its own surrogate distribution, between-animal variance accounted for only about 4–13% of total variance across atoms, and conditional on the fixed effects the random-intercept variance was negligible, so the reported contrasts are not driven by between-animal sampling. For synergy and redundancy, the genotype, age and environment main effects and their non-topology interactions were rank-deficient under the case random intercept and were therefore not estimated, so inference on those two atoms rests on the topology-related terms alone.

Throughout the main figures, stars and brackets denote the model fixed-effect test, whereas the plotted distributions and their intervals derive from the hierarchical bootstrap. Direct between-group bootstrap contrasts are stated explicitly where reported and are generally underpowered at the present per-group *n*, so the load-bearing inference is the model term.

### Network Structural Analysis

Eight network topology metrics (modularity *Q*, weighted clustering coefficient, average path length, number of communities, small-world coefficient *σ* ^101^, degree assortativity, transitivity, and global efficiency) were computed for each session’s backbone graph. Modularity^102^ in the structural analysis was computed with the same Markov-stability partition (PyGenStability) used downstream for PID, applied identically to the observed backbone and to each rewired null graph, so that a single community-detection method is used throughout the pipeline; the strength-preserving null bias-corrected modularity is the primary structural endpoint (Fig. 5). For each metric we fitted a robust linear model (Huber M-estimator) with the same factorial structure as the pair-level LMM but using session-level binary predictors, plus a mean-centred covariate for the per-session number of detected communities:

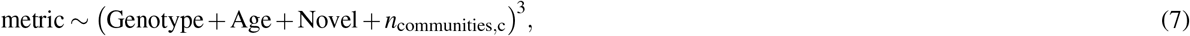

where Genotype = 1 for 5xFAD, Age = 1 for Old, and Novel = 1 for the novel environment, with one observation per session (no random effects required; each session contributes a single observation per metric and predictor combination). The Huber estimator down-weights influential outliers while remaining efficient near Gaussian noise, and the community-count covariate absorbs session-to-session variation in *n*_communities_ that otherwise confounds graph-level summary statistics. We report two-sided Wald *z*-tests on the fixed-effect coefficients.

#### Configuration-Model Nulls (Strength-Preserving)

Because raw graph metrics are dominated by each backbone’s degree and edge-weight sequence, and carry no group structure on that scale (Supplementary Note, “supporting analyses for the network structural results”), we re-evaluated all eight metrics against a strength-preserving configuration-model null (full eight-metric forest in Supplementary Fig. 11E). For each session, the observed weighted backbone was rewired *N*_null_ = 500 times using a custom degree-preserving double-edge swap (Maslov– Sneppen) followed by a weight reassignment that approximately preserves node strength, following the strength-preserving null of Rubinov–Sporns^103,104^, yielding a per-session null distribution for each metric. Null graphs may become disconnected under the double-edge swap; global efficiency is therefore evaluated on the full graph (unreachable pairs contribute 0, which is the standard definition), whereas average path length and the small-world ratio use the largest connected component. We then defined:

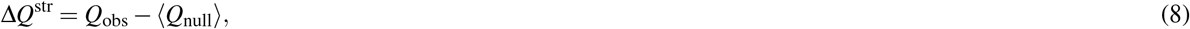

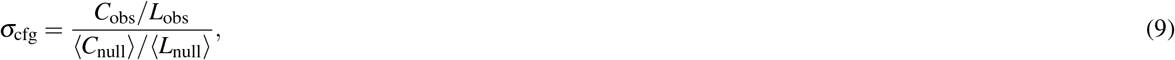

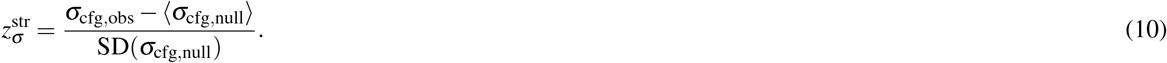

Modularity is null-corrected by subtraction, not by a ratio to the null and not by dividing through SD(*Q*_null_), and the reason is what *Q* already is. Newman modularity is itself a difference: the within-community share of edge weight *minus* its expectation under a degree-matched random graph^102^. It is therefore a residual on an additive scale with a meaningful zero, and the natural further correction on that scale is another subtraction, which leaves Δ*Q*^str^ in units of *Q* and makes each model coefficient readable as a modularity difference. *Q*_obs_*/* ⟨*Q*_null_⟩ would instead be a ratio of two residuals, a dimensionless quantity whose value depends on how large the session’s own null residual happens to be. Clustering, path length and the small-world coefficient are different in kind: they are strictly positive magnitudes rather than residuals, and *σ* is *defined* as a ratio to random graphs^101^, so ratio (and, where the null SD is well behaved, *z*) scaling follows those statistics’ own construction rather than being a scale choice imposed on them.

Two further considerations support the same choice for *Q*. Under the Markov-stability partition the null modularity has a substantially larger and more variable dispersion than under a single-resolution Louvain null, so dividing by SD(*Q*_null_) inflates and destabilises the *z* scale; the subtraction-only correction is scale-immune and matches the convention used for the integrated emergence measures Ψ and Δ (below). And ⟨*Q*_null_⟩ is not constant across sessions (range 0.16–0.42) and is itself correlated with *Q*_obs_ (*r* = 0.63), so a ratio divides the effect of interest by a denominator that partly carries it. The genotype × age interaction is present with the same sign on the ratio scale but only marginally 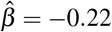, *P* = 0.062, versus 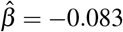, *P* = 0.008 on the subtraction scale), which is a further reason to read the structural result, as we do in the Results, as one metric supported by a coherent direction across the modular family rather than as a single decisive *P*-value. The same Huber-estimator formula was then fit with these null-corrected metrics as dependent variables, and the strength-preserving configuration-model fit is the primary structural analysis reported throughout (Results; Fig. 5; full eight-metric coefficient forest in Supplementary Fig. 11E). Sensitivity of these fits to the community-count covariate, the coefficients on the two small-world scales and the novelty terms are given in the Supplementary Note (“supporting analyses for the network structural results”).

### Community-Level Emergence Measures

Three multivariate emergence measures were computed on the per-community time-series obtained from the optimal Markov stability partition of each session: causal emergence Ψ^57, 58^, downward causation Δ^57, 58^, and O-information^35^. Following the practical criteria of Rosas et al.^57^, 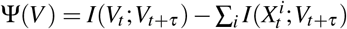 quantifies the predictive surplus of the macro state *V* (the whole-population community signal) over the parts {*X*^*i*^}(the individual per-community signals); 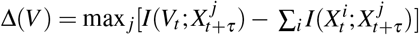 quantifies macro-to-part predictive surplus (downward causation); and the O-information among parts is positive for redundancy- and negative for synergy-dominated higher-order interactions^35^.

The implementation followed previous work^58^, where this framework was applied to population-level neuronal data, using the iterative correction for double counting of redundant information of Sas et al.^105^. For Ψ and Δ, 50 time-shifted surrogates of the community time series were generated per delay; for O-information, 100 column-permuted surrogates per session.

For Ψ and Δ, per-session integrated surrogate-corrected scores were obtained as the sum over delays *τ* ∈ {1, …, 10} of [observed − surrogate median] (baseline-subtracted; *not* divided by *σ*_shuf_, hence retains the units of the underlying mutual information). For O-information, which has no delay axis, a true *z*-score (observed − *µ*_shuf_)*/σ*_shuf_ was computed against the column-permuted surrogate distribution. We refer to all three as “integrated emergence scores” below; only the O-information score is unit-free, so coefficient magnitudes across the three measures are not directly comparable.

The integrated scores were entered into a robust linear model (Huber M-estimator) with fixed-effect predictors Genotype (= 1 for 5xFAD), Age (= 1 for Old), Novel (= 1 for the novel environment), all their two- and three-way interactions, and a mean-centred covariate for the per-session number of detected communities (*n*_communities,c_). *N* = 77 sessions across four groups for Ψ and Δ; for O-information *N* = 75, as two sessions have an undefined surrogate-normalised (*z*) score and drop from that fit. Wald *z*-tests on the fixed-effect coefficients with two-sided *P*-values were used to assess significance.

## Data Availability

Source data underlying all main and Supplementary figure panels, together with the processed per-session PID, network, and community-emergence metrics that enter the statistical models, will be deposited in a public repository upon publication. The raw two-photon calcium-imaging recordings are available from the corresponding author on reasonable request owing to their size.

## Code Availability

The analysis pipeline is provided as the open-source Python package `assembly_pid` (modules: `entropy`, `discretization`, `pid`, `network`, `community`, `community_pid`, `config_nulls`, `statistics`). The package will be deposited on GitHub upon publication. The community-level emergence measures (Ψ, Δ, O-information) were computed with the Causal Emergence Toolbox of Rosas et al.^57^ an Open-Source implementation, with redundancy corrections is available on GitHub.

## Acknowledgements

This work was supported by the Engineering and Physical Sciences Research Council (EPSRC) through the Physics of Life grant [EP/W024020/1]. H.R. is also supported by the Schmidt Sciences LLC., through its AI in Science Research Fellowship.

## Author Contributions Statement

J.J.Y. and H.R. conceived and conducted the computational analysis. M.A.G. performed the experiments and collected the calcium-imaging data. J.J.Y. wrote the manuscript with input from all authors. S.R.S. supervised the project. All authors reviewed the manuscript.

## Supplementary Information

This Supplementary Information comprises 13 Supplementary Figures, one Supplementary Table, six Supplementary Methods sections and two Supplementary Notes. The Methods sections give the full formulation of procedures summarised in the main Methods (multiscale community detection (PyGenStability), population-vector decoding of position from community activity, the ground-truth simulation model and logic-gate benchmarks, choosing the statistical model at each analysis level, the hierarchical (case-level) bootstrap, and the balanced subsampling of the pair-level linear mixed-effects model), and each is placed immediately before the figure it supports. Full fixed-effect coefficients for all four PID metrics are given by the forest plot and effect-direction heatmap in Supplementary Fig. 10A; which terms are estimable, and why the remainder are not, is set out in Methods (“Statistics and Reproducibility”).

**Supplementary Fig. 1.**
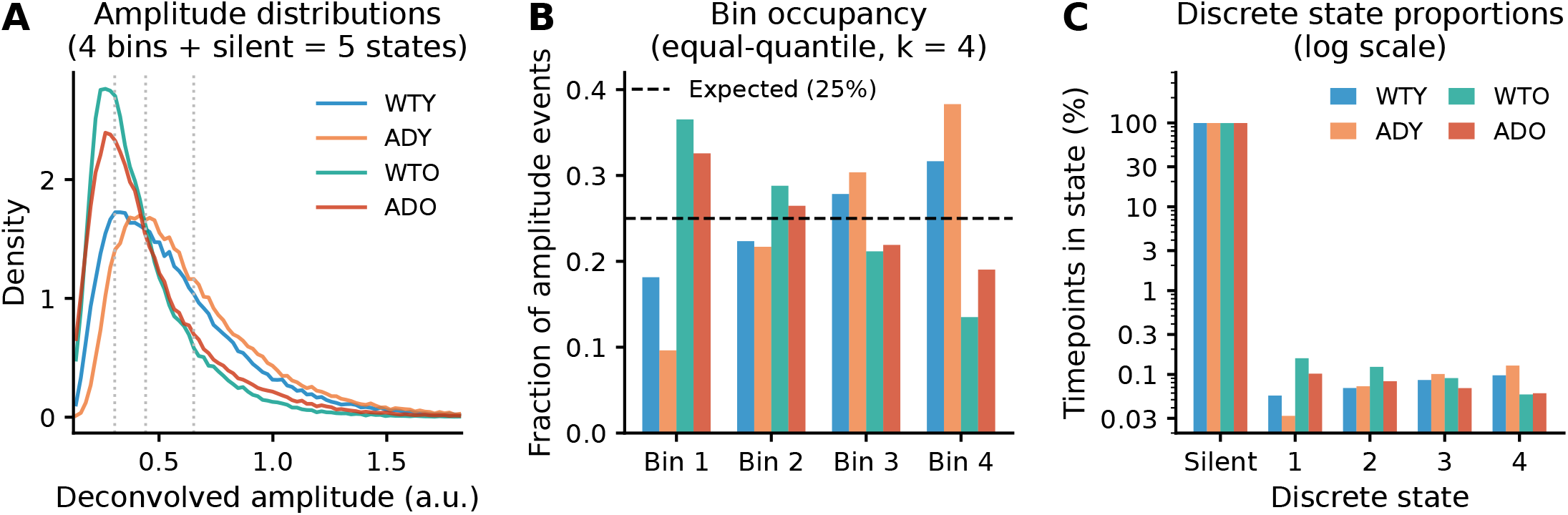
Global discretisation validation. **A**, Deconvolved spike-amplitude density per group, with the three global amplitude-quartile bin edges marked (vertical dotted lines at *Q*_1_ = 0.31, *Q*_2_ = 0.44, *Q*_3_ = 0.65). The total alphabet is *k*_neu_ = 5 states: silent (amplitude = 0) plus four amplitude quartiles. **B**, Bin occupancy per group for the four non-silent amplitude bins; dashed horizontal line marks the expected 25% under equal-occupancy quantile binning. Per-group deviations from 25% reflect group-specific amplitude-distribution shapes; the global edges nonetheless equalise occupancy across the pooled dataset. **C**, Discrete state proportions per group across all five states (silent plus four amplitude bins), on a logarithmic *y* axis. The silent state dominates (99.57–99.69% of timepoints) because most timepoints have zero deconvolved amplitude under the active-epoch filter; the four amplitude states each occupy 0.03–0.16%, a range only visible on a log scale.

**Supplementary Fig. 2.**
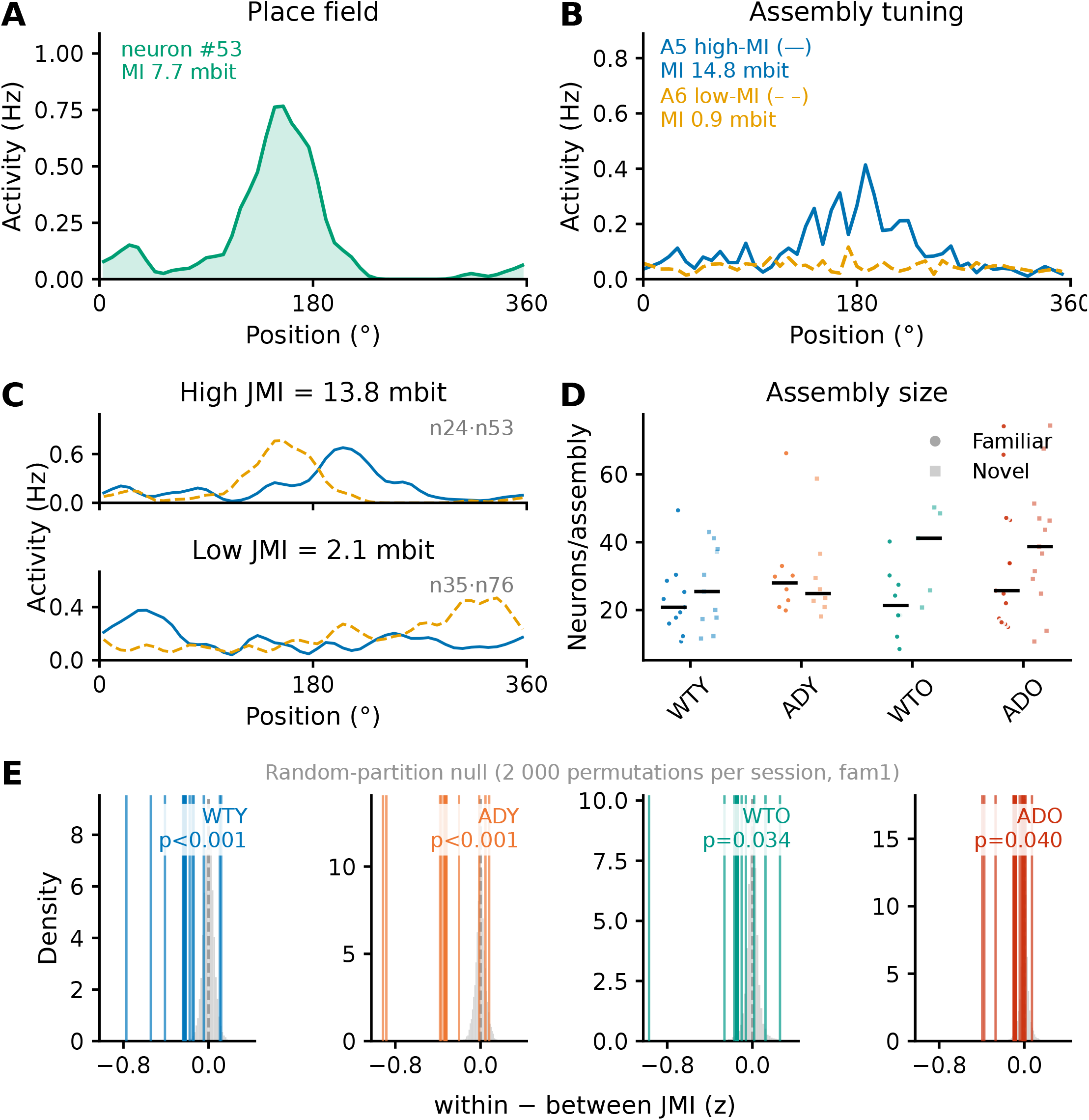
Assembly spatial tuning and partition validation. **A**, Single-cell place field from the example session (case 12): the most informative neuron’s event rate (Hz, circularly smoothed) against track position, annotated with its mutual information with position (*I*(*S*; *T*); neuron n53, 7.7 mbit). Panels **A**–**C** share both axes, activity in Hz and NSB-estimated information in millibits, placing the single-neuron, assembly and pair levels on one scale. **B**, Assembly-level spatial tuning: linearised tuning curves (mean activity, Hz, vs. track position 0–360°) of the highest-MI (blue) and lowest-MI (orange, dashed) assemblies in the same session, accompanying the Fig. 1B partition; each assembly’s mutual information with position is annotated. The high-MI assembly shows a single sharp lobe; the low-MI assembly is broader and lower-amplitude. **C**, Two example neuron pairs from the same session illustrating joint mutual information (JMI; *I*(*S*_1_, *S*_2_; *T*), the information a neuron pair’s joint activity carries about track position). Each panel overlays the two neurons’ spatial tuning (activity, Hz, vs. position; blue solid and orange dashed, neuron identities inset). The high-JMI pair (top; 13.8 mbit, neurons n24/n53, with n53 shown in **A**) has two sharp, near-complementary place fields; the low-JMI pair (bottom; 2.1 mbit, n35/n76) has broad, weakly-tuned activity. **D**, Assembly size (neurons per assembly) per session, by group and environment (filled circles familiar, open squares novel; black bars, group medians). A minimum community size of 5 neurons was applied (smaller communities merged into their strongest neighbour). **E**, Random-partition null model. For each session, 2,000 random partitions of the same community-size distribution were generated and the (within − between) *Z*-JMI gap computed; a positive gap means within-assembly pairs share more spatial information than between-assembly pairs. Coloured vertical lines show the per-session observed gaps and the shaded distributions the per-group surrogate. Observed values fall outside the null at *P <* 0.001 (WTY, ADY), *P* = 0.034 (WTO) and *P* = 0.040 (ADO).

### Supplementary Methods: PyGenStability community detection

#### Rationale and overview of the partition rule

An “assembly” must not be an artefact of a resolution parameter chosen by hand, and the relevant resolution is not known in advance. We therefore wanted a partition rule that (i) is fixed before seeing the data and applied identically to every session, and (ii) returns the scale at which the network’s own diffusion dynamics organise, rather than one arbitrary granularity. Markov stability supplies this: it replaces the fixed resolution of single-scale modularity with a continuous Markov time *t* and identifies the times at which the partition is invariant. Concretely (Methods, “Multiscale Community Detection”), each session’s mutual-information backbone was scanned over 50 logarithmically spaced Markov times; at each time the stability objective was optimised with 50 Louvain restarts; PyGenStability’s block detection flagged the scales that are consistent across neighbouring times; and the analysed partition is the flagged scale of highest Markov stability returning at most 10 communities, after which communities smaller than 5 neurons are merged into their strongest neighbour. This section gives the objective being optimised, the reproducibility measure used to select scales, and how to read the diagnostic scan in Supplementary Fig. 3A.

#### Random-walk model and Markov-stability objective

Starting from the weighted, undirected adjacency matrix **A** (the significance-constrained backbone), define the diagonal strength matrix **D** = diag(**A1**) and the random-walk Laplacian **L** = **I** − **D**^−1^**A**. The continuous-time Markov process has transition matrix **P**(*t*) = *e*^−*t***L**^ and stationary distribution ***π*** = **d***/* ∑_*i*_ *d*_*i*_, where **d** is the vector of node strengths. For a candidate partition *P* the (linearised) Markov stability quantifies the excess probability, relative to chance, that a random walker starts and remains within the same community at time *t*:

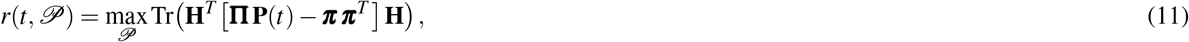

with **H** the community-indicator matrix and **Π** = diag(***π***) (equivalent to the **D**^−1^*e*^−*t***L**^**D** form up to normalisation). As *t* grows the walker explores larger neighbourhoods, so stability is maximised by progressively coarser partitions; the continuous normalised constructor was used throughout.

#### Optimal-scale detection and partition reproducibility

Two labellings *X,Y* are compared by their Normalised Variation of Information, NVI(*X,Y*) = 2*H*(*X,Y*) − *H*(*X*) − *H*(*Y*) */ H*(*X,Y*), the information distance between partitions normalised to [0, 1] (0 = identical; ref.^98^). PyGenStability assembles the cross-scale matrix NVI(*t, t*^′^) over all scanned Markov times and identifies as *optimal scales* the times lying inside low-NVI blocks: contiguous bands of *t* that all return essentially the same partition (a scale-invariant level of structure). These scales are located as minima of the block-averaged NVI. At each individual scale, the NVI across the 50 Louvain restarts measures how reproducibly the optimiser converges. The analysed partition (Methods) is then the flagged optimal scale of highest Markov stability returning at most 10 communities, with communities smaller than 5 neurons merged into their strongest neighbour.

#### Interpretation of the Markov-stability scan

The Markov-stability scan (Supplementary Fig. 3A, the method’s standard output) summarises every candidate partition across scale and is read as follows. The horizontal axis is Markov time *t* (log scale), the single resolution parameter of the method: a random walker explores progressively larger neighbourhoods as *t* grows, so small *t* yields many fine communities and large *t* a few coarse ones.

- **Top panel: available scales**. The Markov-stability objective (blue) measures how well a partition traps the random walk; the number of communities (orange) falls monotonically with *t*. A *plateau* in the number of communities flags a resolution that persists over a range of *t*, a candidate level of organisation.
- **Middle panel: robustness across scales**. The matrix NVI(*t, t*^′^) is the dissimilarity between the partitions obtained at times *t* and *t*^′^ (0 = identical). A *dark, low-NVI square block straddling the diagonal* means a whole band of Markov times returns essentially the same partition, the signature of a robust, scale-invariant level of structure; light regions are transitions between levels. The green curve is the partition’s NVI across the 50 Louvain restarts *at each t*; values near zero mark scales where the optimiser converges reproducibly.
- **Bottom panel: selected scales**. The block NVI condenses the off-diagonal structure around each *t*; its local minima (markers) are the scales PyGenStability flags as optimal, and are drawn as dashed lines across all three panels.

In short, a *trustworthy* partition coincides with a dark diagonal block (agreement across neighbouring scales), a dip in the restart NVI (reproducible optimisation), and a block-NVI minimum, ideally over a plateau in the number of communities. The partition analysed in this paper (solid black line) is the one selected automatically by this criterion (highest Markov stability among the flagged optimal scales, then merging communities *<* 5 neurons; Methods), not a resolution chosen by hand.

**Supplementary Fig. 3.**
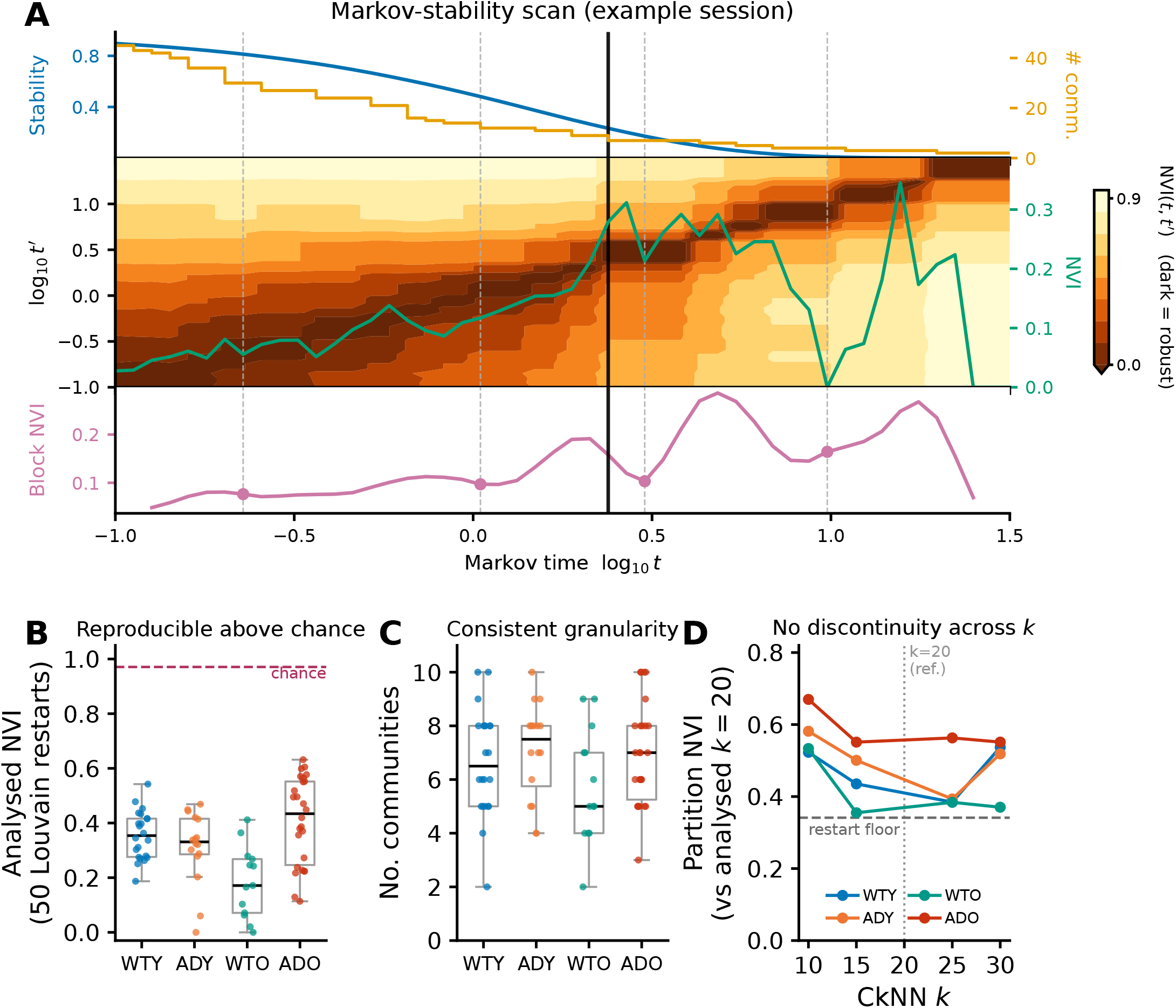
Markov-stability community detection follows a fixed, automated rule and returns reproducible, consistent partitions across all sessions, with no discontinuity at the chosen neighbourhood size *k*. Community detection used an identical, parameter-free selection rule for every session, with no per-session tuning (Methods). **A**, PyGenStability Markov-stability scan for a representative WTY familiar-environment session, produced with the package’s own scan-plotting routine. Top: the Markov-stability objective (blue) and the number of communities (orange) versus Markov time *t*. Middle: the cross-scale matrix of the Normalised Variation of Information between the partitions at times *t* and *t*^′^ (NVI(*t, t*^′^); dark = low NVI), in which robust partitions appear as low-NVI square blocks along the diagonal (a partition stable over a contiguous range of Markov times), with the per-scale NVI across the 50 Louvain optimisation runs overlaid (green). Bottom: the block NVI, whose minima are PyGenStability’s automatically detected optimal scales (markers); these optimal scales are shown as dashed lines across all three panels. The solid black line marks the scale taken downstream (the analysed partition). **B**, Reproducibility of the analysed partition across the 50 Louvain restarts (partition NVI; lower = more reproducible) for all 77 sessions, by group. The median NVI is 0.34, far below the chance level expected for unrelated partitions of the same community-size profile (dashed; random relabelling, NVI ≈ 0.97; cf. the random-partition null, Supplementary Fig. 2E). **C**, Number of communities in the analysed partition per session, by group: a consistent range (2–10 communities) with no degenerate or hand-selected partitions, comparable across genotype and age groups. **D**, Robustness of the partition to the CkNN neighbourhood size *k*. For each session the identical community-detection pipeline was re-run at neighbouring *k* ∈ {10, 15, 25, 30} and each partition compared to the analysed *k* = 20 partition (NVI; *k* = 20 is the self-reference, NVI = 0, so it is marked rather than plotted). The NVI is flat ( ~ 0.5) with no discontinuity at the chosen *k* and of the same order as the within-session Louvain-restart variability (the median of **B**, dashed). Community *membership* is therefore moderately *k*-dependent, which is why network results are reported on relabelling-invariant aggregate statistics rather than on individual memberships (Supplementary Fig. 12).

### Supplementary Methods: population-vector decoding of position from community activity

#### Rationale and overview of the decoding analysis

The communities are built with information-theoretic machinery (a mutual-information backbone partitioned by Markov stability), so we wanted an independent check that they are functionally meaningful and not an artefact of that machinery: if the partition were arbitrary, community activity would not predict the animal’s position better than chance. We therefore decoded the animal’s position on the circular track from community-level activity with a cross-validated population-vector (template) decoder (Supplementary Fig. 4). The decoder is deliberately simple and assumption-light: it learns, per position bin, the mean community activity pattern, and assigns each held-out frame to the bin whose pattern it most resembles, so that any decodability reflects the community code rather than a flexible classifier. The same decoder applied to the full neuron population provides the upper reference, and a position-shuffled run the chance level.

#### Inputs and notation

For each session, neural data were preprocessed exactly as for the information-theoretic analysis (Methods): speed filtering (20–500 mm s^−1^, Gaussian-smoothed speed), the global *k* = 5 amplitude discretisation, and binning of the circular position into *N*_*θ*_ = 50 equal-angle bins of width Δ = 360°*/N*_*θ*_ = 7.2°. This yields, for *t* = 1, …, *T* retained frames, a discrete position *b*_*t*_ ∈ {0, …, *N*_*θ*_ − 1} and a feature matrix **F** ∈ ℝ^*T* ×*d*^ whose row **f**_*t*_ is the activity of the *d* features at frame *t*. Three feature sets were decoded: the **communities** (*d* = number of detected communities, using the community activity signals from the canonical run), the **full neuron population** (*d* = number of neurons; an upper reference), and a **position-shuffled control** (chance, defined below).

#### Rate estimation

Calcium-derived activity is sparse per frame, so each feature was temporally smoothed with a Gaussian kernel *G*_*σ*_ (*σ* = 10 frames) to estimate an instantaneous rate,

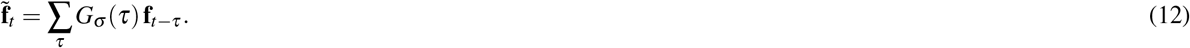

#### Cross-validated template decoder

The frame indices {1, …, *T*} were split into five *contiguous* blocks ℱ_1_, …, ℱ_5_ (temporally separated train/test partitions, which limits leakage from temporal autocorrelation). For fold *k*, the training set is *T*_*k*_ = _*l* ≠*k*_ ℱ_*l*_ and the test set is ℱ_*k*_. For each position bin *b*, a template is the mean training activity in that bin,

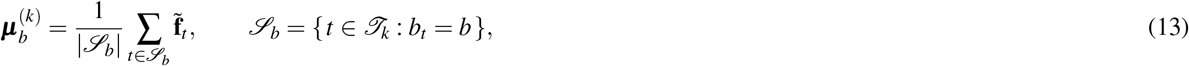

with bins unvisited during training excluded. Each held-out frame is assigned the bin whose template is most

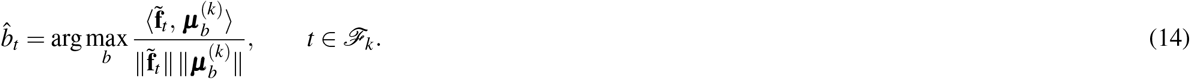

#### Error and chance

Decoding error is the circular distance between the decoded and true bin, in degrees,

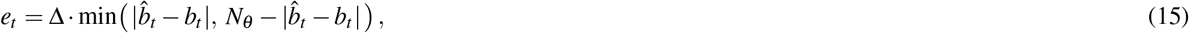

summarised per session by its median 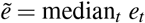. For a decoder with no positional information the expected error is the mean circular distance between independent uniform bins, ≈ 90°. In addition to this analytic chance level we computed an empirical control by circularly shifting the position sequence by ⌊*T/*3⌋ frames, *b*_*t*_ → *b*_(*t*+ ⌊*T/*3⌋) mod *T*_, before both template construction and evaluation; this preserves the marginal statistics of activity and position while destroying their temporal correspondence. Community decoding error was well below both chance levels in every group, though above (i.e. coarser than) that of the full neuron population, quantifying the compression–fidelity trade-off of the community code (Supplementary Fig. 4).

**Supplementary Fig. 4.**
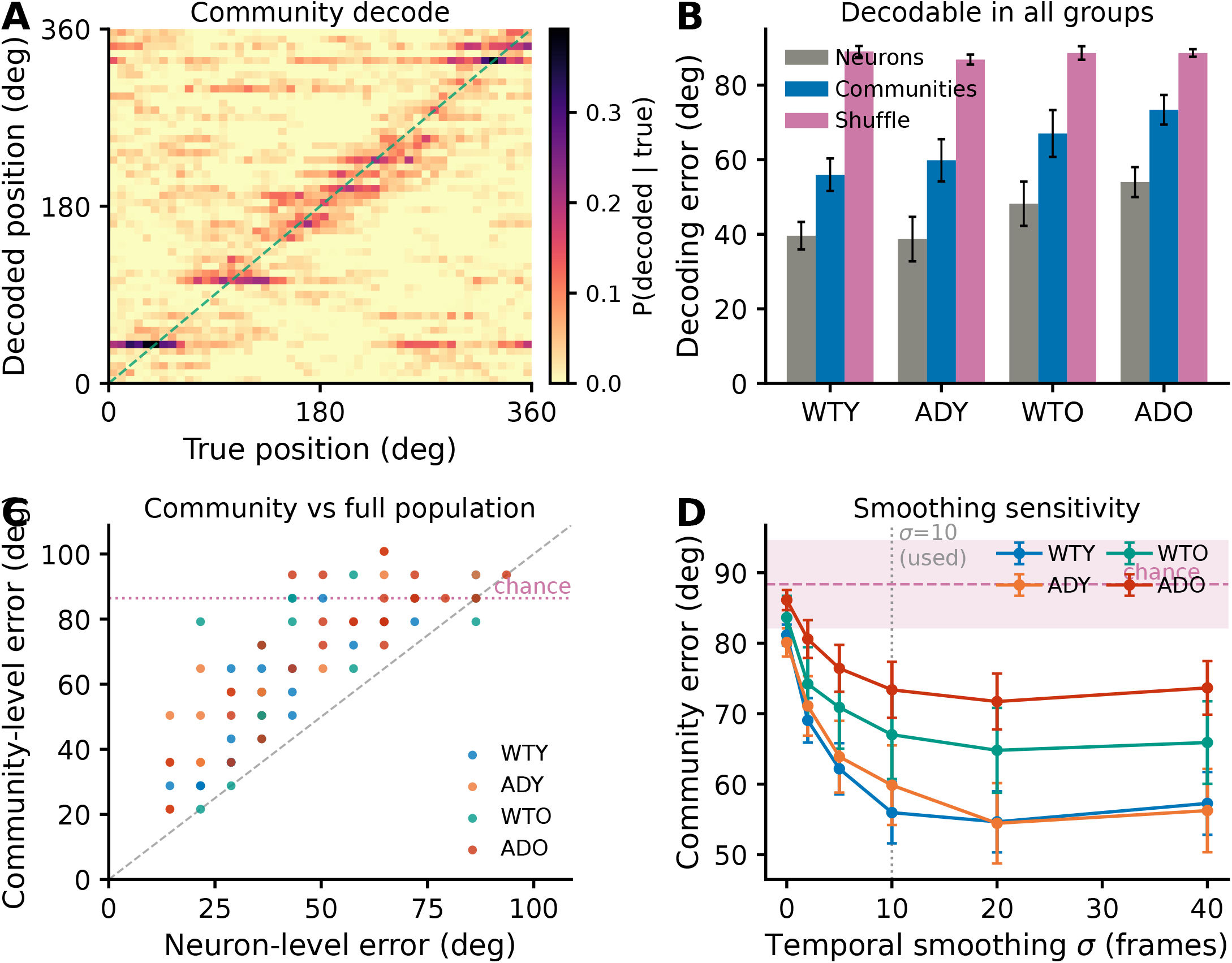
The animal’s track position is decodable from community-level activity. A cross-validated population-vector decoder reconstructs position on the circular track from the activity of the detected communities, quantifying the decodability illustrated schematically in Fig. 1C. For each session, neural data were preprocessed exactly as for the PID analysis (speed-filtered, global *k* = 5 discretisation, position binned into 50 equal-angle bins; Methods); per position bin a template activity vector was built from the training timepoints and each held-out timepoint assigned the bin with the most similar (cosine) template, under 5-fold contiguous cross-validation (temporally separated train/test blocks). Error is the circular distance between decoded and true position (degrees; chance ≈ 90°). **A**, Decoded versus true position for a representative healthy (WTY) session, chosen as the session whose community decoding error is closest to the WTY median (36°), shown as the confusion matrix *P*(decoded | true) over held-out timepoints; probability concentrates on the diagonal (green), i.e. the community code tracks position. **B**, Decoding error per group for three feature sets: the full neuron population (grey, an upper reference), the detected communities (blue), and a position-shuffled control (pink, chance); bars are mean ± s.e.m. across sessions. Community decoding error is well below chance in every group. **C**, Per-session community-level versus neuron-level decoding error (dashed, identity; dotted, chance). Every session lies below chance and above the identity line: the community representation (a ~ 10-fold dimensionality reduction from the full population) retains substantial, though coarser, positional information. **D**, Sensitivity of the community decode to the temporal-smoothing scale *σ* (all 77 sessions, per group; mean ± s.e.m.). Without smoothing (*σ* = 0) the per-frame decode sits at chance: calcium signals are slow, so a single frame carries little instantaneous rate information. The error then falls steeply and plateaus by *σ* ≈ 10 frames (~ 1 s), the value used throughout (dotted), in every group. The decodability is therefore a property of the community code, not an artefact of a finely-tuned smoothing choice.

### Supplementary Methods: ground-truth simulation model and logic-gate benchmarks

#### Rationale and overview of the validation

Redundancy and synergy are estimator-dependent quantities, so we wanted to establish that the atoms reported for the real recordings are properties of the neural code rather than of the estimator, the amplitude discretisation or the sample size. We therefore validated the MMI–PID readout on synthetic data whose decomposition is known analytically, using constructions on which the correct answer is pure redundancy or pure synergy, generated by the same simulation code used throughout the project and passed through the identical analysis pipeline applied to the real recordings (Supplementary Fig. 5A,B). The construction has three stages: (i) a behavioural trajectory and a binary spatial target, (ii) a Poisson spiking model in which each neuron is driven by a binary latent state, and (iii) two logic gates over those latent states that set the ground-truth atom (redundancy or synergy).

#### Trajectory and spatial target

We simulated *N*_*T*_ = 9,000 samples at Δ*t* = 0.1 s of angular position *θ*_*t*_ ∈ [0, 360)° on a circular track of radius *r* = 200. At each step the instantaneous speed is drawn from an exponential distribution, 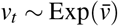 with mean 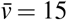, and the angle is advanced by the corresponding angular increment,

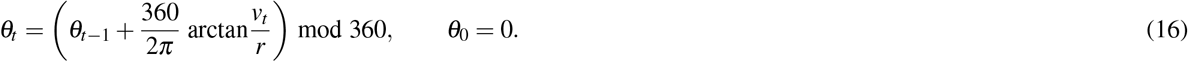

The decoding target is the balanced one-bit “track half” variable

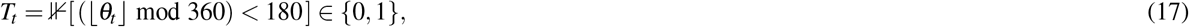

so that the maximal recoverable information about *T* is exactly 1 bit.

#### Poisson spiking model

Each simulated neuron carries a binary latent drive *d*_*t*_ ∈ {0, 1} and emits Poisson spike-amplitude counts whose rate is gated by that drive,

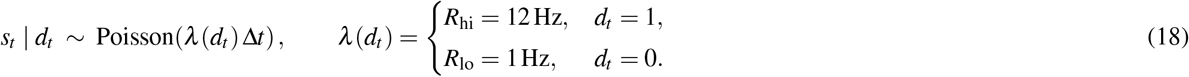

The resulting amplitude matrix is discretised with the same global equal-occupancy *k* = 5 amplitude binning used for the real deconvolved amplitudes (Methods; Supplementary Fig. 1), and the target *T* is treated as a *k*_pos_ = 2 state variable. PID atoms are then computed with the identical NSB–MMI estimator (Methods), averaged over 40 independently simulated neuron pairs per condition (mean ± s.e.m.). Information structure is therefore imposed entirely through the latent drives; the two gates below specify how each pair’s drives (*d*^1^, *d*^2^) relate to the target *T*.

#### COPY gate (ground-truth redundancy)

Neuron 1 faithfully encodes the target, 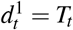 Neuron 2 copies the target with fidelity *c* ∈ [0, 1]: on a fraction *c* of timepoints its drive equals the target, otherwise it is an independent fair coin,

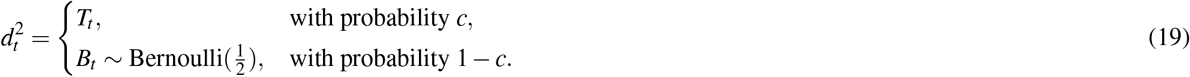

As *c* → 1 the two neurons carry the *same* bit, so both single-source informations approach 1 bit and

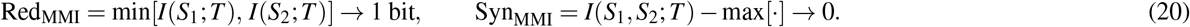

A correct estimator should attribute the copied information to redundancy and leave synergy near zero across the fidelity sweep.

#### XOR gate (ground-truth synergy)

Each pair shares a hidden fair-coin latent *h*_*t*_ ~ Bernoulli 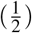, and the drives are

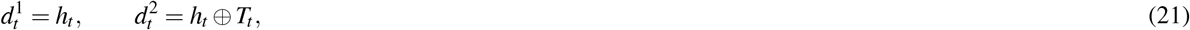

where ⊗ is addition modulo 2 (exclusive-or). Because *h* is an unbiased coin independent of *T*, each neuron alone is uninformative, *I*(*S*_1_; *T*) = *I*(*S*_2_; *T*) = 0, yet the joint state determines the target exactly, *I*(*S*_1_, *S*_2_; *T*) = 1 bit, giving

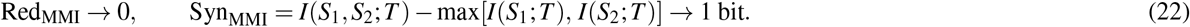

This is the canonical pure-synergy construction^46,92^. We sweep the fraction *f* ∈ [0, 1] of XOR-coded pairs (the remainder being fully copy-coded, *d*^1^ = *d*^2^ = *T*); as *f* rises the population atom shifts continuously from redundancy-to synergy-dominated, and the crossover confirms that the estimator recovers the synergy axis where it is present.

**Supplementary Fig. 5.**
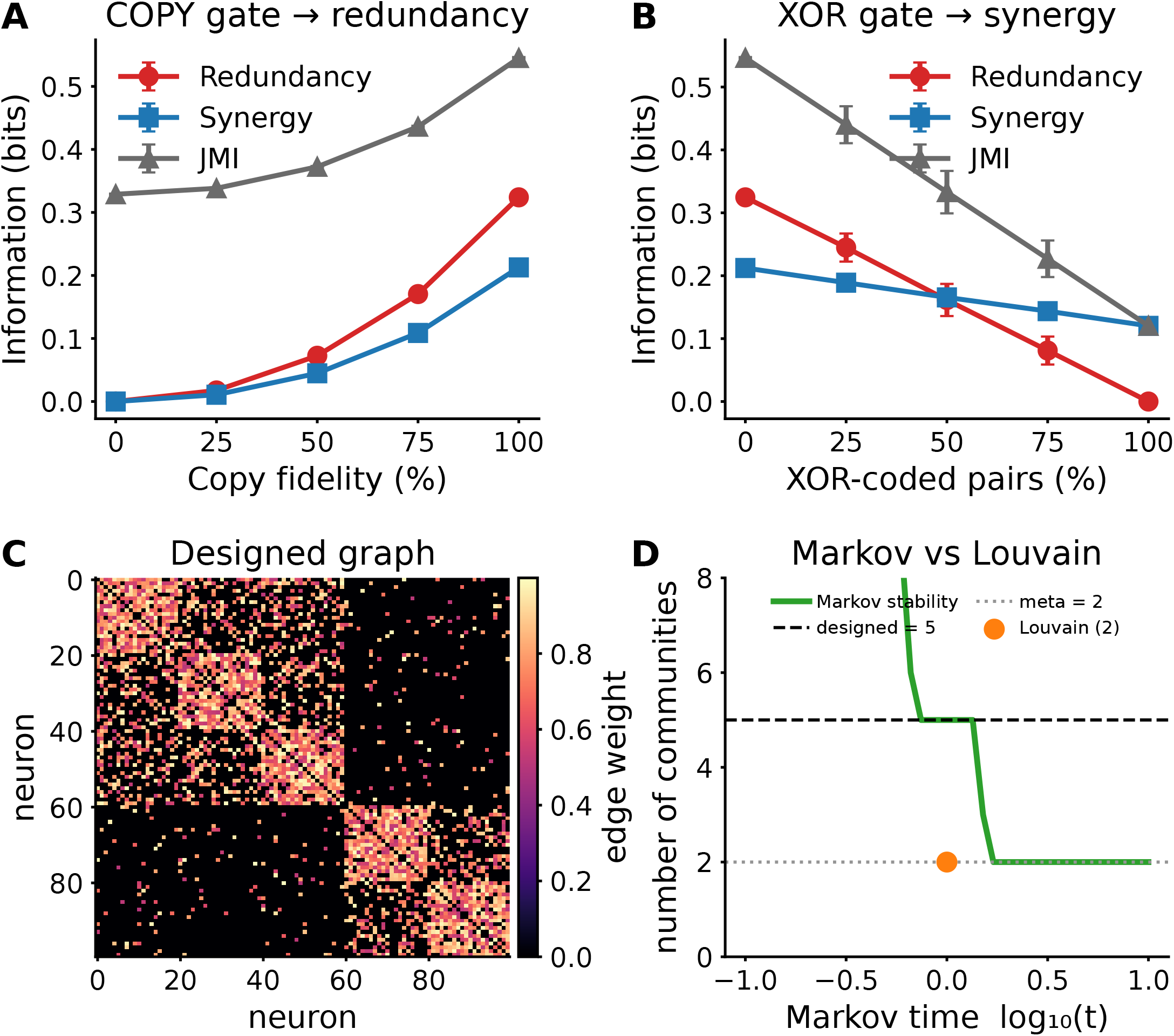
Ground-truth validation of the MMI–PID and Markov stability pipeline using canonical information-theoretic benchmarks. Simulated Poisson spike trains, driven by binary latent states and discretised through the same global equal-occupancy *k* = 5 amplitude binning used for the real data (Methods), were passed through the paper’s MMI–PID estimator. Logic-gate constructions set the ground-truth decomposition^46,92^; all atoms are in bits, averaged over 40 neuron pairs per sweep level (mean ± s.e.m.). **A**, COPY gate (redundancy). Neuron 1 encodes a binary target (track half) and neuron 2 copies that target with copy-fidelity 0–100%. As fidelity rises the two neurons carry the same bit: redundancy (Π_red_, red) increases and remains the dominant atom, synergy (Π_syn_, blue) stays low, and joint mutual information (JMI, grey) tracks the shared information. The estimator correctly attributes copied information to redundancy. **B**, XOR gate (synergy). Pairs are morphed from copy-coded (redundant) to XOR-coded via a shared latent state *h* (drive_1_ = *h*, drive_2_ = *h* ⊗ target), so that neither neuron alone carries information about the target but their joint state determines it. As the XOR-coded fraction rises, redundancy collapses to zero and synergy becomes the dominant atom, the signature of a synergistic code. The estimator recovers synergy where it is present. **C**, Designed hierarchical graph used as community-detection ground-truth: a weighted stochastic block model with five fine communities nested within two meta-communities (adjacency matrix; colour = edge weight). **D**, Markov stability versus Louvain on that graph. Markov-stability community detection (green) resolves the multiscale structure across Markov time, a five-community partition at short Markov times descending through a two-community (meta) plateau at longer times, whereas single-resolution Louvain (orange) returns a single partition matching neither designed scale. Markov stability is therefore used throughout the paper in place of Louvain.

**Supplementary Fig. 6.**
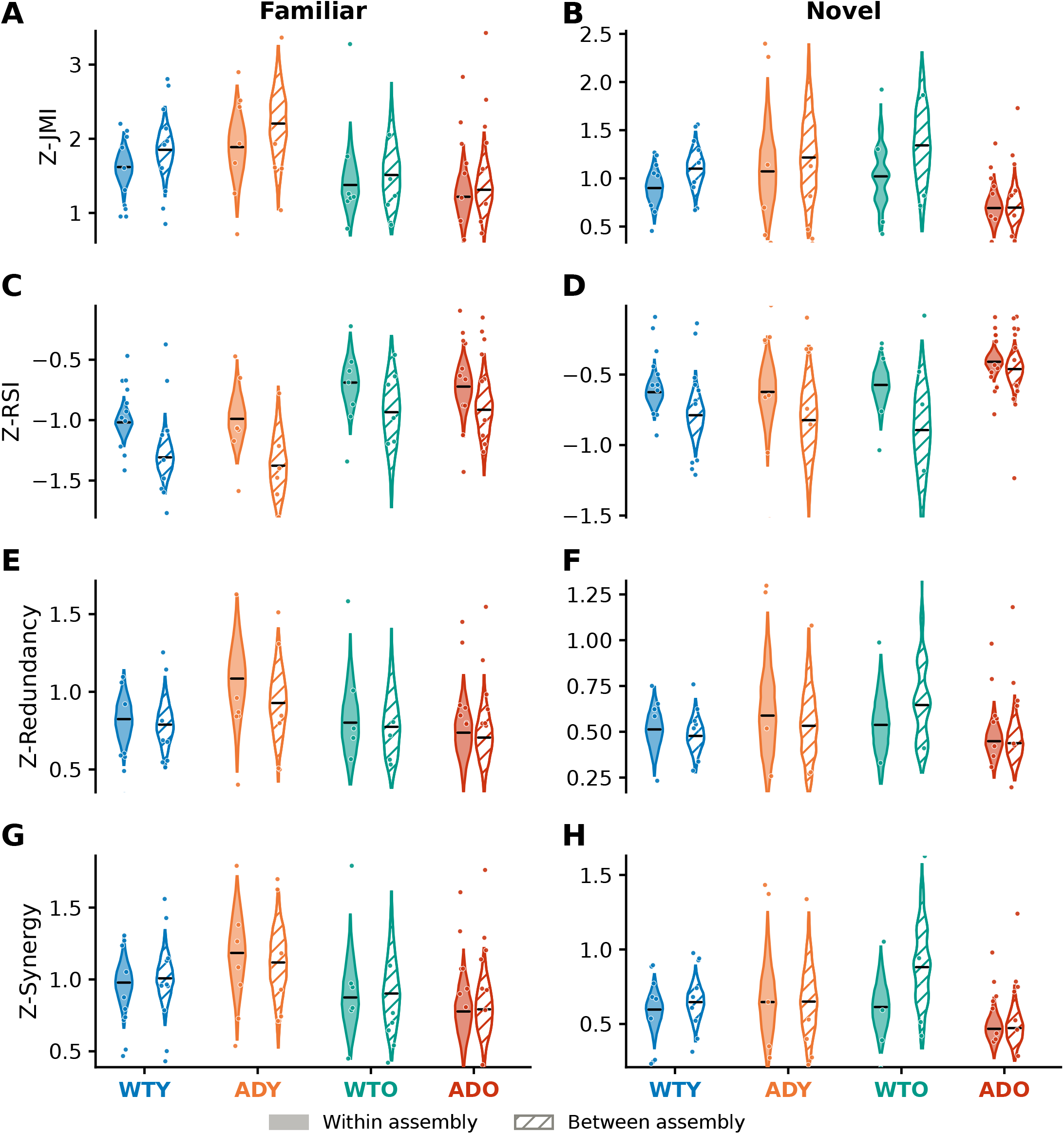
Full PID atom distributions across all groups and both environments. Violin plots (hierarchical bootstrap, *B* = 5,000) of all four *z*-scored PID atoms for within-assembly (solid) and between-assembly (hatched) neuron pairs, in all four groups. Rows are the four PID atoms (JMI, RSI, Redundancy, Synergy); the left column (**A,C,E,G**) is the familiar environment and the right column (**B,D,F,H**) the novel environment. Black brackets mark within- vs. between-assembly contrasts significant at *P <* 0.05 (two-sided hierarchical bootstrap). The interpretable quantity in each panel is the *within-vs-between contrast within a group*: in WTY and ADY, between-assembly pairs carry higher JMI and more negative RSI than within-assembly pairs, and this contrast is attenuated in WTO and ADO, consistent with the three-way collapse (Fig. 3). The synergy contrast (**G,H**) shows the four-way topology × environment × genotype × age interaction (Fig. 4C). Absolute violin heights are *not* directly comparable across groups, because each group’s *z*-score is divided by its own surrogate-distribution width; group differences are read from the within-vs-between contrast and from the LMM, not from raw violin position. Colours: WTY (blue), ADY (orange), WTO (teal), ADO (red); WTO novel *n* = 5 sessions.

### Supplementary Methods: choosing the statistical model at each analysis level

#### Target of estimation

Every headline claim in this paper is an *interaction* rather than a single group difference: whether the effect of the amyloid genotype depends on age (genotype × age) and, at the pair level, whether that dependence itself differs between within- and between-assembly pairs (topology × genotype × age). The estimand is a difference of differences. A two-sample *t*-test estimates one difference of means and contains no parameter for it (Supplementary Fig. 7C), so at both analysis levels we fit a linear model carrying the full set of interaction terms. The levels then differ only in (i) whether that linear model needs a random effect and (ii) how its coefficients are estimated. This section states exactly what was fitted at each level and why.

#### Data structure at each analysis level

The pairwise PID analyses (Figs. 2–4) have one observation per *neuron pair*: 814,031 familiar-environment pairs distributed over 40 cases (animal ×session units), a mean of 20,351 pairs per case. The network and community analyses (Figs. 5, 6) are aggregated: the structural metric table holds exactly one row per session × environment (77 rows, 40 familiar and 37 novel, each with its own case label), and the integrated emergence scores likewise one value per session (*N* = 77; the communities are the *parts* entering Ψ, Δ and the O-information, not separate observations). There is therefore massive within-unit replication at the pair level and none at the aggregated level, which is what dictates the two model classes.

#### Pair level: model specification

We fit the full-factorial linear mixed-effects model value ~ *T* × *E* × *G* × *A* + (1 | case) (topology *T*, environment *E*, genotype *G*, age *A*; maximum likelihood) to the *z*-scored pair-level metrics, by balanced subsampling of pairs within case (Methods, “Statistics and Reproducibility”; Supplementary Fig. 9). Its *fixed effects* are one coefficient per term of the factorial design, the quantities of interest, and its *random intercept* (1 | case) is a per-case offset that absorbs each animal’s baseline level of the metric.

#### Pair level: necessity of the random intercept

Neuron pairs from the same case are not independent: they share an animal, a field of view and a functional network. We quantified this by a one-way random-effects variance decomposition on case, giving an intraclass correlation 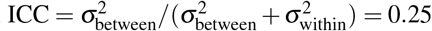 for RSI and 0.17 for redundancy: a quarter of the pair-to-pair variance in RSI is between cases, not within them. Under the standard design-effect correction, 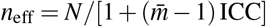 with *N* = 814,031 pairs and 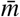 pairs per case, those pairs carry the information of only *n*_eff_ ≈ 159 (RSI) to 237 (redundancy) independent units (Supplementary Fig. 7A). A pooled *t*-test or ordinary least squares would compute the standard error from *N* itself (“pseudoreplication”^99^), inflating it more than a thousand-fold relative to the ~ 40 cases that actually vary; the random intercept ties the fixed-effect standard error to the case count instead.

#### Aggregated level: non-identifiability of the random intercept

Each unit here contributes exactly one value, so there is no within-unit replication from which a between-unit variance could be estimated. Fitting the mixed model anyway (same fixed structure, grouping on the session × environment label) makes the random-intercept and residual variances perfectly confounded: the fit returns the non-informative equal split, a random-effect share of 0.500 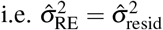 exactly), and reduces to the single-level fit (Supplementary Fig. 7B). We therefore drop the random effect and fit a single-level linear model with the same interaction structure, with environment entering as a fixed effect rather than as a repeated measure.

#### Aggregated level: rationale for robust (Huber) estimation

The fitted model is metric ~ (Genotype + Age + Novel + *n*_communities,c_)^3^, estimated with the Huber M-estimator (Huber weighting, tuning constant *c* = 1.345, iteratively reweighted least squares; full specification in Methods, “Network Structural Analysis” and “Community-Level Emergence Measures”). Robust estimation is used because the sample is small (*N* = 77 sessions, 5–13 per group and environment) and the null-corrected structural and emergence metrics are heavy-tailed with a few high-leverage sessions: ordinary least squares gives every session weight *w* = 1, whereas the Huber weight falls as *c/* |*r*| once a session’s standardised residual |*r*| exceeds *c*, so no single session can drive a coefficient. Across the eight aggregated network metrics, 124 of the 616 session fits (8 metrics × 77 sessions) are down-weighted, the strongest to *w* = 0.10 (Supplementary Fig. 7D).

#### Comparison of robust and ordinary least-squares fits

Every network metric was also refit by ordinary least squares as a check. Each metric is standardised to unit variance first, as in the supplementary network forest (Supplementary Fig. 11), so that the subtract-only modularity endpoint and the seven *z*-scored metrics share one coefficient scale. Over the 24 coefficients (8 metrics × genotype, age and genotype × age) the two estimators agree closely (*r* = 0.96; Supplementary Fig. 7E), so the reported effects are not an artefact of the estimator, and 18 of the 24 are pulled *toward* zero by the robust fit: the choice is the conservative one, tempering rather than creating effects.

#### Scope and limitations of the model-selection argument

The argument above is structural. It justifies the model *class* (interaction terms throughout; a random intercept only where it is identifiable) and the *estimator* (robust where the sample is small and heavy-tailed); it is deliberately not a *P*-value contest between a *t*-test and the mixed model. In particular, the standard errors reported for the LMM coefficients are across-subsample standard deviations and quantify the stability of each estimate under pair subsampling, not between-animal uncertainty (Supplementary Methods, “balanced subsampling of the pair-level LMM”). Animal-level uncertainty is supplied separately by the hierarchical bootstrap (Supplementary Methods, “hierarchical (Saravanan) bootstrap for pair-level metrics”; Supplementary Fig. 8), which yields the confidence intervals reported throughout the pairwise PID figures.

**Supplementary Table 1.** Statistical model used for each analysis. LMM, linear mixed-effects model; HB, hierarchical bootstrap; RLM, robust linear model (Huber M-estimator). *T*, topology; *E*, environment; *G*, genotype; *A*, age; *n*_c_, mean-centred per-session community count. A session is one animal in one environment and is the unit of replication throughout; it is the unit termed a *case* in the figure labels and analysis code.

| Analysis | Observation | Model | Reported |
| --- | --- | --- | --- |
| Pairwise PID atoms: JMI, RSI, redundancy, synergy (Figs. 2–4) | neuron pair | LMM, full factorial $T \times E \times G \times A$, session random effect | Mean $\hat{\beta}$ over 100 balanced subsamples; Wald $z$ |
| Condition-specific PID contrasts, including WTY vs. ADO (Figs. 2–4) | neuron pair | Reduced LMM, single fixed effect within the conditioning stratum, session random effect | Mean $\hat{\beta}$; Wald $z$ |
| Plotted PID distributions and their intervals (Figs. 2–4) | session | HB, $B = 10,000$, two-level resample | 95% percentile CI; $P = 2 \min(\cdot)$ |
| Network structure, eight null-corrected metrics (Fig. 5) | session | RLM, metric $\sim (G + A + \text{Novel} + n_c)^3$ | $\hat{\beta}$, 95% CI; Wald $z$ |
| Community-level emergence $\Psi$, $\Delta$, O-information (Fig. 6) | session | RLM, same factorial form | $\hat{\beta}$, 95% CI; Wald $z$ |

**Supplementary Fig. 7.**
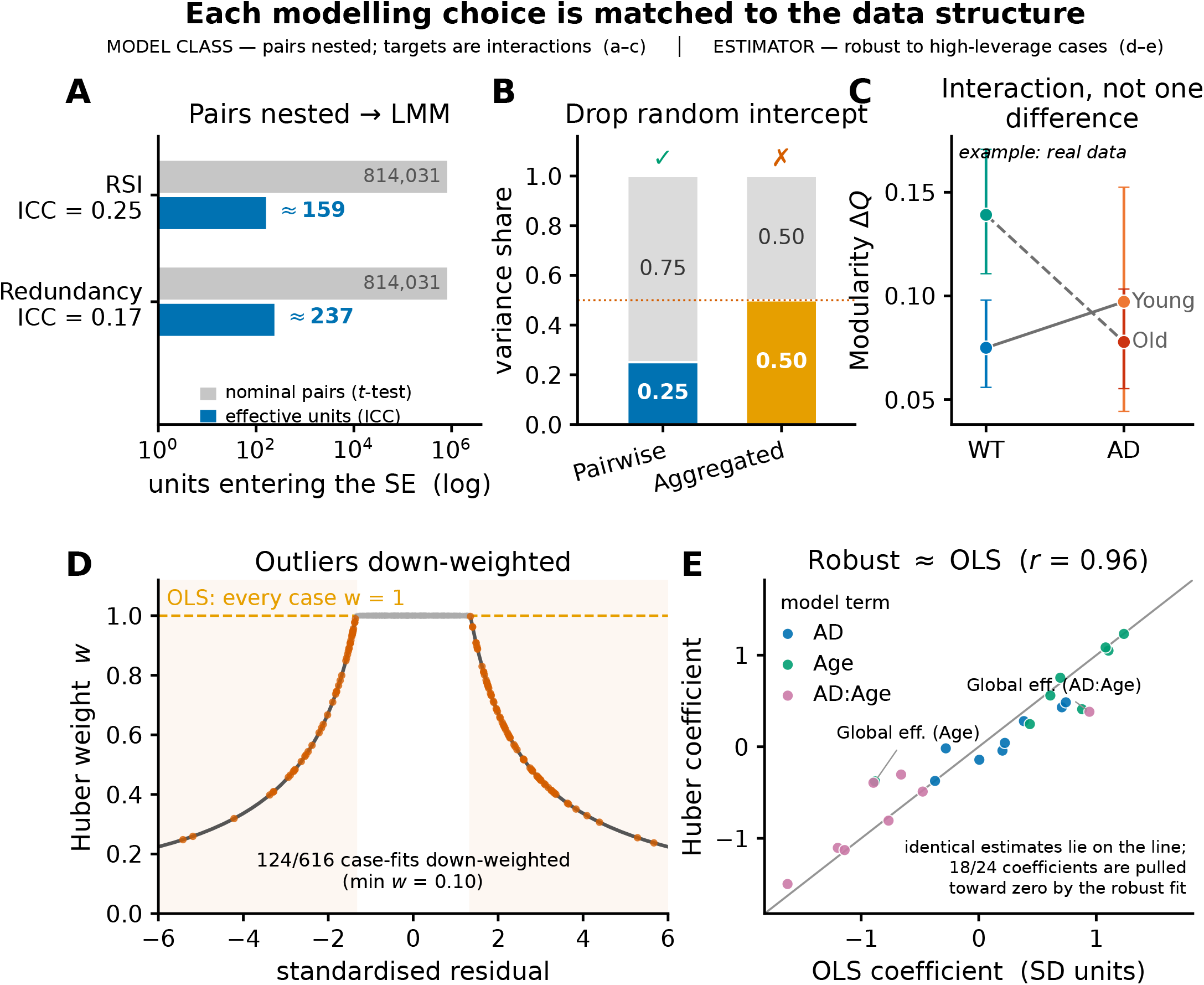
The statistical model is matched to the data structure at each analysis level. Rationale for the linear mixed-effects model used for the pairwise PID metrics and the robust single-level linear model used for the network and emergence metrics (Supplementary Methods); resampling *mechanics* are covered in Supplementary Figs. 8 and 9. **A**, Pair-level are nested: neuron pairs within a case are correlated. The intraclass correlation, the share of the pair-to-pair variance that lies *between* cases rather than within them, is ICC = 0.25 for RSI and 0.17 for redundancy. Applying the design-effect correction 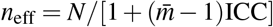, the 814,031 familiar-environment pairs (grey bars, the number a pooled *t*-test would count) carry the information of only ≈ 159–237 independent units (blue bars); note the logarithmic axis. The random intercept (1 | case) ties the fixed-effect standard error to the ~ 40 cases. **B**, The random-intercept (between-case) variance is estimable only with replication within a case. Pairwise it is well defined (share = ICC ≈ 0.25; keep the mixed model); with one value per case it is confounded with the residual and the fit returns the non-informative 0.50 split (red dotted line), so the random effect is dropped for a single-level model. Blue/orange, between-case share; grey, within-case (residual) share. **C**, The scientific target is an interaction. Worked example using the modularity endpoint of Fig. 5 (Δ*Q*^str^, familiar environment): group means ± 95% CI from 5,000 case-level bootstrap resamples (*n* = 11 WTY, 8 ADY, 8 WTO, 13 ADO sessions). Genotype raises Δ*Q* in young animals but lowers it in old ones (non-parallel lines), a difference of differences; a two-sample *t*-test estimates one difference and has no parameter for this class of effect. Colours: WTY blue, ADY orange, WTO teal, ADO red. **D**, The Huber M-estimator down-weights high-leverage cases: weight *w* against a case’s standardised residual *r* (solid curve, *w* = 1 for |*r*| ≤ *c* = 1.345 and *w* = *c/* |*r*| beyond it; OLS fixes *w* = 1, dashed), with per-case residuals from all eight network metrics overlaid. Across the 8 × 77 = 616 session fits, 124 are down-weighted, the strongest to *w* = 0.10. **E**, Ordinary (OLS) versus robust (Huber) coefficients for the genotype, age and genotype × age terms of all eight network metrics, each standardised to unit variance (24 coefficients, SD units): they lie on the identity line (*r* = 0.96), while 18*/*24 are pulled toward zero by the robust fit. The same estimator is used for the emergence metrics (Fig. 6).

### Supplementary Methods: hierarchical (Saravanan) bootstrap for pair-level metrics

#### Rationale and overview of the resampling scheme

We want confidence intervals whose width reflects the number of *animals* we recorded, not the number of neuron pairs those animals happen to yield, while still using the within-animal data rather than collapsing each session to one number. We therefore resample at both levels, as set out below.

Pair-level PID metrics are not independent observations: the several hundred to several tens of thousands of neuron pairs in a session all derive from the same animal, field of view and functional network, and pair counts vary more than hundred-fold across cases (familiar environment: 561 pairs in the smallest case to 69,006 in the largest; per-group medians 9.8k–33k). Treating pairs as independent therefore pseudoreplicates the data^99^, while collapsing each case to a single mean discards most of the within-animal signal and sacrifices power^100^. We instead used the hierarchical bootstrap of Saravanan et al.^100^, which respects the three nested levels of the dataset: experimental group, case (animal × session) and neuron pair. Each of *B* = 10,000 iterations performs a two-step resample: (i) cases are drawn with replacement *within* each group, then (ii) neuron pairs are drawn with replacement *within* each selected case; the resampled pairs are reduced to a per-case mean and the iteration statistic is the grand mean over those per-case means (Methods, “Statistics and Reproducibility”). Resampling at the case level propagates between-animal variability into every confidence interval, so that significance reflects the number of animals rather than the number of pairs. The calibration of this scheme, the pair-level intraclass correlation and effective sample size, the false-positive inflation under naive pair-level resampling, and the leave-one-animal-out stability of the headline interaction are quantified in Supplementary Fig. 8.

**Supplementary Fig. 8.**
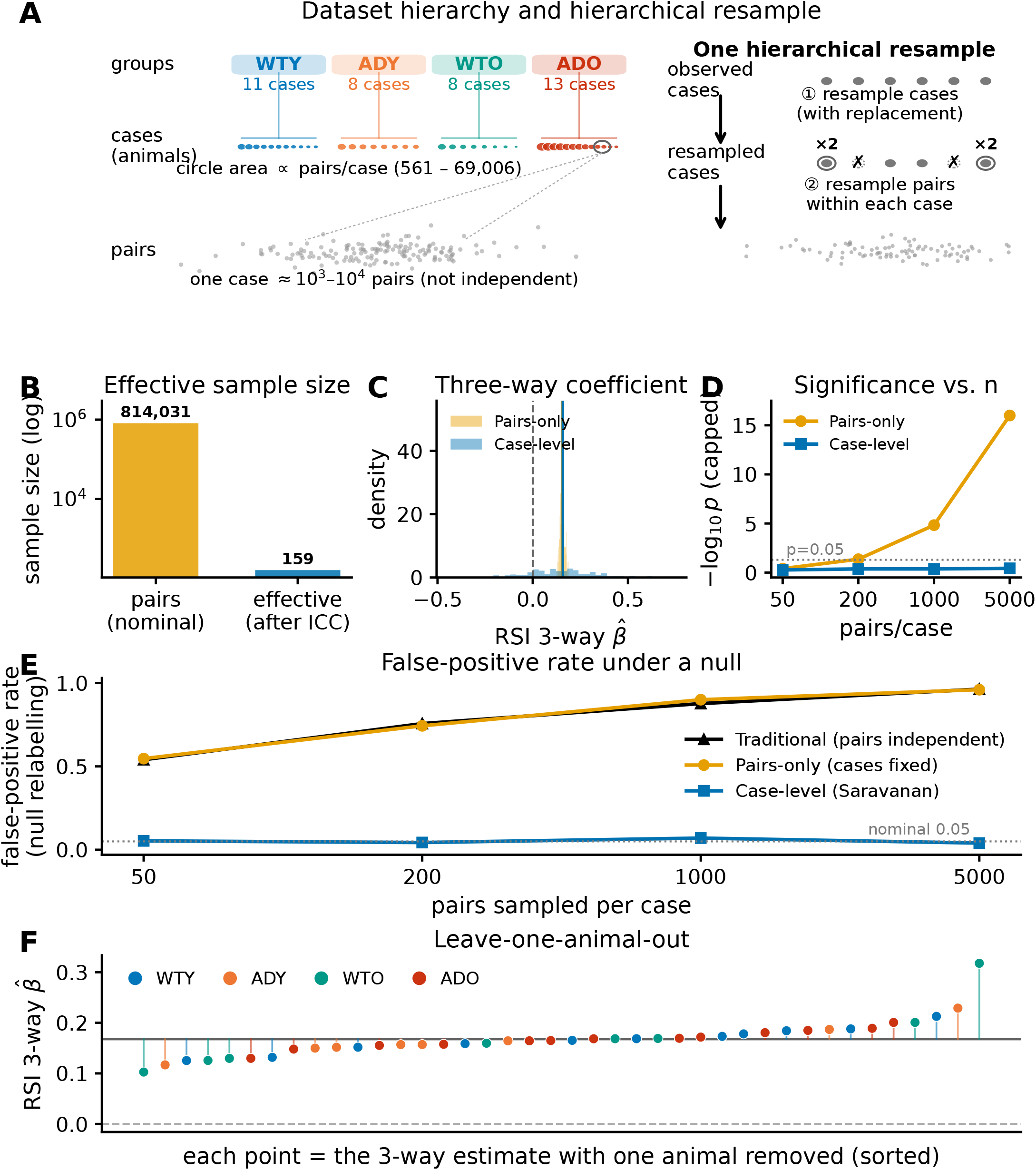
The pair-level analysis requires a hierarchical (case-level) bootstrap. Justification for the animal-level resampling used throughout (Methods, “Statistics and Reproducibility”). **A**, Data hierarchy and the two-step resample: the 814,031 familiar-environment neuron pairs are nested in 40 cases (animal × session units) within four groups; each bootstrap iteration first resamples cases with replacement (some doubled, some dropped), then resamples pairs within ≈each drawn case. **B**, The pair-level intraclass correlation (ICC = 0.25) reduces the nominal 814,031 pairs to an effective sample size of only *n* ≈ 159 independent units. **C**, Bootstrap distribution of a representative three-way interaction coefficient (RSI, topology × genotype × age): resampling pairs only (cases fixed; orange) is artificially narrow, whereas the case-level resample (blue) reflects the true between-case uncertainty at *n* = 8–13 cases (animal–session units) per group. **D**, Under pairs-only resampling the same coefficient’s apparent significance ( −log_10_ *P*) grows without bound as more pairs are sampled per case (reaching *P* → 0), the diagnostic signature of pseudoreplication, whereas the case-level *P* is stable and independent of sampling depth. **E**, False-positive rate under random relabelling of animals (a true null): treating pairs as independent (black) or resampling pairs only (orange) inflates the false-positive rate to 0.5–1.0 and worsens with more pairs, whereas the case-level bootstrap (blue) stays calibrated at the nominal *α* = 0.05 at all sampling depths. **F**, Leave-one-animal-out: recomputing the three-way estimate with each single animal removed shifts it appreciably (range 0.10–0.32), confirming that the estimate depends on a small number of animals and that animal-level resampling is required for valid inference. Colours: WTY blue, ADY orange, WTO teal, ADO red.

### Supplementary Methods: balanced subsampling of the pair-level LMM

#### Rationale and overview of the subsampling scheme

We want a fixed-effect estimate that is not dominated by the handful of sessions contributing the most neuron pairs, and a fit that converges reliably; we do *not* claim animal-level uncertainty from this procedure. We therefore fit the model repeatedly to case-balanced subsamples of the pairs and summarise the estimates across those fits, as set out below.

The full-factorial linear mixed-effects models (Methods, “Statistics and Reproducibility”) are fit to individual neuron pairs, but the number of pairs per case varies more than hundred-fold (familiar environment: 561 pairs in the smallest case to 69,006 in the largest). A single fit to all ~1.6 × 10^6^ pooled pairs is therefore dominated by a handful of large sessions, and at that size the random-intercept optimisation is prone to convergence failure. To balance case contributions and keep each fit tractable, the model is fit by *balanced subsampling*: in each of 100 iterations we draw at most 5,000 within-assembly and 5,000 between-assembly pairs from every case (all available pairs when fewer than 5,000; subsample seed = 42 + *i* for iteration *i*), refit the full-factorial model value ~ *T* × *E* × *G* × *A* + (1|case) by maximum likelihood, and store the 16 fixed-effect coefficients. Each reported coefficient is the mean of the per-iteration estimates and its standard error the standard deviation across the 100 iterations, from which the Wald *z* and *P* follow (Methods, “Statistics and Reproducibility”); the *P*-values themselves are not averaged.

Supplementary Fig. 9 verifies that this estimator is well behaved. The per-iteration coefficient distributions for the headline terms are narrow and, for the topology and topology × genotype × age contrasts, well separated from zero (panel C); the running mean of each coefficient plateaus within ~ 20–30 iterations, so 100 iterations is more than sufficient for a stable estimate (panel D); the per-iteration estimates are approximately Gaussian, which justifies the normal-approximation Wald step (panel E); and the coefficients are essentially unchanged when the per-case cap is varied from 1,000 to 7,500 pairs per type, so the 5,000 cap is not a load-bearing choice (panel F).

The across-iteration standard deviation quantifies the *stability* of each fixed-effect estimate under pair subsampling; it is not a between-animal standard error, because cases are never resampled in this procedure. Animal-level uncertainty is supplied separately by the hierarchical bootstrap (Supplementary Methods, “hierarchical (Saravanan) bootstrap for pair-level metrics”; Supplementary Fig. 8), which yields the confidence intervals and violin distributions reported throughout the pairwise PID figures.

**Supplementary Fig. 9.**
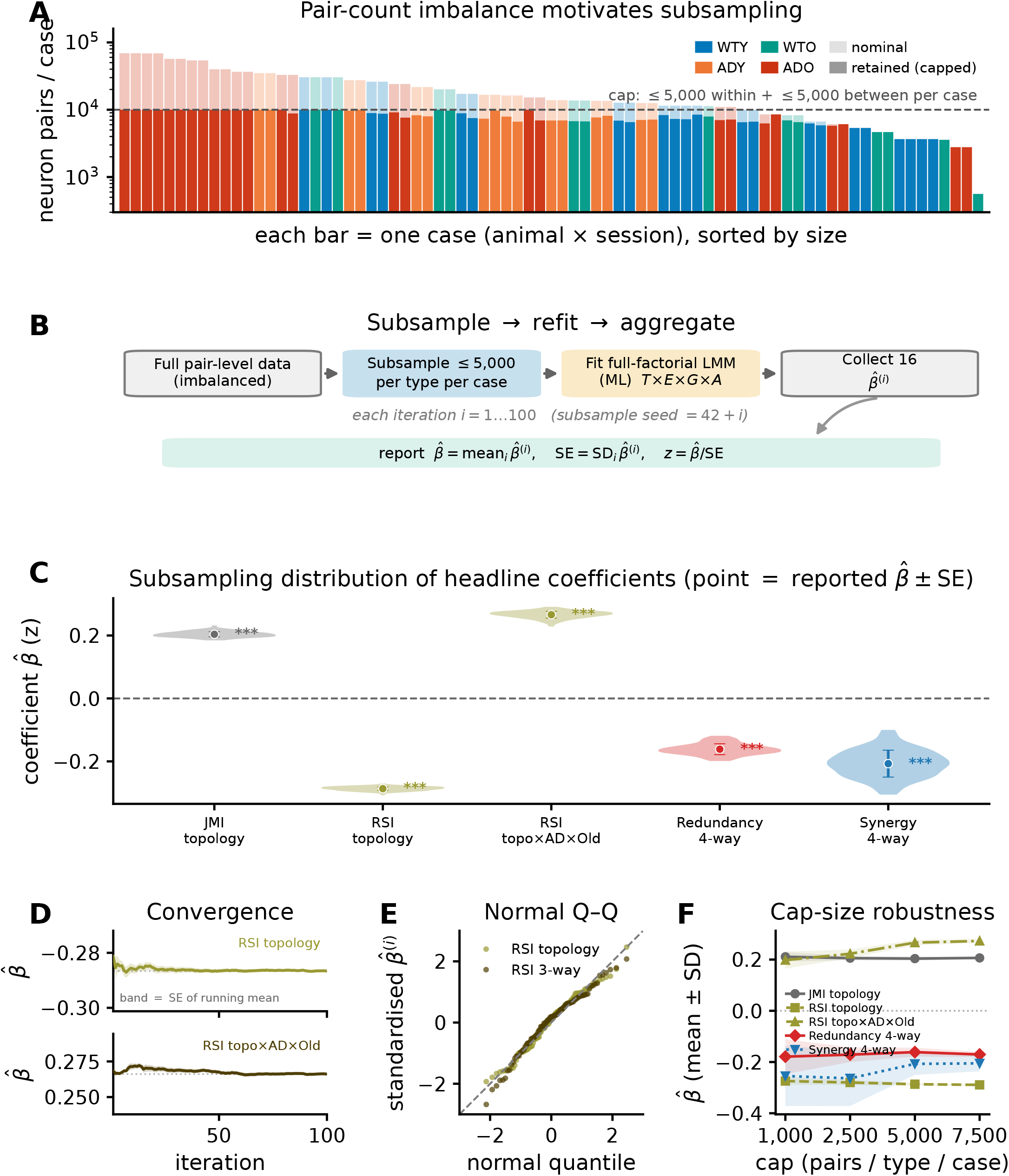
The pair-level linear mixed-effects model is fit by balanced subsampling; its coefficient distribution is stable, convergent and approximately normal. Justification for the subsampling estimator used for the LMM fixed-effect tests throughout the pairwise PID figures (Methods, “Statistics and Reproducibility”); companion to the hierarchical bootstrap in Supplementary Fig. 8. **A**, Neuron-pair counts per case span 561–69,006 (over two orders of magnitude), so a single pooled fit would be dominated by a handful of large sessions. Each iteration therefore draws at most 5,000 within- and 5,000 between-assembly pairs per case (dark = retained, light = nominal), balancing case contributions and bounding the fit size (21*/*77 cases exceed the within-cap, 64*/*77 the between-cap). Colours: WTY blue, ADY orange, WTO teal, ADO red. **B**, The estimator. Each of 100 iterations subsamples the pairs, refits the full-factorial model value ~ *T* × *E* × *G* × *A* + (1 |case) by maximum likelihood, and stores the 16 fixed-effect coefficients; the reported 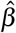 is the across-iteration mean, its standard error the across-iteration standard deviation, and 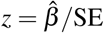. **C**, Subsampling distribution (violins, 100 iterations) of the five headline coefficients; the overlaid point is the reported 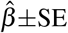 and stars the reported Wald *P* (^*^*P <* 0.05, ^**^*P <* 0.01, ^***^*P <* 0.001). The distributions are narrow and, for the topology and topology × genotype × age terms, well separated from zero. **D**, Per-term running mean of the two headline RSI coefficients, each on its own *y*-scale; the band is the standard error of the running mean, which narrows as 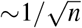. Both estimates are stable from the first few iterations (subsampling CV 2–4%) and the SE band flattens well before 100 iterations, so the iteration count is more than sufficient. **E**, Normal-probability (Q–Q) plot of the standardised per-iteration coefficients: points fall on the identity line, i.e. the across-iteration distribution is approximately Gaussian, supporting the Wald *z*/*P* step. **F**, The headline coefficients are essentially unchanged when the per-case cap is varied from 1,000 to 7,500 pairs per type (the SD narrows as more pairs are used, as expected), so the 5,000 cap is not a load-bearing choice. This subsampling quantifies the *stability* of the fixed-effect estimate under the pair-count imbalance in **A**; it is not a substitute for animal-level uncertainty, which is provided separately by the hierarchical bootstrap (Supplementary Fig. 8) underlying the plotted violins and confidence intervals.

**Supplementary Fig. 10.**
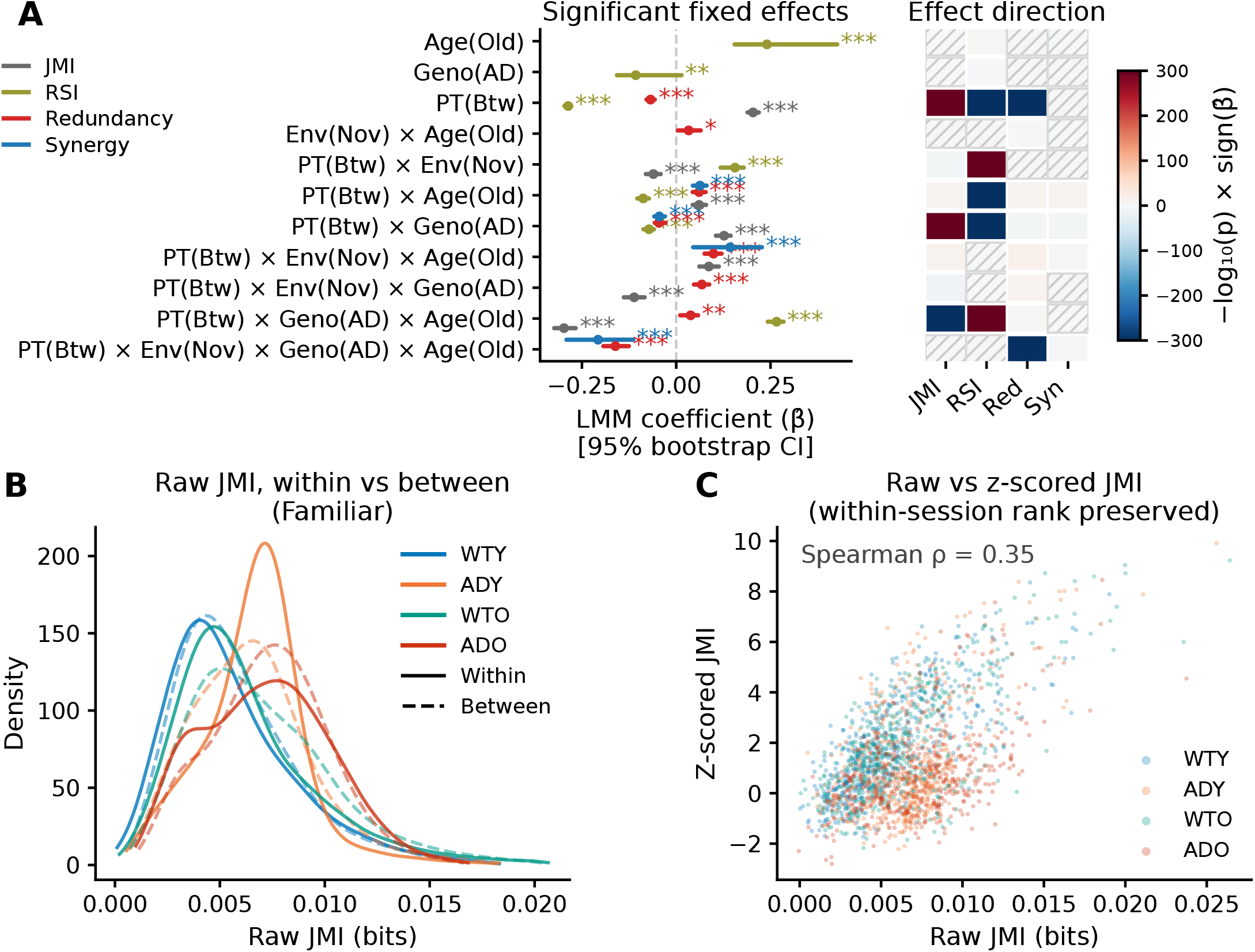
Linear mixed-effects model summary and raw-bits dissociation. **A**, Combined LMM summary. *Left*, forest plot of the significant fixed-effect coefficients from full-factorial LMMs fit to z-scored PID metrics (JMI grey, RSI olive, Redundancy red, Synergy blue) overlaid in one plot. Points indicate coefficient estimates 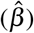, horizontal bars 95% LMM-subsampling interval; stars indicate *P <* 0.05 (^*^), *P <* 0.01 (^**^), *P <* 0.001 (^***^). *Right*, effect-direction heatmap sharing the forest’s term rows (rows aligned, term labels shown once on the left): signed −log_10_(*P*) × sign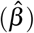 per term × metric (red = positive, blue = negative). Hatched cells denote terms that were estimated but did not reach significance (*P* ≥ 0.05); the heatmap therefore shows every term the forest omits. Cells for terms not estimable for a metric would be plain grey, and none occur here. The topology main effect (between vs. within assembly) and the Topology × genotype × age three-way interaction are the dominant terms for JMI and RSI. The four-way interaction (Topology × environment × genotype × age) is the dominant significant term for Synergy. **B**, Raw (unscaled) JMI density in bits, within-assembly (solid) vs. between-assembly (dashed) per group, familiar environment, showing the absolute information scale. ADO/ADY distributions are shifted toward higher raw values despite their surrogate-normalised deficit. **C**, Raw versus *z*-scored JMI across within-assembly pairs: *z*-scoring against per-pair circular-shift surrogates removes the absolute scale (and the hyperactivity-driven inflation of raw bits) while preserving within-session rank order, which is why the paper reports *z*-scored atoms. This raw/*z* dissociation is the basis for interpreting the coding deficit as a loss of spatial specificity rather than of absolute information (see Discussion).

### Supplementary Note: supporting analyses for the network structural results

Before null correction, the eight structural metrics carry no group structure, which is what motivates the configuration-model correction used throughout the structural analysis (Methods, “Configuration-Model Nulls (Strength-Preserving)”). Fitting the Huber model of Methods (“Network Structural Analysis”) to the raw metric scale, modularity, clustering and average path length showed no significant terms (all *P >* 0.05). Univariate Kruskal–Wallis tests on the familiar-only distributions agreed: modularity *P* = 0.657, path length *P* = 0.650, and clustering *P* = 0.009 with a post-hoc WTO–ADO shift (*P* = 0.019) that did not survive the multivariate model. Raw graph metrics are dominated by each backbone’s degree and edge-weight sequence, so a group difference in the organisation of the network is not separable from a group difference in its density on this scale; the strength-preserving null removes that dependence and is the basis of the primary structural results (Supplementary Fig. 11).

Three details of the null-corrected Huber fits (Methods, “Configuration-Model Nulls (Strength-Preserving)”) support the primary result but are not shown in Fig. 5 or Supplementary Fig. 11. First, the per-group Δ*Q*^str^ means (WTY 0.081, ADY 0.098, WTO 0.127, ADO 0.071) show directly that the age-driven modularity gain seen in wild-type is absent in the aged 5xFAD combination, which falls below the additive prediction. Second, on the subtraction-only scale the genotype × age interaction is stable to the mean-centred community-count covariate (*P* = 0.008 with, *P* = 0.007 without; genotype × age-only model *P* = 0.002), so it does not depend on that nuisance term. Third, the small-world coefficient carries the same-sign interaction on both null-corrected scales: on the ratio scale *σ*_cfg_ (Age 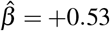; genotype × age 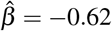) and on the *z* scale 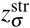 (Age 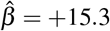; genotype × age 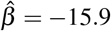), reaching significance on the ratio scale only. Novelty terms, including Age × novel and genotype × age × novel, were non-significant under the strength-preserving null (all *P >* 0.37 on the ratio scale and *P >* 0.97 on the *z* scale), so the apparent novelty-gating of *σ* visible in a preliminary topology-only (degree-only) null analysis is not retained when the degree and strength sequence of the backbone is held fixed.

Finally, the exploratory structure–function relationship of Fig. 5C was assessed with a Pearson correlation between session-level modularity (Δ*Q*^str^) and topology-dependent RSI (within − between) in the familiar environment, computed within each group. Young 5xFAD showed the strongest coupling (*r* = 0.88, *n* = 8, *P* = 0.004); the aged 5xFAD group was also nominally significant (*r* = 0.56, *n* = 13, *P* = 0.046), while the two wild-type groups were not (*r* = 0.18 and *r* = 0.22, both ns). *Neither the k-sweep (Supplementary Note, “robustness of the modularity deficit to the CkNN neighbourhood size”) nor this correlation was pre-registered; both are post-hoc. The correlation family consists of four tests, one per group, each relating* Δ*Q*^str^ *to topology-dependent RSI in the familiar environment. No family-wise error correction was applied. Bonferroni correction over these four tests (α* = 0.0125*) would retain the young-5xFAD coupling (P* = 0.004*) but not the aged-5xFAD coupling (P* = 0.046*). Within-group n is small, so the confidence intervals are wide. We therefore regard the result as hypothesis-generating and in need of replication in independent cohorts*.

**Supplementary Fig. 11.**
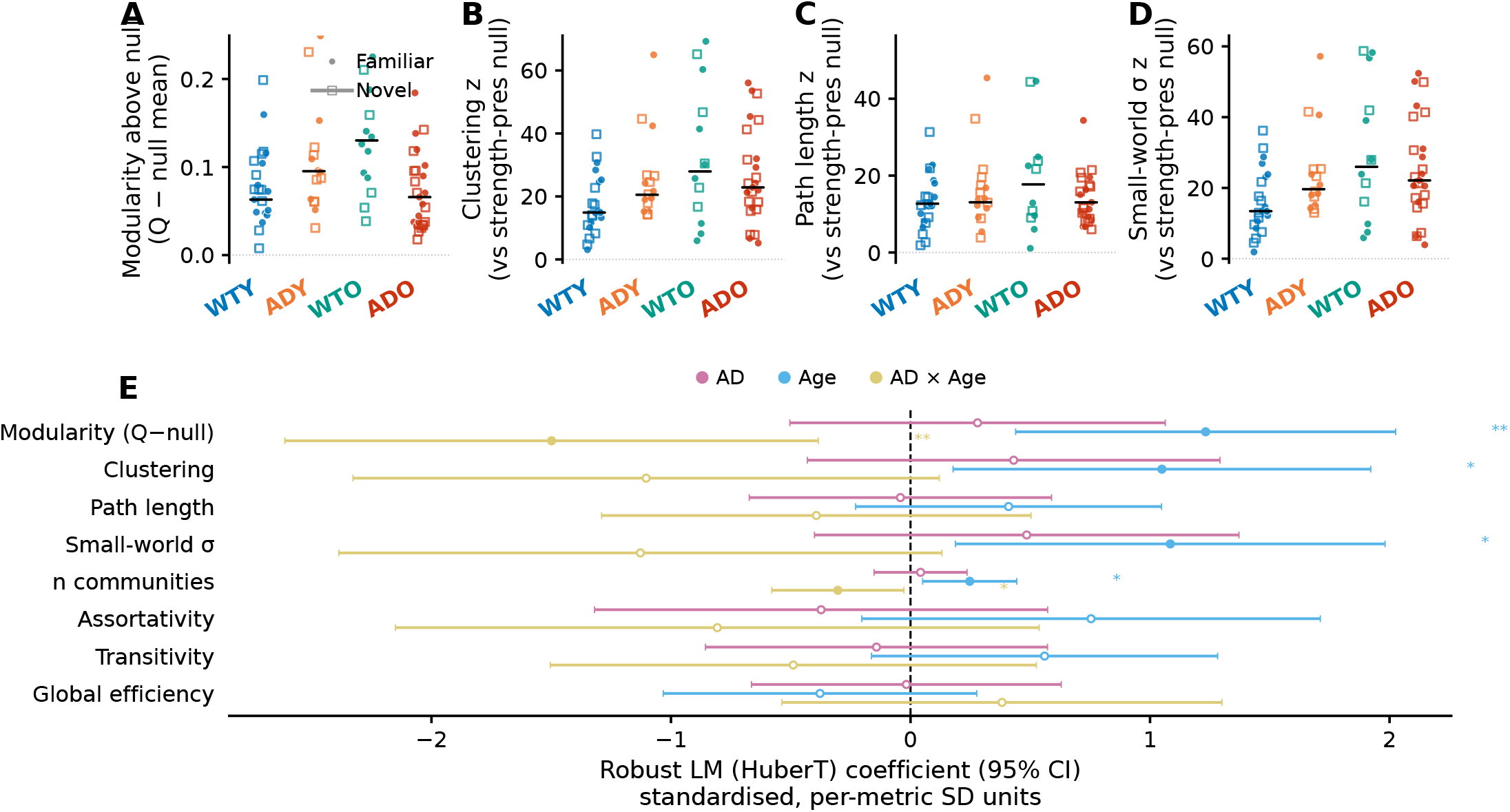
Full network structure under robust LM with community-count covariate (Huber M-estimator), strength-preserving configuration-model bias correction across all eight structural metrics. Full 8-metric structural analysis underlying Fig. 5. Each observed metric is benchmarked against a per-session strength-preserving configuration-model null (*N*_null_ = 500 rewires; Methods). Modularity (partition-dependent, Markov-stability) is bias-corrected by subtraction only (Δ*Q*^str^ = *Q*_obs_ − ⟨*Q*_null_⟩, as in main Fig. 5); the seven partition-free metrics use the per-session null mean and standard deviation to form a *z*-score. These null-corrected values are the dependent variables for the Huber LM. **A–D**, Per-session null-corrected values for the four metrics that carry Huber-model effects: modularity above null (A, Δ*Q*^str^), clustering coefficient (B, *z*), average shortest path length (C, *z*) and small-world coefficient *σ* (D, *z*), per group for familiar (filled circles) and novel (open squares) environments. The remaining four metrics (*n*_communities_, assortativity, transitivity, global efficiency) are near-redundant with these (efficiency ≡ 1*/*path, *ρ* = −0.99; *σ* ≡ clustering, *ρ* = 1.0; transitivity ≈ clustering, *ρ* = 0.85) and are retained only in the forest (E). After configuration-model correction no metric differs across groups at the univariate level (Kruskal–Wallis, familiar-only, all *P >* 0.05). **E**, Robust LM (Huber M-estimator) coefficients for the three interpretable predictors (genotype, age and their interaction) shown per metric (one row per metric) across all eight metrics (modularity, clustering, average path length, small-world *σ*, number of communities, assortativity, transitivity, global efficiency). To place the eight on one axis each dependent variable is standardised to unit variance within-metric before fitting. The full model additionally controls for novelty, mean-centred *n*_communities,c_ and all two- and three-way interactions (metric ~ (Genotype + Age + Novel + *n*_communities,c_)^3^); these covariate and nuisance terms are fitted but not plotted. Filled markers *P <* 0.05; open markers n.s.; horizontal bars 95% CI; stars ^*^*P <* 0.05, ^**^*P <* 0.01. The genotype × age deficit is carried by modularity (Δ*Q*^str^: genotype × age *P* = 0.008, Age *P* = 0.002, genotype main effect n.s. *P* = 0.48), with clustering and the small-world coefficient *σ* showing the same-sign genotype × age effect at trend level (*P* = 0.077 and *P* = 0.079) on significant Age main effects (*P* = 0.018 each); average path length carried no significant term under the Markov-stability community-count covariate (all *P >* 0.2). Assortativity, transitivity and global efficiency show no interpretable genotype × age term; the number-of-communities row’s terms track its own mean-centred covariate and are not interpreted. Reported *P*-values are nominal across an 8-metric × 7-predictor family of 56 tests. *N* = 77 sessions across 4 groups.

### Supplementary Note: robustness of the modularity deficit to the CkNN neighbourhood size

We assessed whether the genotype × age modularity deficit depends on the CkNN neighbourhood size *k* = 20 used throughout. For each *k* ∈ *{*10, 15, 20, 25, 30} we rebuilt the CkNN+MST backbone from the saved raw mutual-information matrices (identical construction to the *k* = 20 backbone; Methods), recomputed the Markov-stability modularity Δ*Q*^str^ = *Q*_obs_ − ⟨*Q*_null_⟩ against *N*_null_ strength-preserving nulls, and refitted the identical Huber model, reading off the genotype × age (AD:Age) interaction. Because Δ*Q*^str^ is a bounded modularity difference (not a *z*-score that inflates with graph density), the interaction is comparable across *k* without renormalisation and is plotted on its native scale (Supplementary Fig. 12C). The interaction is subadditive with a zero-excluding 95% CI at every *k* tested: 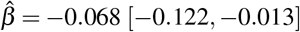 at *k* = 10, −0.075 [−0.134, −0.015] at *k* = 15, −0.077 [−0.139, −0.014] at *k* = 20, −0.098 [−0.165, −0.031] at *k* = 25 and −0.080 [−0.141, −0.019] at *k* = 30, so *k* = 20 is mid-range rather than a maximum. The sweep refits the model on backbones rebuilt from the raw mutual-information matrices, so its *k* = 20 estimate (−0.077) is a close but not identical reproduction of the main-analysis value quoted in the Results (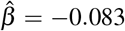, 95% CI [−0.144, −0.021]; Fig. 5). This aggregate-statistic robustness is the relevant test because community *membership* is itself moderately *k*-dependent (Supplementary Fig. 3D).

**Supplementary Fig. 12.**
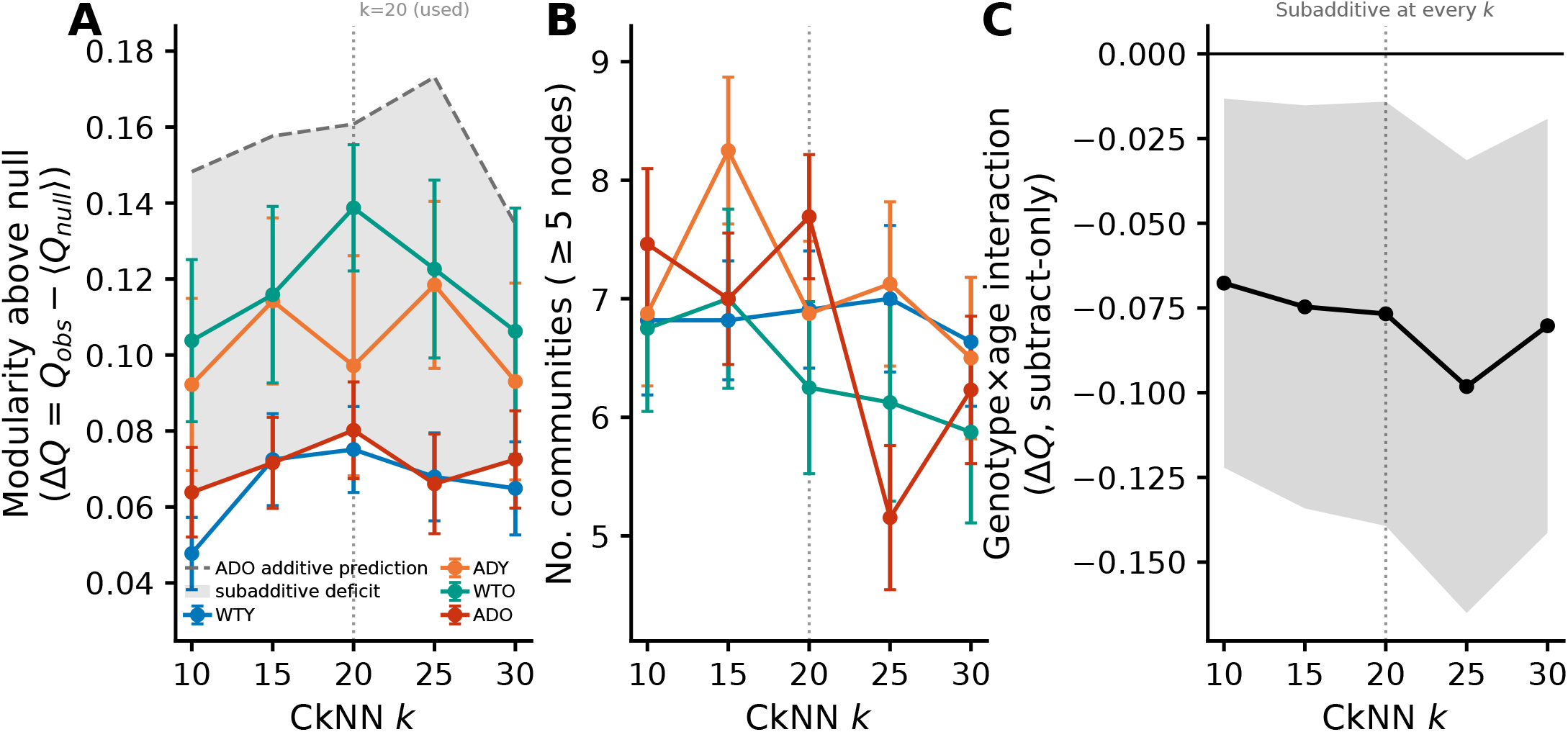
The genotype × age network deficit is robust to the CkNN graph-construction parameter *k*. The functional-network backbone is built with Continuous *k*-Nearest-Neighbours (CkNN) at *k* = 20 (Methods). To confirm that the network findings are not an artefact of this choice, the entire backbone construction and configuration-model analysis were repeated across *k* ∈*{*10, 15, 20, 25, 30}, rebuilding the CkNN+MST backbone from the saved raw mutual-information matrices and recomputing the Markov-stability modularity above null (Δ*Q*^str^ = *Q*_obs_ − ⟨*Q*_null_⟩, subtract-only) for every session. The value used in the paper (*k* = 20) is dotted in each panel. **A**, Per-group Δ*Q*^str^ (the Fig. 5 metric) versus *k* (familiar environment; mean ± s.e.m.); WTY is the lowest group at three of the five *k*, with ADO marginally below it at *k* = 15 and *k* = 25. ADO is the only double-hit (genotype *and* age) group, hence the only one in which an interaction can appear; its dashed line is the additive, no-interaction expectation ADO_pred_ = ADY+ WTO − WTY, where ADO would sit if amyloid and ageing acted independently. Observed ADO falls below it (shaded wedge): the subadditive genotype × age deficit. **B**, Number of communities (≥ 5 neurons) per session versus *k*, per group (mean ± s.e.m.); community count varies gently with *k* (~ 1–2 communities over *k* ∈ [10, 30]) with no abrupt change near *k* = 20. **C**, Genotype × age interaction (AD:Age; robust Huber M-estimator, same model as the main analysis) on Δ*Q*^str^ versus *k*, with 95% CI (shaded). Because Δ*Q*^str^ is a bounded modularity difference (not a *z*-score that inflates with graph density) the interaction is plotted on its native scale, with no scale-free renormalisation. The interaction is subadditive with a 95% CI excluding zero at every *k* tested, ranging from 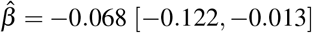 at *k* = 10 to −0.098 [−0.165, −0.031] at *k* = 25; at *k* = 20 it is −0.077 [−0.139, −0.014], mid-range rather than a maximum. This sweep rebuilds each backbone from the raw mutual-information matrices and refits the model, so its *k* = 20 estimate reproduces, without exactly matching, the main-analysis value reported in the Results (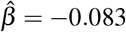, 95% CI [−0.144, −0.021]). The genotype × age modularity deficit is therefore not an artefact of the CkNN *k*. All 385 (session, *k*) cells contributed; each ⟨*Q*_null_ ⟩is the mean over 500 strength-preserving rewirings (median 500, minimum 382 usable after rewiring failures).

**Supplementary Fig. 13.**
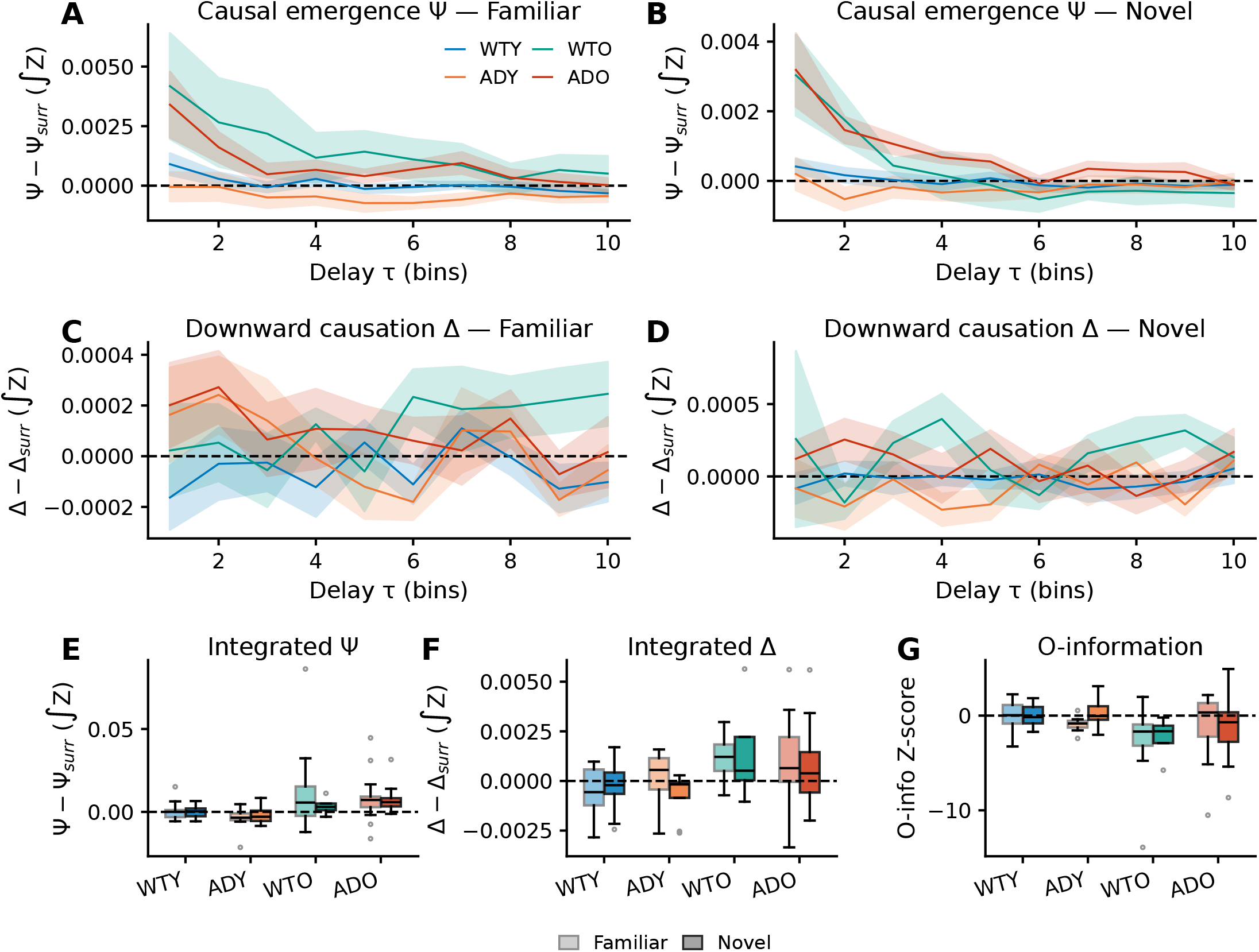
Community-level emergence: delay-resolved traces and per-group integrated scores. **A,B**, Per-delay surrogate-corrected causal emergence Ψ (observed − surrogate median; baseline-subtracted, *not* standardised by *σ*_shuf_, hence in units of mutual information; the “*Z*” suffix in the y-axis label is informal shorthand, *not* a *z*-score) across delays *τ* ∈ *{*1, …, 10}, group means ± s.e.m., for the familiar (A) and novel (B) environments. **C,D**, Per-delay downward causation Δ (observed − surrogate median, same baseline-subtracted convention) for familiar (C) and novel (D) environments. **E–G**, Per-session integrated scores per group × environment (familiar lighter, novel darker): integrated Ψ (E) and Δ (F) (sum over delays of [observed − surrogate median]); and O-information *z*-score (G, standardised by surrogate *σ*). Ψ/Δ retain mutual-information units while O-information is unit-free, so magnitudes are not directly comparable across measures (Methods). Within-group familiar-vs-novel differences are non-significant (permutation test, 10^4^ permutations), consistent with the non-significant Novel main effect for O-information in the robust LM (Fig. 6F). These panels complement the robust-LM coefficients in Fig. 6D–F by showing the underlying per-session distributions and group × environment structure. Boxplots: median, IQR, 1.5 × IQR whiskers; outliers as open circles. Definitions and surrogate construction follow Rosas et al.^57^ and Rajpal et al.^58^; *N* = 77 sessions across four groups (*N* = 75 for the O-information panel; two sessions have an undefined surrogate-normalised score).

## References

1. O’Keefe, J. & Dostrovsky, J. The hippocampus as a spatial map. preliminary evidence from unit activity in the freely-moving rat. Brain Res. 34, 171–175, DOI: 10.1016/0006-8993(71)90358-1 (1971).

2. Moser, E. I., Kropff, E. & Moser, M.-B. Place cells, grid cells, and the brain’s spatial representation system. Annu. Rev. Neurosci. 31, 69–89, DOI: 10.1146/annurev.neuro.31.061307.090723 (2008).

3. Wilson, M. A. & McNaughton, B. L. Dynamics of the hippocampal ensemble code for space. Science 261, 1055–1058, DOI: 10.1126/science.8351520 (1993).

4. Skaggs, W. E., McNaughton, B. L., Gothard, K. M. & Markus, E. J. An information-theoretic approach to deciphering the hippocampal code. In Advances in Neural Information Processing Systems, vol. 5, 1030–1037 (Morgan Kaufmann, 1993).

5. Panzeri, S., Treves, A., Schultz, S. R. & Rolls, E. T. On decoding the responses of a population of neurons from short time windows. Neural Comput. 11, 1553–1577, DOI: 10.1162/089976699300016142 (1999).

6. Dombeck, D. A., Harvey, C. D., Tian, L., Looger, L. L. & Tank, D. W. Functional imaging of hippocampal place cells at cellular resolution during virtual navigation. Nat. Neurosci. 13, 1433–1440, DOI: 10.1038/nn.2648 (2010).

7. Ziv, Y. et al. Long-term dynamics of CA1 hippocampal place codes. Nat. Neurosci. 16, 264–266, DOI: 10.1038/nn.3329 (2013).

8. Schultz, S. R. & Rolls, E. T. Analysis of information transmission in the Schaffer collaterals. Hippocampus 9, 582–598 (1999).

9. Schultz, S. R., Panzeri, S., Rolls, E. T. & Treves, A. Quantitative analysis of a Schaffer collateral model. In Baddeley, R., Hancock, P. & Földiák, P. (eds.) Information Theory and the Brain, 257–272, DOI: 10.1017/CBO9780511665516.019 (Cambridge University Press, Cambridge, UK, 2000).

10. Schultz, S. R. & Treves, A. Stability of the replica-symmetric solution for the information conveyed by a neural network. Phys. Rev. E 57, 3302–3310, DOI: 10.1103/PhysRevE.57.3302 (1998).

11. Holtmaat, A. & Caroni, P. Functional and structural underpinnings of neuronal assembly formation in learning. Nat. Neurosci. 19, 1553–1562, DOI: 10.1038/nn.4418 (2016).

12. Harris, K. D., Csicsvari, J., Hirase, H., Dragoi, G. & Buzsáki, G. Organization of cell assemblies in the hippocampus. Nature 424, 552–556, DOI: 10.1038/nature01834 (2003).

13. Buzsáki, G. Neural syntax: cell assemblies, synapsembles, and readers. Neuron 68, 362–385, DOI: 10.1016/j.neuron.2010.09.023 (2010).

14. Gava, G. P. et al. Integrating new memories into the hippocampal network activity space. Nat. Neurosci. 24, 326–330, DOI: 10.1038/s41593-021-00804-w (2021).

15. Maimon, S. R. et al. Sparse-to-dense coding transformation between hippocampal areas CA3 and CA1. Nature DOI: 10.1038/s41586-026-10537-0 (2026).

16. Bassett, D. S. & Sporns, O. Network neuroscience. Nat. Neurosci. 20, 353–364, DOI: 10.1038/nn.4502 (2017).

17. Curto, C. & Sanderson, N. Topological neuroscience: linking circuits to function. Annu. Rev. Neurosci. 48, 491–518, DOI: 10.1146/annurev-neuro-112723-034315 (2025).

18. Colgin, L. L., Moser, E. I. & Moser, M.-B. Understanding memory through hippocampal remapping. Trends Neurosci. 31, 469–477, DOI: 10.1016/j.tins.2008.06.008 (2008).

19. Leutgeb, S. et al. Independent codes for spatial and episodic memory in hippocampal neuronal ensembles. Science 309, 619–623, DOI: 10.1126/science.1114037 (2005).

20. Bakermans, J. J. W., Warren, J., Whittington, J. C. R. & Behrens, T. E. J. Constructing future behavior in the hippocampal formation through composition and replay. Nat. Neurosci. 28, 1061–1072, DOI: 10.1038/s41593-025-01908-3 (2025).

21. Go, M. A. et al. Amyloid pathology compresses dynamic range and degrades spatial coding in an Alzheimer’s mouse model. bioRxiv DOI: 10.1101/2025.06.27.661987 (2025).

22. Ying, J. et al. Disruption of the grid cell network in a mouse model of early Alzheimer’s disease. Nat. Commun. 13, 886, DOI: 10.1038/s41467-022-28551-x (2022).

23. Cacucci, F., Yi, M., Wills, T. J., Chapman, P. & O’Keefe, J. Place cell firing correlates with memory deficits and amyloid plaque burden in Tg2576 Alzheimer mouse model. Proc. Natl. Acad. Sci. 105, 7863–7868, DOI: 10.1073/pnas.0802908105 (2008).

24. Zhang, H. et al. Degenerate mapping of environmental location presages deficits in object-location encoding and memory in the 5xFAD mouse model for Alzheimer’s disease. Neurobiol. Dis. 176, 105939, DOI: 10.1016/j.nbd.2022.105939 (2023).

25. Mably, A. J., Gereke, B. J., Jones, D. T. & Colgin, L. L. Impairments in spatial representations and rhythmic coordination of place cells in the 3xTg mouse model of Alzheimer’s disease. Hippocampus 27, 378–392, DOI: 10.1002/hipo.22697 (2017).

26. Jun, H. et al. Disrupted place cell remapping and impaired grid cells in a knockin model of Alzheimer’s disease. Neuron 107, 1095–1112.e6, DOI: 10.1016/j.neuron.2020.06.023 (2020).

27. Prince, S. M. et al. Alzheimer’s pathology causes impaired inhibitory connections and reactivation of spatial codes during spatial navigation. Cell Reports 35, 109008, DOI: 10.1016/j.celrep.2021.109008 (2021).

28. Busche, M. A. et al. Clusters of hyperactive neurons near amyloid plaques in a mouse model of Alzheimer’s disease. Science 321, 1686–1689, DOI: 10.1126/science.1162844 (2008).

29. Barnes, C. A., Suster, M. S., Shen, J. & McNaughton, B. L. Multistability of cognitive maps in the hippocampus of old rats. Nature 388, 272–275, DOI: 10.1038/40859 (1997).

30. Wilson, I. A., Ikonen, S., Gallagher, M., Eichenbaum, H. & Tanila, H. Age-associated alterations of hippocampal place cells are subregion specific. J. Neurosci. 25, 6877–6886, DOI: 10.1523/JNEUROSCI.1744-05.2005 (2005).

31. Rajpal, H. et al. Synergy mediates long-range correlations in the visual cortex near criticality. Front. Comput. Neurosci. 20, 1741793, DOI: 10.3389/fncom.2026.1741793 (2026).

32. Ah-Weng, R. & Rajpal, H. Collective dynamics in spiking neural networks beyond Dale’s principle. arXiv preprint arXiv:2602.23202 DOI: 10.48550/arXiv.2602.23202 (2026).

33. Sherrill, S. P., Timme, N. M., Beggs, J. M. & Newman, E. L. Partial information decomposition reveals that synergistic neural integration is greater downstream of recurrent information flow in organotypic cortical cultures. PLoS Comput. Biol. 17, e1009196, DOI: 10.1371/journal.pcbi.1009196 (2021).

34. Koçillari, L. et al. Behavioural relevance of redundant and synergistic stimulus information between functionally connected neurons in mouse auditory cortex. Brain Informatics 10, 34, DOI: 10.1186/s40708-023-00212-9 (2023).

35. Rosas, F. E., Mediano, P. A. M., Gastpar, M. & Jensen, H. J. Quantifying high-order interdependencies via multivariate extensions of the mutual information. Phys. Rev. E 100, 032305, DOI: 10.1103/PhysRevE.100.032305 (2019).

36. Luppi, A. I. et al. A synergistic core for human brain evolution and cognition. Nat. Neurosci. 25, 771–782, DOI: 10.1038/s41593-022-01070-0 (2022).

37. Varley, T. F., Pope, M., Faskowitz, J. & Sporns, O. Multivariate information theory uncovers synergistic subsystems of the human cerebral cortex. Commun. Biol. 6, 451, DOI: 10.1038/s42003-023-04843-w (2023).

38. Newman, E. L., Varley, T. F., Parakkattu, V. K., Sherrill, S. P. & Beggs, J. M. Revealing the dynamics of neural information processing with multivariate information decomposition. Entropy 24, 930, DOI: 10.3390/e24070930 (2022).

39. Panzeri, S., Schultz, S. R., Treves, A. & Rolls, E. T. Correlations and the encoding of information in the nervous system. Proc. Royal Soc. B: Biol. Sci. 266, 1001–1012, DOI: 10.1098/rspb.1999.0736 (1999).

40. Schultz, S. R. & Panzeri, S. Temporal correlations and neural spike train entropy. Phys. Rev. Lett. 86, 5823–5826, DOI: 10.1103/PhysRevLett.86.5823 (2001).

41. Schultz, S. R., Panzeri, S., Treves, A. & Rolls, E. T. Correlated firing and the information represented by neurons in short epochs. Neurocomputing 26–27, 499–504, DOI: 10.1016/S0925-2312(99)00040-5 (1999).

42. Schneidman, E., Bialek, W. & Berry, M. J. Synergy, redundancy, and independence in population codes. J. Neurosci. 23, 11539–11553, DOI: 10.1523/JNEUROSCI.23-37-11539.2003 (2003).

43. Latham, P. E. & Nirenberg, S. Synergy, redundancy, and independence in population codes, revisited. J. Neurosci. 25, 5195–5206, DOI: 10.1523/JNEUROSCI.5319-04.2005 (2005).

44. Panzeri, S. & Schultz, S. R. A unified approach to the study of temporal, correlational, and rate coding. Neural Comput. 13, 1311–1349, DOI: 10.1162/08997660152002870 (2001).

45. Panzeri, S., Petersen, R. S., Schultz, S. R., Lebedev, M. & Diamond, M. E. The role of spike timing in the coding of stimulus location in rat somatosensory cortex. Neuron 29, 769–777, DOI: 10.1016/S0896-6273(01)00251-3 (2001).

46. Williams, P. L. & Beer, R. D. Nonnegative decomposition of multivariate information. arXiv preprint arXiv:1004.2515 DOI: 10.48550/arXiv.1004.2515 (2010).

47. Luppi, A. I., Rosas, F. E., Mediano, P. A. M., Menon, D. K. & Stamatakis, E. A. Information decomposition and the informational architecture of the brain. Trends Cogn. Sci. 28, 352–368, DOI: 10.1016/j.tics.2023.11.005 (2024).

48. Barlow, H. B. Possible principles underlying the transformations of sensory messages. In Rosenblith, W. A. (ed.) Sensory Communication, 217–234 (MIT Press, Cambridge, MA, 1961).

49. Barlow, H. Redundancy reduction revisited. Network: Comput. Neural Syst. 12, 241–253, DOI: 10.1080/net.12.3.241.253 (2001).

50. Gava, G. P. et al. Organizing the coactivity structure of the hippocampus from robust to flexible memory. Science 385, 1120–1127, DOI: 10.1126/science.adk9611 (2024).

51. Barrett, A. B. Exploration of synergistic and redundant information sharing in static and dynamical Gaussian systems. Phys. Rev. E 91, 052802, DOI: 10.1103/PhysRevE.91.052802 (2015).

52. Delvenne, J.-C., Yaliraki, S. N. & Barahona, M. Stability of graph communities across time scales. Proc. Natl. Acad. Sci. 107, 12755–12760, DOI: 10.1073/pnas.0903215107 (2010).

53. Lambiotte, R.Delvenne, J.-C. & Barahona, M. Random walks, Markov processes and the multiscale modular organization of complex networks. IEEE Transactions on Netw. Sci. Eng. 1, 76–90, DOI: 10.1109/TNSE.2015.2391998 (2014).

54. Down, K. J. A., Huntley, J., Mediano, P. A. M. & Bor, D. Synergistic and redundant information dynamics are modulated by Alzheimer’s disease and cognitive impairment. bioRxiv DOI: 10.64898/2026.02.18.706630 (2026). Preprint.

55. Bertschinger, N., Rauh, J., Olbrich, E., Jost, J. & Ay, N. Quantifying unique information. Entropy 16, 2161–2183, DOI: 10.3390/e16042161 (2014).

56. Makkeh, A., Theis, D. O. & Vicente, R. BROJA-2PID: a robust estimator for bivariate partial information decomposition. Entropy 20, 271, DOI: 10.3390/e20040271 (2018).

57. Rosas, F. E. et al. Reconciling emergences: An information-theoretic approach to identify causal emergence in multivariate data. PLoS Comput. Biol. 16, e1008289, DOI: 10.1371/journal.pcbi.1008289 (2020).

58. Rajpal, H., Mediano, P. A., Sas, M. I., Jensen, H. J. & Rosas, F. E. Quantifying the emergence of population-level activity in neuronal systems. bioRxiv DOI: 10.64898/2026.02.13.705719 (2026). https://www.biorxiv.org/content/early/2026/02/16/2026.02.13.705719.full.pdf.

59. Timme, N. M. et al. High-degree neurons feed cortical computations. PLOS Comput. Biol. 12, e1004858, DOI: 10.1371/journal.pcbi.1004858 (2016).

60. Sherrill, S. P., Timme, N. M., Beggs, J. M. & Newman, E. L. Correlated activity favors synergistic processing in local cortical networks in vitro at synaptically relevant timescales. Netw. Neurosci. 4, 678–697, DOI: 10.1162/netn_a_00141 (2020).

61. Zhu, X., Raina, A. K., Perry, G. & Smith, M. A. Alzheimer’s disease: the two-hit hypothesis. The Lancet Neurol. 3, 219–226, DOI: 10.1016/S1474-4422(04)00707-0 (2004).

62. Palop, J. J. & Mucke, L. Network abnormalities and interneuron dysfunction in Alzheimer disease. Nat. Rev. Neurosci. 17, 777–792, DOI: 10.1038/nrn.2016.141 (2016).

63. Herrup, K. Reimagining Alzheimer’s disease: an age-based hypothesis. J. Neurosci. 30, 16755–16762, DOI: 10.1523/JNEUROSCI.4521-10.2010 (2010).

64. Li, Y. et al. Early spatial and contextual coding deficits in hippocampal CA1 precede performance decline in an Alzheimer’s disease model. bioRxiv DOI: 10.1101/2025.02.05.636661 (2026). Preprint.

65. Beggs, J. M. & Plenz, D. Neuronal avalanches in neocortical circuits. J. Neurosci. 23, 11167–11177, DOI: 10.1523/JNEUROSCI.23-35-11167.2003 (2003).

66. Rajpal, H. & Goodman, D. Emergent generalization by representation learning in artificial neural networks. arXiv preprint arXiv:2607.10430 (2026).

67. Tononi, G., Sporns, O. & Edelman, G. M. A measure for brain complexity: relating functional segregation and integration in the nervous system. Proc. Natl. Acad. Sci. 91, 5033–5037, DOI: 10.1073/pnas.91.11.5033 (1994).

68. Saito, T. et al. Single app knock-in mouse models of alzheimer’s disease. Nat. neuroscience 17, 661–663 (2014).

69. Foley, A. M., Ammar, Z. M., Lee, R. H. & Mitchell, C. S. Systematic review of the relationship between amyloid-β levels and measures of transgenic mouse cognitive deficit in Alzheimer’s disease. J. Alzheimer’s Dis. 44, 787–795, DOI: 10.3233/JAD-142208 (2015).

70. Sasaguri, H. et al. APP mouse models for Alzheimer’s disease preclinical studies. The EMBO J. 36, 2473–2487, DOI: 10.15252/embj.201797397 (2017).

71. Fortunato, S. & Barthélemy, M. Resolution limit in community detection. Proc. Natl. Acad. Sci. 104, 36–41, DOI: 10.1073/pnas.0605965104 (2007).

72. Good, B. H., de Montjoye, Y.-A. & Clauset, A. Performance of modularity maximization in practical contexts. Phys. Rev. E 81, 046106, DOI: 10.1103/PhysRevE.81.046106 (2010).

73. Papo, D., Zanin, M., Martínez, J. H. & Buldú, J. M. Beware of the Small-World Neuroscientist! Front. Hum. Neurosci. 10, 96, DOI: 10.3389/fnhum.2016.00096 (2016).

74. Lopes-dos Santos, V., Ribeiro, S. & Tort, A. B. L. Detecting cell assemblies in large neuronal populations. J. Neurosci. Methods 220, 149–166, DOI: 10.1016/j.jneumeth.2013.04.010 (2013).

75. Oakley, H. et al. Intraneuronal β-amyloid aggregates, neurodegeneration, and neuron loss in transgenic mice with five familial Alzheimer’s disease mutations: potential factors in amyloid plaque formation. J. Neurosci. 26, 10129–10140, DOI: 10.1523/JNEUROSCI.1202-06.2006 (2006).

76. Dana, H. et al. High-performance calcium sensors for imaging activity in neuronal populations and microcompartments. Nat. Methods 16, 649–657, DOI: 10.1038/s41592-019-0435-6 (2019).

77. Chen, T.-W. et al. Ultrasensitive fluorescent proteins for imaging neuronal activity. Nature 499, 295–300, DOI: 10.1038/nature12354 (2013).

78. Friedrich, J., Zhou, P. & Paninski, L. Fast online deconvolution of calcium imaging data. PLOS Comput. Biol. 13, e1005423, DOI: 10.1371/journal.pcbi.1005423 (2017).

79. Giovannucci, A. et al. CaImAn an open source tool for scalable calcium imaging data analysis. eLife 8, e38173, DOI: 10.7554/eLife.38173 (2019).

80. Strong, S. P., Koberle, R., de Ruyter van Steveninck, R. R. & Bialek, W. Entropy and information in neural spike trains. Phys. Rev. Lett. 80, 197–200, DOI: 10.1103/PhysRevLett.80.197 (1998).

81. Wolpert, D. H. & Wolf, D. R. Estimating functions of probability distributions from a finite set of samples. Phys. Rev. E 52, 6841–6854, DOI: 10.1103/PhysRevE.52.6841 (1995).

82. Nemenman, I., Shafee, F. & Bialek, W. Entropy and inference, revisited. In Advances in neural information processing systems, vol. 14, 471–478 (2002).

83. Paninski, L. Estimation of entropy and mutual information. Neural computation 15, 1191–1253, DOI: 10.1162/089976603321780272 (2003).

84. Nemenman, I., Bialek, W. & de Ruyter van Steveninck, R. Entropy and information in neural spike trains: Progress on the sampling problem. Phys. Rev. E 69, 056111, DOI: 10.1103/PhysRevE.69.056111 (2004).

85. Nemenman, I. Coincidences and estimation of entropies of random variables with large cardinalities. Entropy 13, 2013–2023, DOI: 10.3390/e13122013 (2011).

86. Archer, E., Park, I. M. & Pillow, J. W. Bayesian and quasi-Bayesian estimators for mutual information from discrete data. Entropy 15, 1738–1755, DOI: 10.3390/e15051738 (2013).

87. Archer, E., Park, I. M. & Pillow, J. W. Bayesian entropy estimation for countable discrete distributions. The J. Mach. Learn. Res. 15, 2833–2868 (2014).

88. Panzeri, S., Senatore, R., Montemurro, M. A. & Petersen, R. S. Correcting for the sampling bias problem in spike train information measures. J. Neurophysiol. 98, 1064–1072, DOI: 10.1152/jn.00559.2007 (2007).

89. Marsili, S. ndd: Bayesian entropy estimation from discrete data. https://github.com/simomarsili/ndd (2016).

90. Pola, G., Schultz, S. R., Petersen, R. S. & Panzeri, S. A practical guide to information analysis of spike trains. In Kötter, R. (ed.) Neuroscience Databases: A Practical Guide, 139–154, DOI: 10.1007/978-1-4615-1079-6_10 (Kluwer Academic Publishers, Boston, MA, 2003).

91. Quian Quiroga, R. & Panzeri, S. Extracting information from neuronal populations: information theory and decoding approaches. Nat. Rev. Neurosci. 10, 173–185, DOI: 10.1038/nrn2578 (2009).

92. Timme, N. M. & Lapish, C. A tutorial for information theory in neuroscience. eNeuro 5, DOI: 10.1523/ENEURO.0052-18.2018 (2018).

93. Berry, T. & Sauer, T. Consistent manifold representation for topological data analysis. Foundations Data Sci. 1, 1–38, DOI: 10.3934/fods.2019001 (2019).

94. Arnaudon, A. et al. Algorithm 1044: PyGenStability, a multiscale community detection framework with generalized Markov stability. ACM Transactions on Math. Softw. 50, 1–8, DOI: 10.1145/3651225 (2024).

95. Blondel, V. D., Guillaume, J.-L., Lambiotte, R. & Lefebvre, E. Fast unfolding of communities in large networks. J. Stat. Mech. Theory Exp. 2008, P10008, DOI: 10.1088/1742-5468/2008/10/P10008 (2008).

96. Traag, V. A., Waltman, L. & van Eck, N. J. From Louvain to Leiden: guaranteeing well-connected communities. Sci. Reports 9, 5233, DOI: 10.1038/s41598-019-41695-z (2019).

97. Meilă, M. Comparing clusterings—an information based distance. J. Multivar. Analysis 98, 873–895, DOI: 10.1016/j.jmva.2006.11.013 (2007).

98. Vinh, N. X., Epps, J. & Bailey, J. Information theoretic measures for clusterings comparison: Variants, properties, normalization and correction for chance. J. machine learning research 11, 2837–2854 (2010).

99. Aarts, E., Verhage, M., Veenvliet, J. V., Dolan, C. V. & van der Sluis, S. A solution to dependency: using multilevel analysis to accommodate nested data. Nat. Neurosci. 17, 491–496, DOI: 10.1038/nn.3648 (2014).

100. Saravanan, V., Berman, G. J. & Sober, S. J. Application of the hierarchical bootstrap to multi-level data in neuroscience. Neurons, Behav. Data analysis, Theory 3, 1–25 (2020).

101. Humphries, M. D. & Gurney, K. Network ‘Small-World-Ness’: a quantitative method for determining canonical network equivalence. PLoS ONE 3, e0002051, DOI: 10.1371/journal.pone.0002051 (2008).

102. Newman, M. E. J. & Girvan, M. Finding and evaluating community structure in networks. Phys. Rev. E 69, 026113, DOI: 10.1103/PhysRevE.69.026113 (2004).

103. Maslov, S. & Sneppen, K. Specificity and stability in topology of protein networks. Science 296, 910–913, DOI: 10.1126/science.1065103 (2002).

104. Rubinov, M. & Sporns, O. Weight-conserving characterization of complex functional brain networks. NeuroImage 56, 2068–2079, DOI: 10.1016/j.neuroimage.2011.03.069 (2011).

105. Sas, M. I. et al. Improved estimators of causal emergence for large systems. arXiv preprint arXiv:2601.00013 DOI: 10.48550/arXiv.2601.00013 (2026).

